# RepGene: Auditing Condition-Specific Fusion Biases for Multi-View Gene Representations

**DOI:** 10.64898/2026.06.11.731512

**Authors:** Haiyang Hou, Tianyi Xia, Luni Hu, Yong Zhang, Yuxiang Li, Shuangsang Fang, Lei Cao

**Author notes:** Corresponding author: {, }.

## Abstract

This study evaluates how multi-view gene representations should be compared under a matched contract. Each of 18,460 genes is described by five aligned views: DNA sequence-derived features (D), RNA sequence-derived features (R), protein language-model embeddings (P), curated text knowledge (K), and single-cell expression (X). The contract fixes the gene universe, the 256-D output, label-free outer-training-only fitting, one random-forest probe, and task-equal Macro-F1 over nine registered binary tasks, and varies only the fusion operator across four protocol-defined input states: Core (DNA+RNA+protein), Core+K, Core+X, and Core+K+X. Under these states, the top-scoring operator changes with the available view combination and all task-bootstrap intervals overlap, establishing a condition-dependent rather than universal ranking pattern. An independently configured external RF track provides a separate check of the numerical range, while a gene-to-cell boundary diagnostic shows that gene-level differences attenuate toward a dimension-matched random control under expression-weighted aggregation, as predicted by a dilution identity. Historical log-derived multiplicative-interaction screening is retained only as magnitude-only supplementary evidence because fold-level records are unavailable. The contribution is therefore the matched contract and its audit trail: an evidence-graded protocol for turning fusion claims into comparable, condition-bounded measurements.

## 1 Introduction

A single gene is described by multiple biological views that are related but not interchangeable: sequence-derived DNA (D) features, sequence-derived RNA (R) features, protein language-model embeddings (P), curated text knowledge (K), and single-cell expression embeddings (X). These views differ in dimensionality, scale, coverage, and statistical geometry, and prior work reports task-dependent rather than uniformly dominant views (Du et al., 2019; Brechtmann et al., 2023; Zhong et al., 2025). The open question is downstream and consequential for every model that consumes such representations: given aligned view vectors for the same gene, which *fusion bias* should combine them, and how stable is that choice when the available views change?

DNA, RNA, and protein form the *Core*: the three sequence-bearing molecular layers of the central dogma are generally jointly available for an annotated gene, whereas K and X depend on curation or assay coverage. We retain all five views—each has distinct biological content and an established pretrained basis (Brixi et al., 2026; Chen et al., 2022; He et al., 2024; Rives et al., 2021; Elnaggar et al., 2022; Cui et al., 2024; Chen & Zou, 2025)—because cross-view differences matter more than within-view encoder choice (Brechtmann et al., 2023) (low cross-view CKA and neighbour overlap, Appendix 26). We therefore fix one sufficiently large encoder per view to isolate fusion, without claiming within-view equivalence.

That question is usually answered badly, because multiview comparisons change the input set, preprocessing, output dimension, fusion operator, probe, and split at the same time, so a score difference cannot be attributed to the fusion operator at all. The protocol therefore defines a *matched contract*: the gene universe, output budget, fold construction, label-free fitting boundary, probe, and aggregation rule are fixed, while only the fusion operator varies. The primary comparison covers six operators—concatenation and projection baselines, explicit pairwise products, a low-rank higher-order residual, and input-conditioned view weighting—with two attention operators analysed separately. The four conditions are controlled protocol states on a shared universe, not natural per-gene coverage.

The study answers four questions, introduced by Figure 1 and addressed in Results in order: which aligned views enter the contract, and whether K and X produce the same task response; how the preferred fusion bias varies across the four registered input states; whether the condition-dependent pattern survives an independently configured external probe; and where gene-level utility stops transferring to cells after expression-weighted aggregation. The last is explicitly a boundary diagnostic, not a universal cell-level claim. Because the paper reports evidence of different strengths, every result carries one of four evidence grades (Appendix 1).

**Figure 1:**
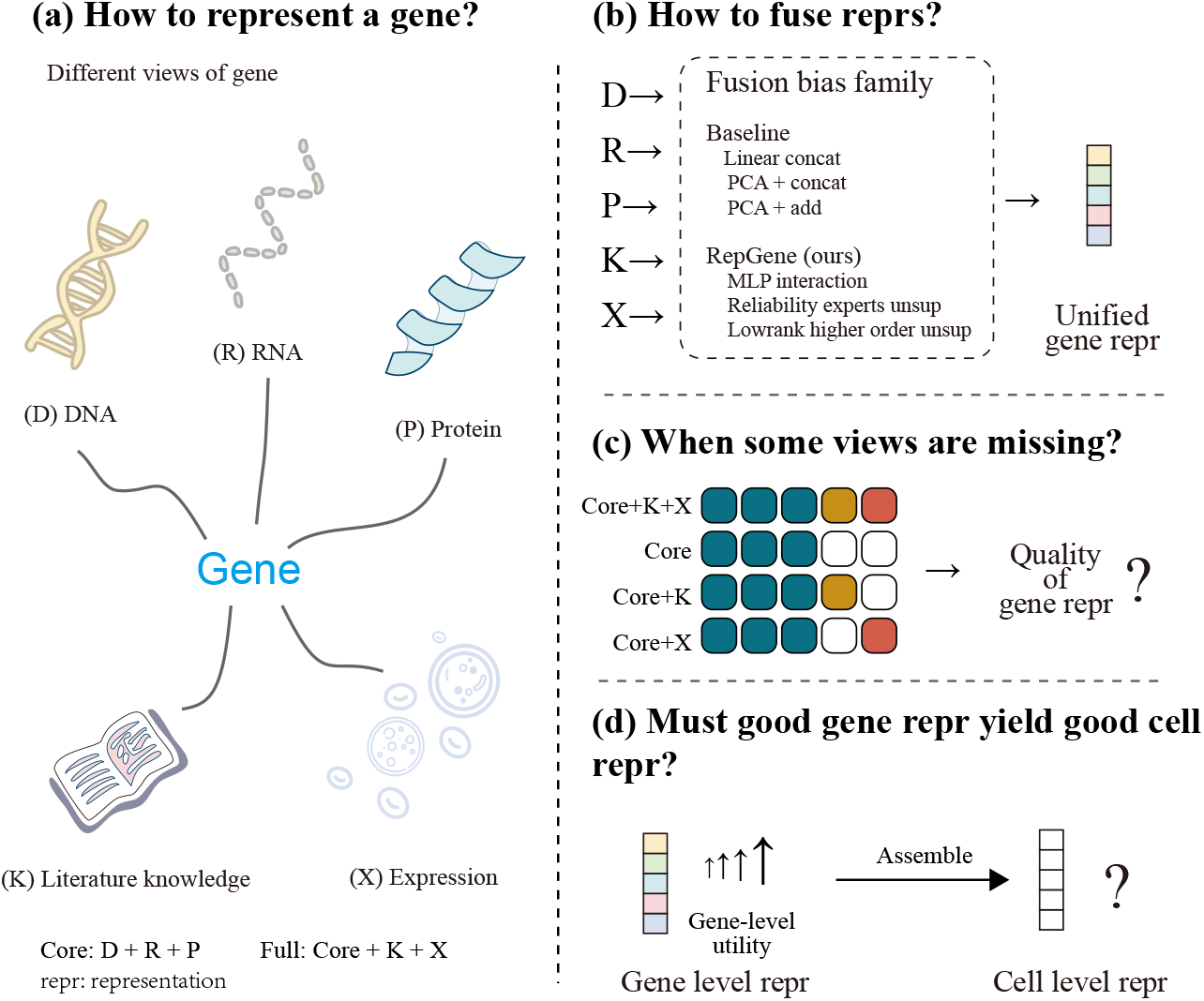
Four-question evaluation map for RepGene and condition-specific fusion selection. (a) Five aligned views per gene. (b) Six primary operators compared under a shared contract. (c) The internal study tests whether the preferred fusion changes across the four protocol states; all four states are controlled ablations on the fully aligned 18,460-gene universe, not natural missingness, imputation, or deployment robustness. (d) The gene-to-cell transfer question is a *boundary diagnostic* (out of the validated scope), not a completed fourth question. Conceptual map of study scope; not an empirical result.

Results follows the same order, and all intervals are descriptive task-bootstrap summaries over the fixed register, not significance tests. The attention operators, multiplicative-interaction sweep, and routing criteria remain appendix analyses and are not silently promoted to additional primary questions (Appendices 11, 24, 25).

We claim no new fusion architecture: the deliverable is the contract, a register-bounded measurement of the condition response, and the audit trail that makes every claim re-checkable from the released configurations. Three contributions organise the paper.

- **A matched fusion-bias contract as the primary method contribution**. A new fusion operator enters by exporting a 256-dimensional representation for the same gene universe and re-running one frozen probe; nothing else may be re-tuned. This converts fusion claims from leaderboard deltas into protocol-bounded comparisons (Section 3; protocol card in Appendix 19).
- **A register-bounded, condition-dependent picture with task-level heterogeneity preserved**. The largest task-equal mean changes with the input condition, all intervals overlap, and per-task winners switch in thirteen of fifteen tasks—stably across the binary sub-register, the full register, and a semantic-leakage control. Reading only the full-input column reverses the conclusion supported by the matched condition contrasts (Section 4).
- **Three boundary analyses that are findings in their own right**. (i) A granularity argument for why token-wise attention does not lead under a one-vector-per-view contract; (ii) an aggregation-dilution bound that predicts, before measurement, the attenuation of gene-level advantages at cell level; and (iii) a historical sweep bounding explicit multiplicative interaction below its capacity-matched ceiling (Sections 4.5, 4.4; Appendices 11, 24).

The primary research question is thereby narrowed to: *under a fixed contract, does the preferred fusion bias change with the protocol-defined input condition, and how large is the change relative to task heterogeneity?* (Dynamic routing is a separate evaluation target, Appendix 25.)

## 2 Related Work

### Gene representations across biological views

Gene representations have been built from co-expression, sequence, protein, interaction, and literature signals, with utility shown to depend on the downstream task rather than on a uniformly best modality (Du et al., 2019; Brechtmann et al., 2023). Established view resources include ProtVec, GeneMANIA, HumanNet, STRING (Asgari & Mofrad, 2015; Mostafavi et al., 2008; Hwang et al., 2019; Szklarczyk et al., 2023), and text-derived embeddings such as GenePT (Chen & Zou, 2025). Foundation models supply strong per-view encoders— for proteins (Rives et al., 2021; Elnaggar et al., 2022; Jumper et al., 2021), single cells (Cui et al., 2024; Yang et al., 2022; Theodoris et al., 2023), and integration workflows (Wolf et al., 2018; Lopez et al., 2018; Korsunsky et al., 2019; Hie et al., 2019; Chen et al., 2015; Welch et al., 2019b). We do not replace these encoders: their aligned outputs are our input views, and we isolate the downstream fusion question. The choice is deliberate: each sequence or cell view has a pretrained basis, while K contributes curated functional context. Multi-omics factor and generative models (MOFA, MOFA+, totalVI, MultiVI) address joint structure under a different estimand (Argelaguet et al., 2018; 2020; Gayoso et al., 2022; Ashuach et al., 2023; Stoeckius et al., 2017).

### Multiview fusion, attention, and where this setting differs

Attention is the default fusion choice in much of multimodal learning (Vaswani et al., 2017; Tsai et al., 2019; Baltrusaitis et al., 2018), where the fused model is trained end-to-end with task labels. The classical tensor-fusion family is the closest relative of our interaction experts: the Tensor Fusion Network (Zadeh et al., 2017) models unimodal, bimodal, and trimodal interactions through an explicit outer product, and Low-rank Multimodal Fusion (Liu et al., 2018) factorizes the same high-order interaction tensor; our Interaction MLP and Low-rank higher-order residual are label-free, one-vector-per-view analogues evaluated under a matched contract, not end-to-end supervision. Classical multiview statistics offers a second, older family of label-free fusion rules: canonical correlation analysis and its multi-set generalisation (Hotelling, 1936; Kettenring, 1971) and partial-least-squares variants (Wold, 1975) satisfy the contract’s entry conditions (a 256-D export plus the frozen probe), so they are legal entrants and the cheapest next additions to the register; the current package does not score them. Among molecular integration platforms, PRISME (Zheng et al., 2025) integrates heterogeneous molecular embeddings with an autoencoder trained under its own reconstruction objective; it enters this study only as an external-track protocol extension (Section 4.3), because the primary contract requires every operator to export 256-D representations fitted under the shared label-free objective. This setting differs twice from the supervised fusion literature: fitting is label-free, and the fused representation is consumed by a fixed downstream probe. Whether attention’s supervised advantages survive here is empirical; they do not lead in the reported Core+K+X comparison (Appendix 11), and we analyse why. The missing-modality literature (Wang et al., 2023; Wu et al., 2026; Chandrashekar et al., 2023; Khodaee et al., 2025) studies natural availability—imputation and deployment robustness—whereas our conditions are registered protocol states answering a different estimand; missing-modality operators remain legal contract entrants. Concurrent audit-scale benchmarks report the same phenomenon at the model level—rankings shifting with modality, preprocessing, and metric (Chen et al., 2026)— whereas we vary only the fusion operator at a fixed contract. Leakage-aware benchmarking motivates our outer-training-only fitting boundary (Luecken & Theis, 2019; Tran et al., 2020; Soneson & Robinson, 2018; Mereu et al., 2020; Hafemeister & Satija, 2019); Appendix 2 discusses the task-set accounting in detail.

## 3 Approach: A Matched Fusion-Bias Contract

The approach fixes a common representation interface and varies only the fusion map. The three learned experts share the trained per-view adapters, the 256-D output budget, the folds {3} and seeds {41, 42, 43}, the label-free objective, and one probe; the three closed-form baselines share the same outer-training-only standardization, budget, folds, seeds, and probe, but bypass the trained adapters and apply label-free PCA/linear maps to the standardized raw view vectors 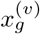 directly (Appendix 5). The resolved configuration (hidden width *h* = 96, dropout 0.1, eight epochs, batch size 1,024, Adam) and the label-free weights (*λ*_var_, *λ*_cov_, *λ*_align_, *λ*_avail_) = (0.1, 0.01, 0.05, 0.1) on a unit-weight reconstruction term are given in Appendix 6. The internal RF probe uses 300 trees, depth 16, minimum leaf size 2 (Breiman, 2001); the external track uses a separately configured probe (Appendix 12).

### Notation and contract

Let *g* ∈ *G* index a gene in the aligned universe (|*G*| = 18,460) and let *V* = {*D, R, P, K, X*} denote DNA, RNA, protein, knowledge (text), and single-cell views. A condition *c* selects *V*_*c*_ ∈ {*V*_Core_, *V*_Core+K_, *V*_Core+X_, *V*_Core+K+X_} with *V*_Core_ = {*D, R, P*}, *V*_Core+K_ = {*D, R, P, K*}, *V*_Core+X_ = {*D, R, P, X*}, and *V*_Core+K+X_ = *V*. Each view is adapted to the common width by

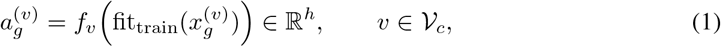

where every standardization and reduction inside fit_train_ is estimated on outer-training features only, and no downstream label is visible during representation fitting (Eq. 5). Every operator exports 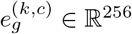 under the same budget.

### Operators and their biases

Six operators are compared under every condition, and two attention operators under Core+K+X (Appendix 11). Table 8 (Appendix 5) summarises the biases and the learned-parameter accounting. The three baselines carry no learned parameters: their PCA/linear maps are fitted label-free on outer-training features and operate on the fold-standardized raw views 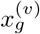, not on the adapter outputs 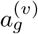. Linear concat projects the concatenated standardized views to 256 dimensions by PCA; PCA add reduces each view to 256 dimensions and averages across views; PCA then concat splits the 256-D budget across views before concatenation (per-view targets 86/85/85 under Core, 64 under the four-view conditions, 52/51/51/51/51 under Core+K+X). Because baselines and experts differ both in fusion bias *and* in whether learnable interaction terms exist, we record differences between function classes under a shared budget rather than decomposing them into “bias versus capacity” (Section 4.5).

### Pairwise interaction expert

The first expert exposes non-additive relations without a full high-order tensor. With the view stack 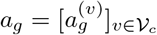, *F*_1_ a two-layer MLP:

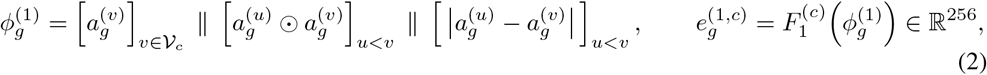

(2400 → 512 → 256 under Core+K+X). A view-pairing shuffle control separates genuine paired-input benefit from generic nonlinear capacity: breaking gene-wise pairing in a capacity-matched twin costs +0.0259 in paired task-equal Macro-F1 (Appendix 28.3).

### Low-rank higher-order residual

The second expert augments the pairwise representation with a controlled higher-order path,

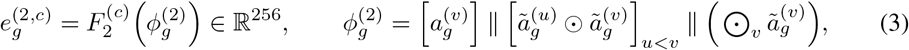

with 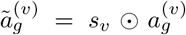 and per-view factors *s*_*v*_ ∈ ℝ^*h*^ (|*V*_*c*_| · *h* = 480 parameters, initialized *N*(1.0, 0.08^2^)). The resolved implementation is a *diagonal*, single-full-term specialisation of a general factorized multilinear residual; rank and factor structure are fixed by the released snapshot. These are representation-level interaction terms, not molecular interaction claims.

### Reliability experts (input-conditioned view-weighting expert)

The third expert is, in the resolved implementation, a per-gene input-conditioned convex weighting over adapted view vectors:

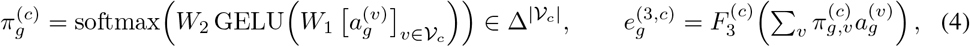

with *W*_1_ ∈ ℝ^96*×*480^ and *W*_2_ ∈ ℝ^5*×*96^ under Core+K+X. The name denotes an input-conditioned view-weighting expert; the weights are not gene importance, assay-quality calibration, or biological reliability in a causal sense. A *branch-level* router over the three experts is prespecified but unvalidated; its equations, falsifiable predictions, and confirmation design live in Appendix 25, and nothing in the main text depends on it.

### Label-free objective and leakage boundary

Representation fitting uses no downstream labels; the objective combines variance (anti-collapse), covariance (decorrelation), cross-view alignment, and availability-consistency terms with the coefficients above. For an outer split (*G*_tr_, *G*_te_),

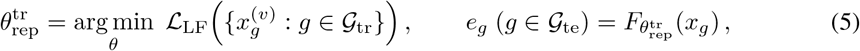

and the probe is fitted on _tr_ labels and evaluated once on _te_. Every number reported below comes from one frozen configuration whose files are released, and no quantity was selected by comparing it against the reported metric. The experts form a progressively richer *architectural* ladder (concatenation → pairwise → higher-order residual → input-conditioned weighting); this containment is a design statement, not a tested-risk ordering, and the routed top of the ladder lies outside the fixed-expert evaluation (Appendix 25).

## 4 Experiments

The evaluation tests whether the preferred fusion bias changes across controlled input conditions. The internal study uses the completed summary of `core_k_x_fulltask_20260908`; the external track uses the completed RF comparison `benchmark_gene_external_2`. All intervals are descriptive task-bootstrap intervals over the register (10,000 cluster resamples over tasks after seed/fold aggregation, fixed seed; Appendix 9); they summarize register heterogeneity and are not pairwise significance tests or population-level inference (evidence grade G1; the four grades are restated in Appendix 1).

### Task register and evaluation scope

The eligibility-filtered task set contains 54 of 79 candidate tasks (9 binary, 42 multiclass, 3 multilabel); the reported register is the completed subset, **fifteen tasks**—nine binary plus six completed multiclass tasks—under four conditions with six primary operators, three seeds, and three outer folds (3,240 records), plus 162 attention records under Core+K+X. It is fixed before score inspection (Table 1); task descriptions and gene counts are in Appendix 7, binary9 is retained only for continuity diagnostics, attention, and the external RF track, and the remaining 39 eligible tasks contribute no numbers (Appendix 3).

**Table 1:** The fifteen-task register by family; per-task descriptions and counts are in Appendix 7. Definitions precede score inspection.

| Family | Tasks | Type | Labeled genes |
| --- | --- | --- | --- |
| complex/network | CCD Protein/Transcript, N1 network/targets | binary | 280–1,579 |
| TF/chromatin | TF vs non-TF, bivalent pairs, dosage/range TF | binary | 127–2,652 |
| expression cluster | Blood, Brain, Cell-line | multiclass | 11,881–18,460 |
| pathology | Breast, Cervical, Colorectal | multiclass | 16,156–16,566 |

### 4.1 Controlled view additions change the measured task profile

Adding K and X changes the measured task profile in different ways, rather than producing interchangeable “extra-view” gains: on the fifteen-task register, PCA then concat changes by +0.0492 from Core to Core+X, +0.0332 from Core to Core+K, and +0.0629 in full. The family decomposition below shows the X response concentrated in expression-cluster tasks and the K response strongest in complex/network tasks; complete task-level values and register accounting are in Appendix 10. The practical consequence is that a single-input-set leaderboard cannot speak to fusion bias in the other input states.

### 4.2 Fusion preferences change across protocol conditions

The four conditions are protocol states on the same shared gene universe, not natural missing-view patterns. Table 2 reports the task-equal means: the leader changes with the condition (the input-conditioned view-weighting expert under Core and Core+K, PCA then concat once X enters), yet all descriptive intervals overlap (widths ≈ ± 0.13), so no operator separates from its nearest competitor. The result is therefore a condition-dependent ranking pattern rather than an operator win.

**Table 2:** Task-equal Macro-F1 by protocol condition over the fifteen-task register (six operators, three seeds, three folds); boldface marks the per-condition leader. All intervals overlap (widths ≈ ±0.13).

| Method | Core | Core+K | Core+X | Core+K+X |
| --- | --- | --- | --- | --- |
| Linear concat | 0.4358 | 0.4446 | 0.4547 | 0.4586 |
| PCA add | 0.3832 | 0.3718 | 0.3846 | 0.3765 |
| PCA then concat | 0.4560 | 0.4892 | **0.5052** | **0.5188** |
| Interaction MLP | 0.4943 | 0.5029 | 0.5005 | 0.5112 |
| Low-rank higher-order | 0.4895 | 0.5043 | 0.5025 | 0.5122 |
| Reliability experts | **0.4962** | **0.5049** | 0.5025 | 0.5142 |

#### Matched condition effects are non-additive

Table 3 reports matched condition contrasts. PCA then concat has the largest Core-to-Core+K+X change (+0.0629), roughly three times the changes of the other operators (− 0.0067 to +0.0228). The non-additivity term *N*_*KX*_ = *M*_Core+K+X_ − *M*_Core+K_ − *M*_Core+X_ + *M*_Core_ is an *arithmetic recorder* of the condition response: it holds for any four numbers and by itself separates nothing about mechanism. What it exposes descriptively is that the operator with the largest gain is also the least decomposable (*N*_*KX*_ = − 0.0195), whereas the learned experts are near-additive (+0.0022 to +0.0030; Low-rank higher-order − 0.0051); reading only the Core+K+X column would reverse the conclusion. These are protocol-state changes under a fixed design, not causal effects; the binary sub-register shows the same ordering with larger magnitudes (Appendix 10).

**Table 3:** Matched condition deltas relative to Core and the non-additivity term *N*_*KX*_ over the fifteen-task register; boldface marks the largest gain per contrast. Descriptive decompositions, not causal effects or significance tests.

| Method | $\Delta_K$ | $\Delta_X$ | $\Delta_{KX}$ | $N_{KX}$ |
| --- | --- | --- | --- | --- |
| Linear concat | +0.0088 | +0.0189 | +0.0228 | -0.0049 |
| PCA add | -0.0114 | +0.0014 | -0.0067 | +0.0034 |
| PCA then concat | **+0.0332** | **+0.0492** | **+0.0629** | **-0.0195** |
| Interaction MLP | +0.0086 | +0.0061 | +0.0169 | +0.0022 |
| Low-rank higher-order | +0.0149 | +0.0130 | +0.0227 | -0.0051 |
| Reliability experts | +0.0087 | +0.0062 | +0.0179 | +0.0030 |

#### Multiplicity note

With intervals of width ≈ ± 0.13 over a 6-operator × 4-condition grid, top-operator swaps and extreme *N*_*KX*_ values are expected under noise; the cleanest no-dominance observation is distributional (no leader wins more than eight of fifteen per-task races; thirteen of fifteen tasks switch winners; Figure 10), and Table 3 is a map of where re-measurement is cheap, not a decision procedure.

#### Task-family composition explains the aggregate response

The source of the condition dependence is visible at family level (Table 17, Appendix 10): the expression-cluster family is driven by the single-cell view (PCA then concat: 0.0540 → 0.0982 from Core to Core+K+X), the complex/network family shows the largest K response (0.5631 → 0.6811), the TF/chromatin family is led by the learned experts throughout (≈ 0.795 at Core+K+X), and the pathology family sits at the constant-prediction floor (Macro-F1 ≈ 0.327) in every condition, contributing coverage rather than condition signal. Each family thus has an interpretable condition response, and the condition-dependent ranking emerges from their composition.

#### The registered semantic-leakage control preserves the pattern

The knowledge view embeds curated functional text and five register tasks draw labels from overlapping curation resources, so a semantic-leakage path exists by construction (Appendix 21). A registered control recomputes every condition contrast after removing all five overlapping tasks and bounds the concern: the K-condition gain is not concentrated in the overlapping group (per-task Δ_*K*_: PCA then concat +0.0260 vs. +0.0368; Reliability experts +0.0041 vs. +0.0110), and on the ten-task non-overlapping register every qualitative conclusion is unchanged (PCA then concat remains the largest gainer, Δ_*KX*_ = +0.0600, and least decomposable, *N*_*KX*_ = − 0.0214). The control does not certify the K view as leak-free, and K-condition gains are still not interpreted mechanistically.

### 4.3 An independent probe preserves the numerical range

The external track evaluates the nine binary tasks under Core+K+X with a separately configured RF probe (200 trees, depth 12, balanced subsample), frozen before inspection of the internal results. Table 4 reports the nine completed entries: the top three cluster within 0.006 and all intervals overlap, so the track identifies no statistical winner. The RepGene entries exceed the PRISME+scRNA protocol extension (a five-view extension, not the original method), the GenePT text-only reference, and a random 256-D control. This track is the paper’s independent soundness check: a probe configured before the internal results reproduces the numerical range without reproducing a winner.

**Table 4:** External RF track over the nine binary tasks under Core+K+X: task-equal Macro-F1 with descriptive 95% intervals (two decimals); the probe was configured before the internal results were inspected.

| Method | Macro-F1 [95%] |
| --- | --- |
| Random 256-D | 0.4240 [0.41, 0.45] |
| PCA add | 0.5103 [0.46, 0.57] |
| GenePT (text only) | 0.5479 [0.48, 0.64] |
| Linear concat | 0.6342 [0.54, 0.73] |
| PRISME+scRNA ext. | 0.7116 [0.62, 0.80] |
| Interaction MLP | 0.7192 [0.63, 0.81] |
| PCA then concat | 0.7221 [0.62, 0.81] |
| Reliability experts | 0.7227 [0.63, 0.82] |
| Low-rank higher-order | **0.7251** [0.63, 0.81] |

#### Metric-sensitivity check

Macro-F1 is the prespecified primary metric, and the ordering is *not* preserved across metrics: the three learned experts lead under balanced accuracy in all four conditions (e.g., Core+K+X: Reliability experts 0.5224), while PCA then concat is numerically highest under ROC-AUC in all four (0.7299–0.7894; Appendix 14). The mechanism is threshold sensitivity on class-imbalanced tasks (frozen 0.5 threshold versus rank-integrated AUC), confirmed by a registered extension (Appendix 28.4): PCA then concat ranks best but is calibrated worst (ECE 0.1270), losing at the frozen threshold (0.6866 vs. 0.7045) and winning once thresholds are tuned (0.7708 vs. 0.7372). The picture is therefore metric-, probe-, and threshold-dependent; Macro-F1 remains primary by pre-registration, and no superiority claim is made under AUC.

### 4.4 Gene-level utility does not transfer automatically to cells

Gene-level and cell-level utility are separate estimands: the diagnostic aggregates gene representations to cell level on one aorta annotation task (expression-weighted aggregation, library-size normalization, logistic-regression classifier, five seeds). As Table 5 shows, the picture repeats the gene level (expression-only reference leading, and a dimension-matched random control within 0.004 of the best fused representation). The inversion follows from the aggregation algebra: writing *p*_*c*_(*g*) = *x*_*cg*_*/* ∑_*j*_ *x*_*cj*_ and decomposing *e*_*g*_ = *µ* + *b*_*t*(*g*)_ + *ε*_*g*_, the cell representation 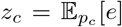 satisfies

**Table 5:** Gene-to-cell boundary diagnostic on the aorta annotation task: Macro-F1 over five seeds (± seed SD). The expression-only reference leads and a dimension-matched random 256-D control sits within 0.004 of the best fused representation; one fixed design, not a validated method.

| Representation | Macro-F1 |
| --- | --- |
| Expression only | **0.9295** ± 0.0047 |
| PCA on expression | 0.9123 ± 0.0101 |
| Linear concat | 0.9115 ± 0.0136 |
| PCA then concat | 0.9091 ± 0.0104 |
| PCA add | 0.9078 ± 0.0138 |
| Low-rank higher-order | 0.9024 ± 0.0129 |
| Random 256-D control | 0.8987 ± 0.0100 |
| Interaction MLP | 0.8985 ± 0.0241 |
| Reliability experts | 0.8889 ± 0.0246 |

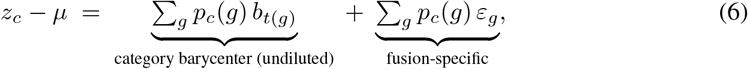

where *S*_*c*_ = ∑_*g*_ *p*_*c*_(*g*)^2^ = 1*/N*_eff_ (*c*) is the inverse participation ratio and the fusion-specific component obeys the root-mean-square bound 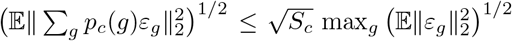 under zero-mean, uncorrelated cross-gene residuals—not an algebraic identity for arbitrary vectors (derivation and counterexample in Appendix 13). The channel the cell task rewards—which genes are expressed, and how much—passes through undiluted, even for random embeddings, while uncorrelated fusion-specific structure is attenuated by roughly two orders of magnitude, below the seed noise (0.01–0.02).

### 4.5 Negative results and interpretation limits

Three negative results are retained as results. (i) *No operator dominates under this contract*: intervals overlap in every condition. (ii) *Token-wise attention does not lead under this contract* (binary sub-register, Core+K+X: cross-attention 0.7152, transformer encoder 0.7115, both below PCA then concat’s 0.7238). The granularity itself explains this—one vector per view per gene leaves attention a mixture over at most _*c*_ |*V* | ≤ 5 whole-view tokens—and the budget extension rules out undertraining as an account: cross-attention never leads at any tested budget, and the residual gap stays within the seed-noise band (Appendices 28.2, 11). (iii) *Explicit multiplicative interaction does not clear its capacity-matched ceiling on real tasks*: the historical screening stays below the ceiling (Appendix 24), reported as magnitude-only evidence because fold-level records are unavailable.

#### Scope boundaries and their controls

Each boundary below is paired with the control that makes it auditable. Scope is bounded deliberately: the claims hold for the fifteen-task register, two recorded RF protocols, and registered input states; task-equal aggregation pools seeds and folds *before* the task-level bootstrap, so condition swaps are not verified per fold or per seed. Register tasks share annotation families, so the fifteen are correlated evidence, and class differences are recorded as such rather than decomposed into bias versus capacity (Section 4.2). The probe, budget, pairing, and threshold treatment are varied in registered extensions (Appendix 28): the structural findings survive, while the leading operator and the threshold ordering are protocol-dependent.

## 5 Discussion

Two descriptive statements carry the reported results. First, under one frozen probe and aggregation rule, the top-scoring operator changes with the protocol-defined input condition, and no operator is separated from its nearest competitor by a task-bootstrap interval. Second, matched condition contrasts are operator-dependent and structured by task family (Section 4.2): thirteen of fifteen tasks switch winners, and the two exceptions keep PCA then concat throughout, so family composition, not noise, drives the aggregate. The register cannot recommend an operator for unseen tasks; it defines a low-cost comparison in which the contract is applied and the condition response measured, with conditional-selection predictors left as hypotheses (Appendix 25). The text view and several labels overlap curated resources, so a semantic-leakage path exists by construction, bounded by the registered control.

All learned operators share one label-free objective and an eight-epoch budget (8–16 gradient updates per fit), so under-trained capacity is an alternative explanation for disadvantages of heavier operators; the budget extension narrows this for cross-attention, which improves with budget but never leads (Appendix 28.2). Three objections are answerable from the same evidence: whether the budget is sufficient (all operators improve with more budget and the ordering does not change, Appendix 28.2); whether the text view leaks labels (the registered removal control preserves every qualitative conclusion, Section 4.2); and whether the ordering is metric- or threshold-dependent (it is, jointly on probe and threshold, which is why Macro-F1 is pre-registered, Section 4.3). The evidence grades remain explicit (Appendix 1): G1 Figure 2, G2 Table 4, G3 the dilution bound (Eq. 6), and G4 the magnitude-only multiplicative results. A new operator enters by exporting a 256-D representation for the same gene universe and rerunning the frozen probe; changing folds, seeds, probe, or aggregation breaks comparability. This fairness claim is relative to the shared adapters and budget, not absolute across capacities: the protocol card (Appendix 19) specifies the remaining asymmetries, the probe family, budget, and pairing control are reported as registered extensions (Appendix 28), and adapter-matched baselines, gradient-boosted probes, and multiclass epoch sweeps remain legal entries left to future work.

**Figure 2:**
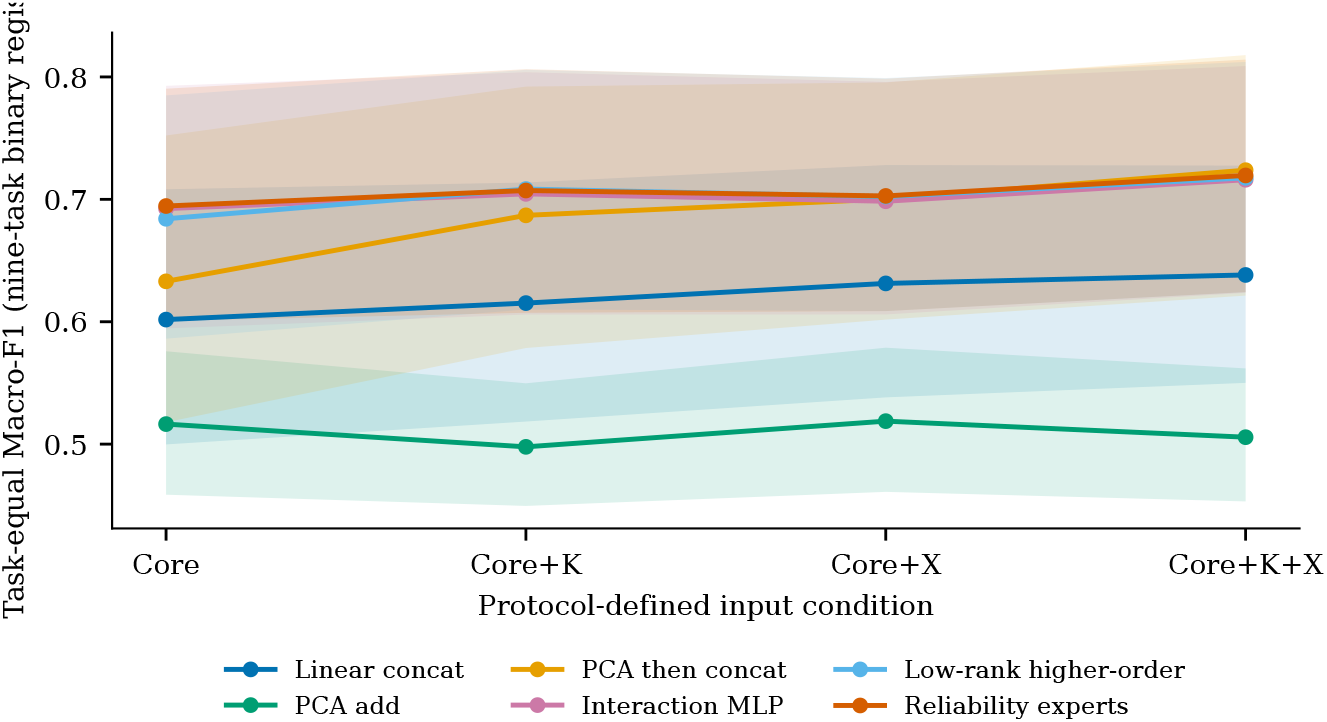
Binary-sub-register diagnostic (not the primary full-register result): task-equal Macro-F1 across the nine registered binary tasks and four protocol-defined conditions. Shading indicates descriptive 95% task-bootstrap intervals; no auxiliary dashed curves are plotted. The top-scoring operator changes with the condition, and intervals overlap throughout. The corresponding fifteen-task analysis is reported in Appendix 10.

## 6 Conclusion

Answering the four questions of Figure 1 in sequence: adding K and X does not produce interchangeable condition responses, and family composition determines the aggregate profile. The preferred operator changes with the protocol state—Reliability experts lead Core and Core+K, PCA then concat leads Core+X and Core+K+X—while all descriptive intervals overlap, thirteen of fifteen tasks switch winners, and the semantic-leakage control preserves the pattern; the independently configured external RF track reproduces the numerical range but identifies no statistical winner; and one gene-to-cell diagnostic places fused representations near a dimension-matched random control, consistent with attenuation of fusion-specific residuals, not a general cell-level law. The protocol recommends no operator for unseen tasks and certifies no mechanism: it is an evidence-graded audit framework for condition-bounded fusion claims.

## Impact Statement

This work studies how the complexity of a gene-representation fusion operator should be compared under a fixed, matched evaluation protocol. Its intended positive impact is methodological: it provides a way to compare additive, pairwise-interaction, low-rank higher-order, and input-conditioned fusion without treating a single architecture as universally optimal, and it demonstrates reporting discipline (claim–evidence ledger, evidence grades) that other audits can reuse. The study is not intended to provide clinical decision support, diagnose disease, or establish biological mechanisms from representation geometry or routing weights.

The principal risks concern overinterpretation and downstream misuse. A higher score on the reported gene-level tasks could be incorrectly presented as evidence that a view is biologically causal, that a higher-order term corresponds to a molecular interaction, or that a weighting vector measures assay quality. The four conditions are protocol-defined matched states, not natural missingness or deployment robustness; the register is restricted to fifteen tasks (nine binary, six multiclass) on an aligned five-view gene intersection that over-represents well-studied genes; and gene-level utility does not automatically transfer to cell-level endpoints. Safeguards: retain the manifests; fit all preprocessing on outer-training data only; report condition-specific comparisons with their intervals; and disclose evidence grades, including the limited provenance of the historical sweep.

## AI Use Statement

Generative AI was used as an assistive tool during manuscript preparation to draft and revise English prose in the Introduction, Related Work, Approach, Experiment, Conclusion, and this statement; to suggest organization and transitions consistent with the planned ICLR paper structure; to inspect and summarize author-provided planning documents and experiment records; and to assist with LaTeX file organization, cross-reference checks, and compilation-error diagnosis. These uses correspond to manuscript drafting/editing, literature and document summarization, research-methodology and experiment-organization feedback, and software/formatting assistance under the ICLR 2027 AI policy.

Generative AI was not used to generate synthetic datasets, fabricate or alter experimental results, create labels, select test results after observing outcomes, or replace the authors’ responsibility for the research design, implementation, statistical analysis, interpretation, or final claims. The numerical results and evidence boundaries reported here were taken from the project artifacts identified in the claim–evidence manifest. All AI-assisted text, code-related suggestions, citations, cross-references, and scientific claims must be checked by the authors against the underlying files and executed compilation or experiment records before submission. The authors retain full responsibility for the accuracy, originality, integrity, and compliance of the final manuscript and artifact, including any content produced with generative-AI assistance.

## Reproducibility Statement

Code and artifacts are available from the anonymized repository at https://anonymous.4open.science/r/RepGene-E0A8/. The released embedding data are available at https://drive.google.com/drive/folders/1a86OycK98xvCq7ZNaDWZjtpyu2T-dghh?usp=drive_link; data and upstream model access remain subject to their original licenses.

The released artifact freezes, in its current state: (i) the resolved configuration files of the primary evaluation (`core_k_x_fulltask_20260908`, hidden width 96, eight epochs, seeds 41– 43, three outer folds) and of the external RF track (`benchmark_gene_external_2`, 200 trees, depth 12, balanced subsample); (ii) the per-task, per-condition, per-method summary tables (3,240 internal records on the fifteen-task register; 729 external fold-level records for the nine main entries, plus 162 attention records, 891 in total); (iii) the figure source CSVs with a SHA-256 `source_manifest.json`, from which every figure regenerates without re-running any model; (iv) the claim–evidence ledger (Appendix 15), figure-to-claim map (Appendix 17), and protocol decision log (Appendix 18); and (v) the embedding-geometry analysis scripts and tables behind Appendix 26. Exact parameter counts in Table 8 are computed from the released model definitions for the seven operators whose configurations are frozen in the snapshot; the transformer-encoder configuration gap is recorded explicitly.

Known gaps are recorded rather than promised away: peak-memory and FLOP measurements are *not* part of this package; the multiplicative-sweep summary values lack fold-level records and are interpreted as magnitude-only supplementary evidence; and a clean-environment re-run, immutable data-access instructions, and identifier-mapping checksums are packaged with the supplementary bundle rather than re-derived here. The scope limitations are listed in Appendix 21.

## Ethics Statement

This study analyzes previously collected gene-level and single-cell resources and makes no clinical claim. No individual-level clinical decision is produced by the reported models. Model weights, controlled condition contrasts, and geometry diagnostics are not interpreted as biological mechanisms. Any public release must document data-access conditions, licenses, upstream model exposure, identifier mappings, and restrictions on redistribution. If later work applies the representations to human or clinically linked data, that work should separately establish the applicable institutional review, privacy, security, fairness, and data-governance requirements; those requirements are not established by this benchmark.

## Appendix outline

This appendix gives the definitions, algorithmic derivations, evaluation scope, complete results, and evidence boundaries needed to interpret every number in the main text. Appendix 1 restates the four evidence grades used in the main text; Appendix 2 gives the extended related work; Appendix 3 gives the experimental accounting; Appendix 4 specifies the views; Appendix 5 specifies the operators with exact parameter counts; Appendix 6 states the label-free objective; Appendices 7–9 list tasks, data processing, and metrics including the bootstrap algorithm; Appendix 10 reports complete per-task results, the task-family stratification, and the semantic-leakage control table; Appendix 11 covers the two attention operators and their granularity mismatch; Appendix 12 details the external track; Appendix 13 derives the gene-to-cell aggregation; Appendix 14 reports secondary metrics; Appendix 26 adds an embedding-geometry analysis of the five views and the eight fused representations; Appendices 15–18 state the evidence boundaries, artifact definitions, figure-to-claim correspondence, and design decisions; Appendix 19 specifies the comparison contract; Appendix 21 collects all limitations; Appendix 27 collects additional evaluation figures; Appendices 24–25 report supplementary multiplicative-interaction results and define the dynamic-routing evaluation criteria; and Appendix 28 reports four registered protocol extensions that vary the probe family, the training budget, the gene-wise pairing of the interaction expert, and the calibration/threshold treatment of the stored probabilities.

### 1 Evidence grades used throughout this paper

All register-level statements in this paper carry one of the four evidence grades below, defined with the scope of the test that produces them. The main text marks each result with its grade and refers here for the definitions.

#### The four grades

Every register-level statement carries one of the four grades defined here. **(G1) Register-level descriptive**: task-equal means with task-bootstrap intervals over the fixed register (Section 4); not population inference. **(G2) Independently configured external check**: the same tasks under a separately configured probe, frozen before internal results were inspected (Section 4.3). **(G3) Boundary diagnostic**: a single-design-point probe of where a claim stops holding; not a validated method. **(G4) Magnitude-only supplementary**: historical summary-level records without fold-level data (Appendix 24).

### 2 Extended Related Work

#### Gene representations across biological views

Gene representations have been constructed from co-expression, sequence, protein, interaction, and literature-derived signals. Gene2vec learns distributed gene representations from co-expression structure (Du et al., 2019), and a systematic evaluation of input choices shows that the utility of a modality depends on the downstream task and its data-generating process (Brechtmann et al., 2023). Complementary sequence- and network-based resources, including ProtVec, GeneMANIA, HumanNet, and STRING, provide established biological views (Asgari & Mofrad, 2015; Mostafavi et al., 2008; Hwang et al., 2019; Szklarczyk et al., 2023). Text-based gene representations such as GenePT use language-model-derived biomedical knowledge (Chen & Zou, 2025). These studies motivate heterogeneous views, but they do not determine how a fixed fusion operator should respond when the statistical roles of the views change.

Biological foundation models supply strong modality-specific encoders: protein language models (Elnaggar et al., 2022; Rives et al., 2021), structure-informed representations (Jumper et al., 2021), single-cell models (Cui et al., 2024; Yang et al., 2022), transfer from molecular to network-level prediction (Theodoris et al., 2023), and integration workflows (Wolf et al., 2018; Lopez et al., 2018; Korsunsky et al., 2019; Hie et al., 2019; **?**; Welch et al., 2019a). The present study does not replace these upstream encoders; it treats their aligned outputs as input views and isolates the downstream fusion question.

#### Multiview fusion and higher-order interactions

Standard concatenation retains view identity and delegates cross-view relationships to a subsequent predictor. Additive fusion assumes contributions can be aligned and combined coordinate-wise. Explicit multiplicative or difference features expose non-additive pairwise relationships, and higher-order parameterizations can represent joint dependencies among more than two views. This family has canonical roots in multimodal tensor fusion: the Tensor Fusion Network (Zadeh et al., 2017) enumerates unimodal, bimodal, and trimodal interactions through an explicit outer product, and Low-rank Multimodal Fusion (Liu et al., 2018) factorizes the high-order interaction tensor into modality-specific low-rank factors, which is the direct conceptual ancestor of the diagonal low-rank residual evaluated here (Eq. 3). Additional expressivity can reduce approximation bias when joint information is useful, but it can also increase parameter count and estimation variance when evidence is weak or conflicting. Attention-based fusion is the default choice in much of the multimodal literature (Vaswani et al., 2017; Tsai et al., 2019; Baltrusaitis et al., 2018), where fused models are trained end-to-end with task labels; our label-free, probe-matched, one-vector-per-view setting is a different operating point, and Appendix 11 analyses the consequence. Among molecular integration platforms, PRISME (Zheng et al., 2025) couples heterogeneous molecular embeddings through an autoencoder trained under its own reconstruction objective; because the primary contract requires every entry to export 256-D representations fitted under the shared label-free objective, PRISME enters only through the external-track five-view protocol extension (Appendix 12), and a native PRISME entry remains an open extension rather than a controlled comparison.

#### Label-free objectives and conditional selection

Unsupervised or self-supervised fitting is useful when task labels should not determine the representation before downstream evaluation. Label-free input descriptors—agreement/conflict statistics, availability masks, uncertainty proxies—can drive input-conditioned weighting, but such weights are not gene importance, assay quality, causal contribution, or calibrated biological reliability. Selection analyses must separate fixed experts, static mixtures, learned routers, and label-observing oracles; the oracle is an upper-bound diagnostic, never a deployable method (Appendix 25).

#### Incomplete views and evaluation estimands

Multimodal biological learning has considered integration, imputation, shared-specific representations, and genotype–phenotype prediction under heterogeneous measurements (Chandrashekar et al., 2023; Khodaee et al., 2025; Wu et al., 2026). Multi-omics factor models and deep generative approaches—MOFA, MOFA+, totalVI, MultiVI, and CITE-seq-style joint measurements—address shared and modality-specific structure (Argelaguet et al., 2018; 2020; Gayoso et al., 2022; Ashuach et al., 2023; Stoeckius et al., 2017). These settings do not share the estimand of the controlled comparison here: the Core/Core+K/Core+X/Core+K+X states are registered protocol conditions on a common gene universe, not per-gene observations of natural assay availability, imputation quality, or deployment robustness.

#### Leakage-aware evaluation

Representation comparisons are sensitive to leakage because upstream encoders, labels, networks, and preprocessing statistics can share data sources (Luecken & Theis, 2019; Tran et al., 2020; Soneson & Robinson, 2018; **?**; Hafemeister & Satija, 2019). The protocol therefore fits all preprocessing and label-free objectives on outer-training features only, trains the probe on outer-training labels, and evaluates once on the held-out fold. Task-equal aggregation is applied after seed and fold aggregation so that a task with more genes does not dominate the register.

#### Task-set and evaluation accounting

A three-fold eligibility audit retains 54 of 79 candidate tasks: 9 binary, 42 multiclass, and 3 multilabel. The registered matrix of 54 × 4 × 6 × 3 × 3 = 11,664 task–fold evaluations is the protocol plan, not completed work. This revision completes fifteen tasks: the nine binary tasks plus six multiclass tasks from the expression-cluster and pathology-prognostics families (3,240 internal records, plus 162 attention records on the binary sub-register; 729 external records for the nine main entries, plus 162 attention records, 891 in total). The 39 remaining eligible tasks are excluded by completion status—only these fifteen tasks were run to completion under this contract—not by any result-dependent selection, and no number from them appears anywhere in this paper.

### 3 Experimental Accounting

The primary evaluation uses five aligned views grouped as Core (DNA, RNA, protein), K (knowledge), and X (single-cell expression), retains 18,460 aligned genes, and evaluates fifteen registered tasks (nine binary, six multiclass). Six primary operators are evaluated under all four conditions; two additional attention operators are evaluated under Core+K+X only, on the binary sub-register:

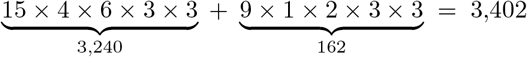

task–condition–method–seed–fold records, with seeds 41, 42, 43 and three outer folds. Representation fitting is executed once per (task, condition, operator, seed, outer fold) cell: stratified outer folds are constructed within each task’s labeled gene set, and the fitting pipeline (seeded at the job seed plus fold index) is re-run inside every cell, giving 15 × 4 × 6 × 3 × 3 = 3,240 primary fits plus 9 × 1 × 2 × 3 × 3 = 162 attention fits (3,402 in total). The three seeds and three folds therefore both index distinct fits; no representation is shared across tasks, and no fit observes a task label or a heldout gene. Task-equal summaries assign equal weight to registered tasks; they do not treat genes, folds, or rows as independent biological populations, and all intervals are descriptive task-bootstrap intervals over the register tasks (10,000 resamples, fixed seed), never significance tests.

The external RF track contains nine binary tasks, three seeds, and three folds. Nine completed non-attention entries contribute 9 × 9 × 3 × 3 = 729 fold-level records; the two attention operators add 9 × 2 × 3 × 3 = 162, giving 9 × 11 × 3 × 3 = 891 records across all eleven entries in the Core+K+X state. Table 4 in the main text reports the nine non-attention entries; Table 20 reports the two attention operators.

### 4 Views and Protocol Conditions

**Table 6:**
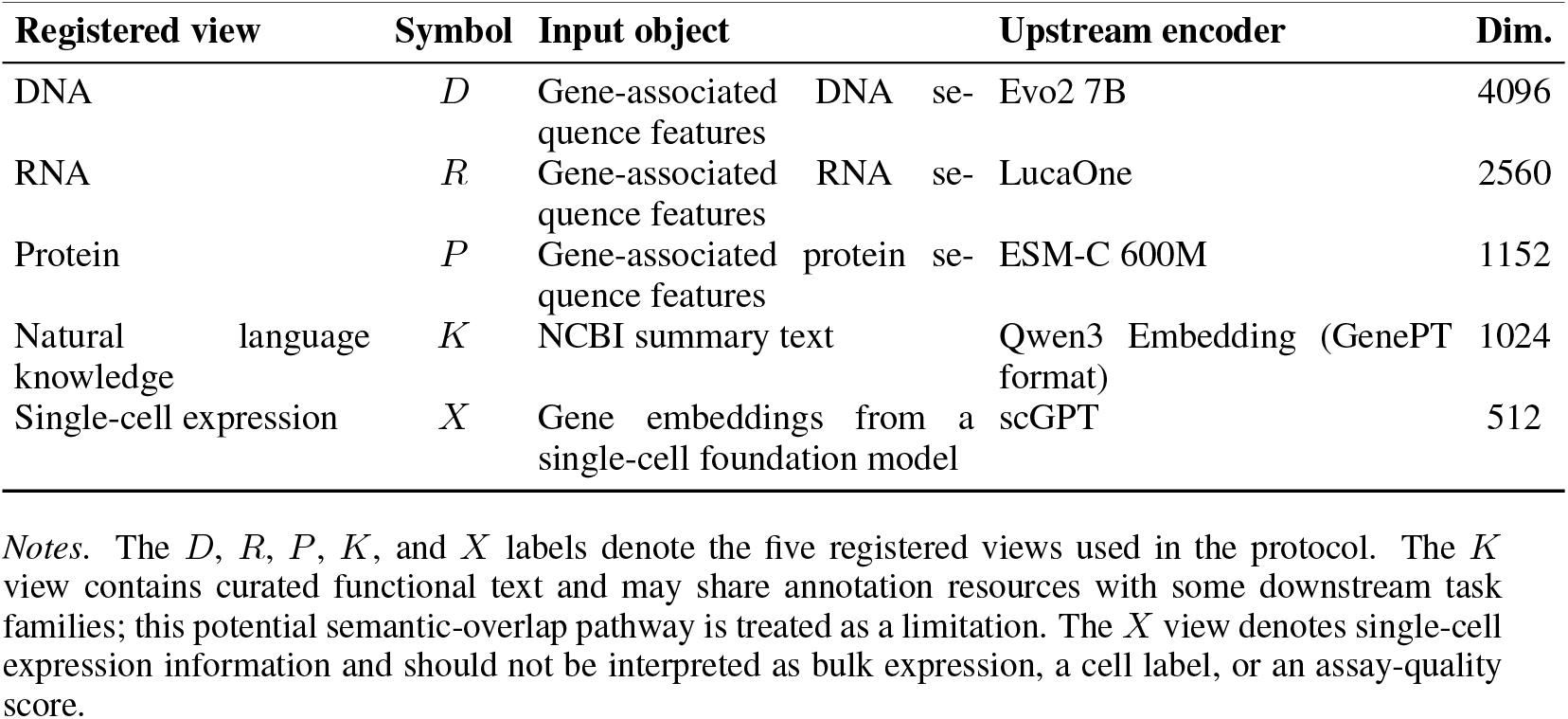
Registered biological views and upstream representation sources. Each input vector is defined at the gene level before shared per-view adaptation; all fused representations are exported at 256 dimensions.

Core, Core+K, Core+X, and Core+K+X are protocol-defined input states applied to a *shared* gene universe. They are not natural per-gene coverage, not imputation experiments, and not deployment robustness tests. Every condition uses the same genes, the same outer splits, the same seeds, and the same probe.

### 5 Fusion Operators and Hyperparameters

#### Resolved implementation notes (exact, reconstruction-ready)

All learned operators share per-view adapters Linear(*d*_*v*_ → 96) + LayerNorm + GELU + Dropout(0.1) and a two-layer fusion head, differing only in the feature map feeding that head. **Interaction MLP** concatenates the adapted views with all pairwise products and absolute differences (2400 → 512 → 256 under Core+K+X). **Low-rank higher-order** learns one diagonal scale vector *s*_*v*_ ∈ ℝ^96^ per view (init *N*(1.0, 0.08^2^); |V_*c*_| · 96 parameters), forms pairwise products of the rescaled views plus a single element-wise full _*c*_-way product, and feeds the concatenation (1536 features) to the same head—a diagonal, single-full-term specialisation of the general factorized residual. **Reliability experts** computes pergene softmax weights over views with a gate Linear(480 → 96) + GELU + Linear(96 → |*V*_*c*_|) applied to the adapted concatenation and mixes the adapted views convexly before the head; it is a view weighting, not a branch router. **Cross-attention** applies one multi-head attention block (4 heads, width 96) over the view stack with mean pooling. The exact learned-parameter counts appear in Table 8. The three baselines operate on the fold-standardized raw views with closed-form PCA/linear maps fitted on outer-training features only; they never touch the trained adapters, so their learned-parameter count is exactly zero. The learned experts’ counts include the shared adapters (898,464 parameters across five views) plus the operator-specific fusion path. A reconstruction decoder branch is active during fitting with unit weight (Section 3) but is never used at export time; its parameters are training-only and are excluded from these counts. For the transformer-encoder operator, the block count, attention-head count, and feed-forward width were not frozen into the released configuration snapshot, so its exact learned-parameter count cannot be reconstructed from the released artifacts and is recorded as a reproducibility gap (Appendix 21).

**Table 7:** Six primary fusion biases and two additional attention operators. All operators share the view adapters, the 256-D output budget, the folds, the seeds, the label-free objective, and the probe; the attention operators are evaluated under Core+K+X only.

| Family | Operator | Fusion bias |
| --- | --- | --- |
| Baseline | Linear concat | Concatenate fold-standardized raw views, outer-train PCA projection to 256-D |
|  | PCA add | Per-view PCA to 256-D on fold-standardized raw views, then average across views |
|  | PCA then concat | Per-view PCA on fold-standardized raw views (256-D budget split across views: 86/85/85 under Core; 64 per view under four-view conditions; 52/51/51/51/51 under Core+K+X), then concatenation |
| RepGene | Interaction MLP | Explicit pairwise products and absolute differences (Eq. 2) |
|  | Cross-attention | Directed cross-view read-out with residual (Eq. 16) |
|  | Transformer encoder | Permutation-equivariant self-attention stack with mean pooling (Eq. 18) |
|  | Low-rank higher-order | Low-rank residual over higher-order interactions |
|  | Reliability experts | Per-gene input-conditioned softmax view weighting (Eq. 4) |

Shared training configuration: hidden width 96, dropout 0.1, eight epochs, batch size 1024, Adam with learning rate 10^−3^ and weight decay 10^−5^, and label-free weights (*λ*_var_, *λ*_cov_, *λ*_align_, *λ*_avail_) = (0.1, 0.01, 0.05, 0.1) on top of a unit-weight reconstruction term (Eq. 10). The primary random-forest probe uses 300 trees, maximum depth 16, and minimum leaf size 2. The external probe is configured separately (Appendix 12).

### Algorithmic Derivation of the Fusion Operators

This section derives the representations used in the comparison from the common per-view inputs.

Let 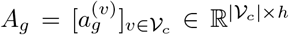, with *h* = 96. The baselines first map each view to a budgeted subspace. For PCA-then-concat, if 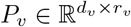 is fitted on outer-training features, then

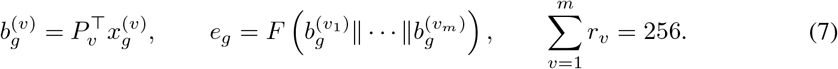

For PCA-add, each *r*_*v*_ = 256 and 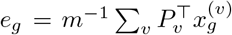; linear concat instead applies one PCA map to the concatenated standardized input. These maps are closed-form preprocessing operators and introduce no label-dependent parameters.

For the learned maps, the pairwise expert augments the view stack with all Hadamard products and absolute differences. Since each pair contributes *h* + *h* = 2*h* coordinates, its feature dimension is 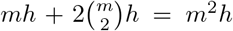; for *m* = 5, this is 2400. The low-rank map replaces each view by 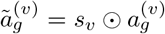, forms the pairwise products and one full product, and applies the shared head *F*_2_. The factorisation reduces the per-view scaling parameters to *mh* while retaining a controlled *m*-way interaction term.

**Table 8:**
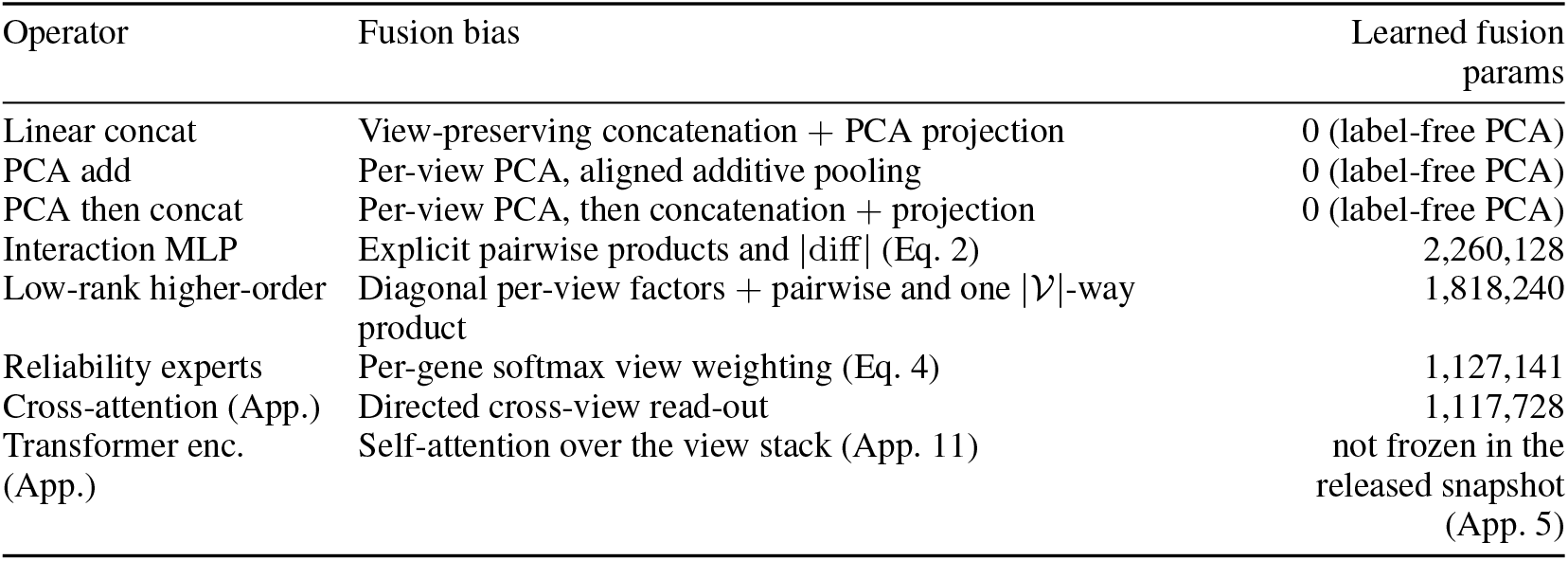
Operator specification under Core+K+X (*h* = 96, 256-D export). Learned parameters are exact counts from the released snapshot for the seven frozen configurations (adapters plus fusion path); the training-only reconstruction decoder is excluded. The transformer encoder’s block/head configuration was not frozen into the snapshot; no exact count is recoverable and we record this as a reproducibility gap rather than reconstructing post hoc (Appendix 21). Full definitions in Appendix 5.

The reliability expert is obtained by differentiating a convex-mixture parameterisation with respect to its gate logits. Let *q*_*g*_ = *W*_2_ GELU(*W*_1_ vec(*A*_*g*_)) and *π*_*g*_ = softmax(*q*_*g*_). Then 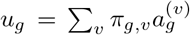 lies in the convex hull of the adapted views and *e*_*g*_ = *F*_3_ (*u*_*g*_). The Jacobian of the mixture with respect to a logit is

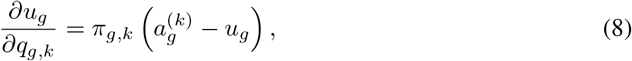

which shows that the gate learns to emphasize a view only when its adapted vector differs from the current mixture. The softmax therefore implements input-conditioned view weighting, not an unidentifiable post-hoc importance score.

All learned operators are fitted by minimizing the label-free objective on outer-training genes. With *θ* denoting adapters, fusion map, and training-only decoders,

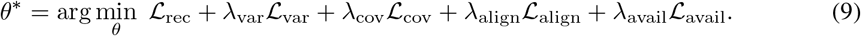

The downstream probe is then fitted independently on training labels. Thus, for a held-out gene *g* ∈ *G*_te_, *e*_*g*_ = *F*_*θ*_*\** (*x*_*g*_) contains no information from its label; this establishes the leakage boundary used by all reported Macro-F1 values.

### 6 Label-free Objective and Leakage Boundary

Training uses no downstream label. For a mini-batch *B* ⊆ *G*_tr_ of size *B*, condition view set *V*_*c*_, adapted views 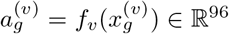, and exported representation *e*_g_ ∈ ℝ^256^ (the operator’s fused output before the probe), the resolved objective is

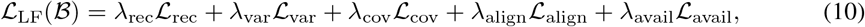

with (*λ*_rec_, *λ*_var_, *λ*_cov_, *λ*_align_, *λ*_avail_) = (1, 0.1, 0.01, 0.05, 0.1). The five terms are defined as follows.

#### Reconstruction (training-only)

Each view carries a decoder *D*_*v*_ that maps the export back to the standardized input space:

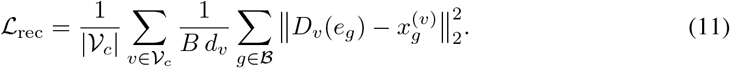

The decoders are used only by this term during fitting and are discarded at export; their parameters are excluded from the counts in Table 8.

#### Variance (anti-collapse)

Applied to the exported representation. With per-coordinate batch statistics 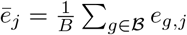 and 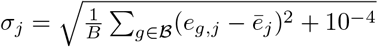,

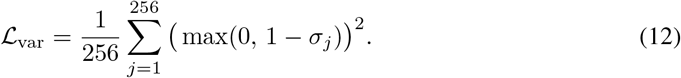

#### Covariance (decorrelation)

With the batch covariance 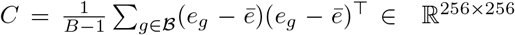,

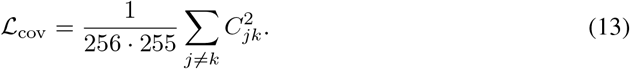

#### Cross-view alignment

Computed on the adapted view vectors (not the export). With 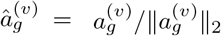 and all 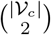 view pairs,

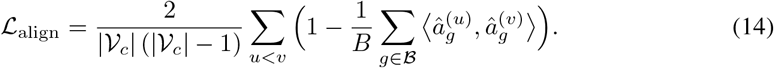

#### Availability consistency

For each gene in the batch, a view-availability mask 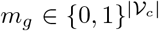 is sampled uniformly from the registered condition definitions (Core, Core+K, Core+X, Core+K+X, intersected with the active view set, partial auxiliary views allowed); masked views are zeroed before the fusion map. With *ê* = *e/*∥*e*∥_2_, the masked export *e*_*g*_(*m*_*g*_), and stop-gradient sg(·),

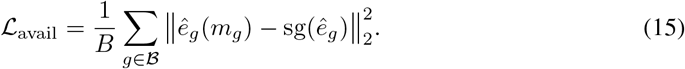

Standardization, dimensionality reduction, and every unsupervised fitting operation are estimated on outer-training features only and applied unchanged to validation and test folds. No component observes test labels, test-fold features, or test-fold view availability at fitting time.

### 7 Task Register

The main text defines the fifteen-task register compactly (Table 1); this appendix adds the task descriptions. No multilabel task is included in the current package.

### 8 Data Processing and Universe Construction

Views are aligned on gene identifiers before any fitting. The aligned universe contains 18,460 genes present in all five views. Per-view standardization and dimensionality reduction are fitted on outer-training features. Identifier mapping, source versions, and checksums are recorded in the released manifest; no per-gene clinical or individual-level data are used.

### 9 Metrics and Aggregation

The primary metric is Macro-F1. Task-equal aggregation averages the per-task metric with equal weight, so no task dominates because it has more genes. We additionally report balanced accuracy, ROC-AUC, and log loss in the released tables. Because the fifteen tasks are a fixed register rather than a random sample of a population, bootstrap intervals describe heterogeneity across the register; they do not support population inference.

**Table 9:** The fifteen registered gene-level tasks with descriptions. Types and labeled-gene counts are listed in Table 1.

| Task | Description |
| --- | --- |
| CCD Protein | Protein-level complex membership |
| CCD Transcript | Transcript-level complex membership |
| N1 network | Network neighbourhood membership |
| N1 targets | Regulatory-target membership |
| TF vs non-TF | Transcription-factor annotation |
| Bivalent vs non-methylated | Chromatin-state contrast |
| Bivalent vs lys4-only | Chromatin-state contrast |
| Dosage-sensitive TF | Dosage-sensitivity annotation |
| Long- vs short-range TF | Regulatory-range annotation |
| Blood expression cluster | Blood-expression cluster membership (multiclass) |
| Brain expression cluster | Brain-expression cluster membership (multiclass) |
| Cell line expression cluster | Cell-line expression cluster membership (multiclass) |
| Pathology prognostics - Breast cancer | Pathology prognostic class, breast cancer (multiclass) |
| Pathology prognostics - Cervical cancer | Pathology prognostic class, cervical cancer (multiclass) |
| Pathology prognostics - Colorectal cancer | Pathology prognostic class, colorectal cancer (multiclass) |

**Table 10:** The fifteen-task register. Left: the nine binary tasks (the sub-register of the previous protocol version); right: the six multiclass tasks. Family groups the task by label source; genes give the labeled universe before outer splitting. Register definitions are fixed before inspection of operator scores.

| Task | Family | Genes | Task | Family | Genes |
| --- | --- | --- | --- | --- | --- |
| CCD Protein | complex/network | 1,388 | Blood cluster | expr. cluster | 11,881 |
| CCD Transcript | complex/network | 1,579 | Brain cluster | expr. cluster | 16,674 |
| N1 network | complex/network | 1,084 | Cell-line cluster | expr. cluster | 18,460 |
| N1 targets | complex/network | 280 | Pathology - Breast | pathology | 16,566 |
| TF vs non-TF | TF/chromatin | 2,652 | Pathology - Cervical | pathology | 16,449 |
| bivalent vs non-methyl. | TF/chromatin | 127 | Pathology - Colorectal | pathology | 16,156 |
| bivalent vs lys4-only | TF/chromatin | 164 |  |  |  |
| dosage-sens. vs insens. TF | TF/chromatin | 474 |  |  |  |
| long vs short range TF | TF/chromatin | 168 |  |  |  |

#### Bootstrap algorithm (recomputable)

Intervals are produced as follows, with a fixed generator seed. (1) Average each task’s metric over seeds and outer folds first, yielding fifteen per-task values per operator and condition on the full register (nine on the binary sub-register). (2) Resample the register tasks with replacement (10,000 iterations, task-cluster resampling; seeds and folds are never resampled). (3) Recompute the task-equal mean per iteration. (4) Report the 2.5th/97.5th percentiles. Fold- and seed-level variance is therefore absorbed into the task means before resampling; the interval measures register heterogeneity only.

### 10 Complete Per-Task Results

Tables 11–14 report Macro-F1 for every task and every primary operator in each condition on the fifteen-task register. Bold marks the largest value within a row. Table 15 retains the Core+K+X per-task values of the two attention operators on the binary sub-register, the only task set on which they were run. Table 17 reports the task-family stratification cited in Section 4.2, and Table 18 reports the semantic-leakage control cited in Section 4.2.

#### Per-task winners

Table 16 lists the best-scoring operator for each task under each condition. Thirteen of the fifteen tasks switch winners as the condition changes; only N1 network and TF vs non-TF keep a uniform winner (PCA then concat) across all four conditions. The spread column shows that only some switches are consequential (CCD Transcript, 0.146), while others (Pathology prognostics - Cervical cancer, 0.001) are within run-to-run noise.

**Table 11:** Per-task Macro-F1 under the Core condition (fifteen-task register). Rows above the midrule are the binary sub-register.

| Task | Linear concat | PCA add | PCA then concat | Interaction MLP | Low-rank higher-order | Reliability experts |
| --- | --- | --- | --- | --- | --- | --- |
| CCD Protein | 0.5204 | 0.5045 | 0.5267 | 0.5319 | 0.5223 | **0.5364** |
| CCD Transcript | 0.4145 | 0.4076 | 0.4421 | 0.5352 | 0.5335 | **0.5428** |
| N1 network | 0.6439 | 0.5939 | **0.6916** | 0.6119 | 0.5994 | 0.6178 |
| N1 targets | 0.4727 | 0.4430 | 0.5919 | 0.6697 | **0.6727** | 0.6655 |
| TF vs non-TF | 0.7608 | 0.6417 | **0.8307** | 0.7753 | 0.7732 | 0.7934 |
| bivalent vs non-methylated | 0.5126 | 0.4224 | 0.4417 | **0.8185** | 0.7974 | 0.8079 |
| bivalent vs lys4-only | 0.8084 | 0.6556 | 0.8482 | **0.8694** | 0.8552 | 0.8574 |
| dosage sensitive vs insensitive TF | 0.8545 | 0.5431 | 0.9034 | 0.9285 | **0.9295** | 0.9226 |
| long vs short range TF | 0.4281 | 0.4355 | 0.4207 | 0.4913 | 0.4737 | **0.5071** |
| Blood expression cluster | 0.0332 | 0.0276 | 0.0395 | **0.0404** | 0.0390 | 0.0401 |
| Brain expression cluster | 0.0507 | 0.0446 | 0.0565 | 0.0615 | 0.0598 | **0.0634** |
| Cell line expression cluster | 0.0556 | 0.0474 | 0.0659 | 0.0655 | 0.0666 | **0.0686** |
| Pathology prognostics - Breast cancer | 0.3276 | 0.3276 | 0.3276 | 0.3281 | **0.3316** | 0.3310 |
| Pathology prognostics - Cervical cancer | 0.3260 | 0.3260 | 0.3260 | 0.3260 | 0.3260 | **0.3264** |
| Pathology prognostics - Colorectal cancer | 0.3274 | 0.3274 | 0.3274 | 0.3619 | 0.3618 | **0.3633** |

**Table 12:** Per-task Macro-F1 under the Core+K condition (fifteen-task register). Rows above the midrule are the binary sub-register.

| Task | Linear concat | PCA add | PCA then concat | Interaction MLP | Low-rank higher-order | Reliability experts |
| --- | --- | --- | --- | --- | --- | --- |
| CCD Protein | 0.5475 | 0.5022 | **0.5540** | 0.5478 | 0.5348 | 0.5483 |
| CCD Transcript | 0.4467 | 0.4078 | 0.4929 | 0.5505 | 0.5389 | **0.5510** |
| N1 network | 0.6703 | 0.5947 | **0.7136** | 0.6368 | 0.6493 | 0.6518 |
| N1 targets | 0.5242 | 0.4608 | 0.6974 | 0.6915 | 0.6926 | **0.6995** |
| TF vs non-TF | 0.7951 | 0.6307 | **0.8649** | 0.8212 | 0.8197 | 0.8232 |
| bivalent vs non-methylated | 0.5108 | 0.4240 | 0.6445 | **0.8155** | 0.8129 | 0.8111 |
| bivalent vs lys4-only | 0.7879 | 0.5773 | 0.8733 | 0.8695 | 0.8797 | **0.8798** |
| dosage sensitive vs insensitive TF | 0.8338 | 0.4401 | 0.9136 | 0.9305 | **0.9325** | 0.9286 |
| long vs short range TF | 0.4207 | 0.4426 | 0.4281 | 0.4768 | **0.5152** | 0.4716 |
| Blood expression cluster | 0.0352 | 0.0267 | 0.0407 | 0.0447 | 0.0439 | **0.0455** |
| Brain expression cluster | 0.0550 | 0.0417 | 0.0624 | 0.0665 | 0.0659 | **0.0671** |
| Cell line expression cluster | 0.0602 | 0.0469 | 0.0714 | **0.0735** | 0.0716 | 0.0731 |
| Pathology prognostics - Breast cancer | 0.3276 | 0.3276 | 0.3276 | 0.3287 | 0.3287 | **0.3297** |
| Pathology prognostics - Cervical cancer | 0.3260 | 0.3260 | 0.3260 | 0.3260 | 0.3260 | **0.3268** |
| Pathology prognostics - Colorectal cancer | 0.3274 | 0.3274 | 0.3274 | 0.3645 | 0.3528 | **0.3669** |

#### Task-family stratification

Table 17 reports, per family and condition, the top-scoring operator and its task-equal Macro-F1. The condition response differs qualitatively across families (Section 4.2): the expression-cluster family responds almost exclusively to the X view, the complex/network family responds strongly to K, the TF/chromatin family is led by the learned experts throughout, and the pathology family remains at the constant-prediction floor for every operator.

**Table 13:** Per-task Macro-F1 under the Core+X condition (fifteen-task register). Rows above the midrule are the binary sub-register.

| Task | Linear concat | PCA add | PCA then concat | Interaction MLP | Low-rank higher-order | Reliability experts |
| --- | --- | --- | --- | --- | --- | --- |
| CCD Protein | 0.5396 | 0.5070 | **0.5470** | 0.5272 | 0.5395 | 0.5361 |
| CCD Transcript | 0.5113 | 0.4150 | **0.6824** | 0.5634 | 0.5690 | 0.5572 |
| N1 network | 0.6834 | 0.5934 | **0.7670** | 0.6299 | 0.6413 | 0.6625 |
| N1 targets | 0.5086 | 0.4328 | 0.6281 | 0.6474 | **0.6599** | 0.6587 |
| TF vs non-TF | 0.7714 | 0.6309 | **0.8314** | 0.7749 | 0.7844 | 0.7959 |
| bivalent vs non-methylated | 0.5544 | 0.4915 | 0.6290 | 0.8143 | **0.8203** | 0.8134 |
| bivalent vs lys4-only | 0.8208 | 0.6822 | 0.8630 | 0.8675 | **0.8692** | 0.8654 |
| dosage sensitive vs insensitive TF | 0.8576 | 0.4963 | 0.9195 | **0.9366** | 0.9318 | 0.9294 |
| long vs short range TF | 0.4347 | 0.4200 | 0.4347 | **0.5246** | 0.5044 | 0.5063 |
| Blood expression cluster | 0.0381 | 0.0265 | **0.0648** | 0.0456 | 0.0451 | 0.0443 |
| Brain expression cluster | 0.0604 | 0.0452 | **0.1277** | 0.0723 | 0.0726 | 0.0702 |
| Cell line expression cluster | 0.0588 | 0.0475 | **0.1020** | 0.0733 | 0.0721 | 0.0724 |
| Pathology prognostics - Breast cancer | 0.3276 | 0.3276 | 0.3276 | 0.3310 | 0.3287 | **0.3331** |
| Pathology prognostics - Cervical cancer | 0.3260 | 0.3260 | 0.3260 | 0.3263 | **0.3269** | 0.3260 |
| Pathology prognostics - Colorectal cancer | 0.3274 | 0.3274 | 0.3274 | **0.3726** | 0.3718 | 0.3665 |

**Table 14:** Per-task Macro-F1 under the Core+K+X condition (fifteen-task register, six primary operators). Rows above the midrule are the binary sub-register.

| Task | Linear concat | PCA add | PCA then concat | Interaction MLP | Low-rank higher-order | Reliability experts |
| --- | --- | --- | --- | --- | --- | --- |
| CCD Protein | 0.5564 | 0.4941 | **0.5612** | 0.5427 | 0.5423 | 0.5521 |
| CCD Transcript | 0.5307 | 0.4105 | **0.6886** | 0.5750 | 0.5813 | 0.5749 |
| N1 network | 0.6928 | 0.5743 | **0.7663** | 0.6722 | 0.6732 | 0.6891 |
| N1 targets | 0.5361 | 0.4669 | **0.7084** | 0.6790 | 0.6868 | 0.6842 |
| TF vs non-TF | 0.7875 | 0.6286 | **0.8675** | 0.8189 | 0.8223 | 0.8325 |
| bivalent vs non-methylated | 0.5607 | 0.4667 | 0.6923 | 0.8118 | 0.8151 | **0.8187** |
| bivalent vs lys4-only | 0.8009 | 0.6513 | 0.8810 | 0.8736 | **0.8919** | 0.8817 |
| dosage sensitive vs insensitive TF | 0.8433 | 0.4380 | 0.9209 | **0.9389** | 0.9302 | 0.9337 |
| long vs short range TF | 0.4355 | 0.4207 | 0.4281 | **0.5319** | 0.5150 | 0.5088 |
| Blood expression cluster | 0.0348 | 0.0285 | **0.0604** | 0.0484 | 0.0499 | 0.0486 |
| Brain expression cluster | 0.0573 | 0.0428 | **0.1249** | 0.0755 | 0.0755 | 0.0745 |
| Cell line expression cluster | 0.0619 | 0.0449 | **0.1020** | 0.0785 | 0.0775 | 0.0771 |
| Pathology prognostics - Breast cancer | 0.3276 | 0.3276 | 0.3276 | 0.3325 | 0.3292 | **0.3352** |
| Pathology prognostics - Cervical cancer | 0.3260 | 0.3260 | 0.3260 | 0.3264 | 0.3260 | **0.3273** |
| Pathology prognostics - Colorectal cancer | 0.3274 | 0.3274 | 0.3274 | 0.3633 | 0.3666 | **0.3742** |

#### Semantic-leakage control

Table 18 reports the matched condition contrasts on the full register and on the ten-task register that removes every task sharing curation resources with the K view (Section 4.2). Every qualitative conclusion is unchanged under the control.

**Table 15:**
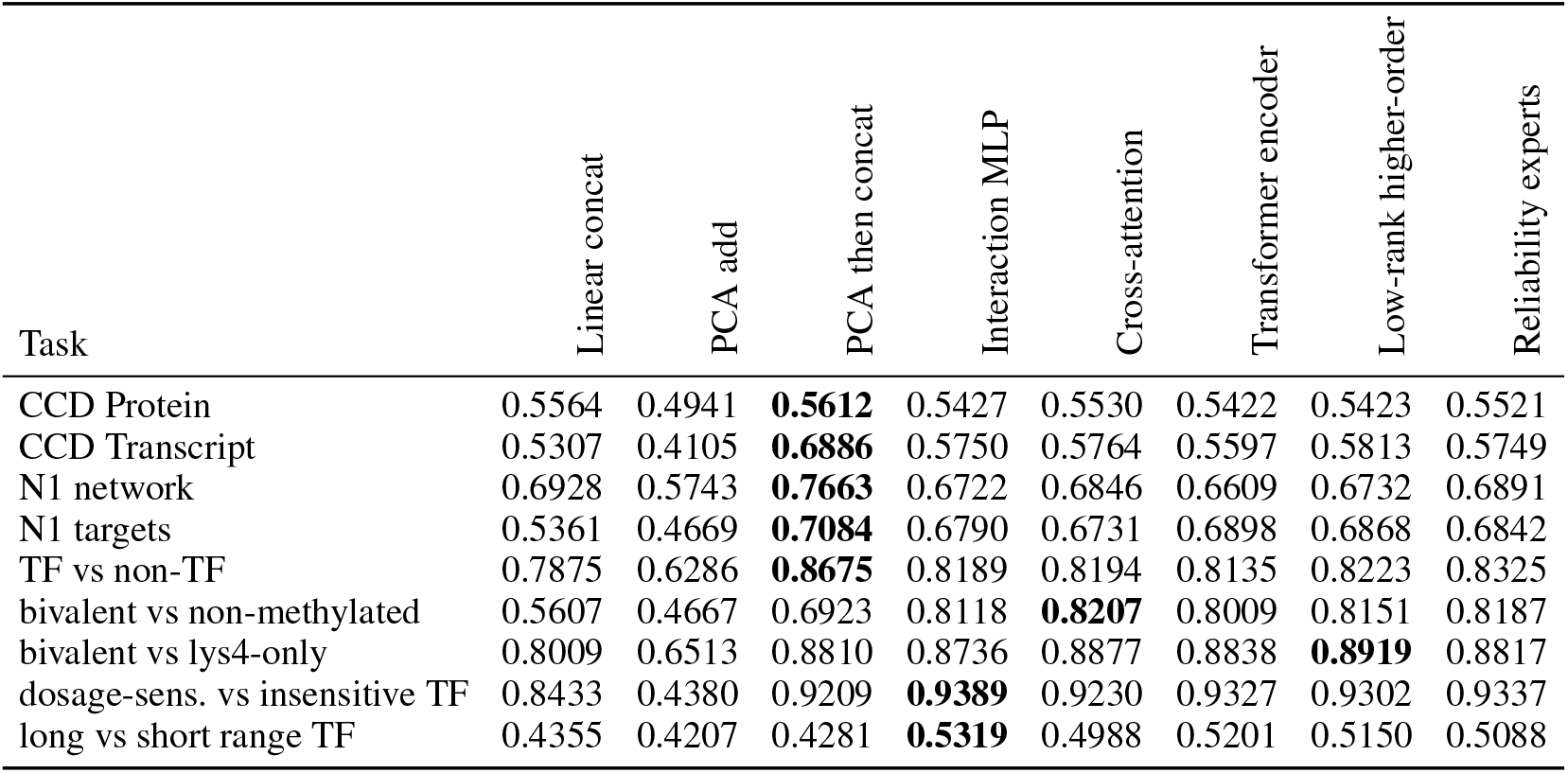
Per-task Macro-F1 under the Core+K+X condition on the binary sub-register, including the two attention operators (nine tasks, eight operators).

**Table 16:** Best-scoring operator per task under each protocol-defined condition, with the spread of the winning score across conditions. Winners are selected exclusively among the six primary operators evaluated under all four conditions; the two attention operators (Core+K+X only, Table 15) are not part of this summary. LC: Linear concat; PA: PCA add; PC: PCA then concat; INT: Interaction MLP; HO: Low-rank higher-order; REL: Reliability experts. Thirteen of fifteen tasks switch winners across conditions; small spreads mean the operator choice matters less than the task.

| Task | Core | Core+K | Core+X | Core+K+X | Spread |
| --- | --- | --- | --- | --- | --- |
| Blood expression cluster | INT | REL | PC | PC | 0.024 |
| Brain expression cluster | REL | REL | PC | PC | 0.064 |
| CCD Protein | REL | PC | PC | PC | 0.025 |
| CCD Transcript | REL | REL | PC | PC | 0.146 |
| Cell line expression cluster | REL | INT | PC | PC | 0.033 |
| N1 network | PC | PC | PC | PC | 0.075 |
| N1 targets | HO | REL | HO | PC | 0.048 |
| Pathology prognostics - Breast cancer | HO | REL | REL | REL | 0.005 |
| Pathology prognostics - Cervical cancer | REL | REL | HO | REL | 0.001 |
| Pathology prognostics - Colorectal cancer | REL | REL | INT | REL | 0.011 |
| TF vs non-TF | PC | PC | PC | PC | 0.037 |
| bivalent vs lys4-only | INT | REL | HO | HO | 0.023 |
| bivalent vs non-methylated | INT | INT | HO | REL | 0.005 |
| dosage sensitive vs insensitive TF | HO | HO | INT | INT | 0.009 |
| long vs short range TF | REL | HO | INT | INT | 0.025 |

#### Binary sub-register condition effects

For continuity with the previous protocol version, Table 19 reports the matched condition contrasts on the nine-task binary sub-register; the ordering matches the full register with larger magnitudes.

### 11 Attention-Based Operators: Definitions, Results, and a Granularity Mismatch under a Label-Free Contract

This appendix collects the two additional attention operators separately from the six primary operators. The operators are defined first, their results are reported under the identical contract, and the granularity mismatch is then discussed without treating it as an isolated causal explanation.

**Table 17:** Task-family stratification: per-family top operator and task-equal Macro-F1 under each condition (LC/PA/PC/INT/HO/REL abbreviations as in Table 16).

| Family | Core |  | Core+K |  | Core+X |  | Core+K+X |  |
| --- | --- | --- | --- | --- | --- | --- | --- | --- |
|  | op | F1 | op | F1 | op | F1 | op | F1 |
| complex/network (4 binary) | REL | 0.5906 | PC | 0.6145 | PC | 0.6561 | PC | 0.6811 |
| TF/chromatin (5 binary) | REL | 0.7777 | HO | 0.7920 | INT | 0.7836 | REL | 0.7951 |
| expression cluster (3 multiclass) | REL | 0.0574 | REL | 0.0619 | PC | 0.0982 | PC | 0.0958 |
| pathology (3 multiclass) | REL | 0.3403 | REL | 0.3412 | INT | 0.3433 | REL | 0.3455 |

**Table 18:** Semantic-leakage control: condition contrasts with and without the five tasks whose labels share curation resources with the K (text) view.

| Method | $\Delta_{KX}$ (full) | $\Delta_{KX}$ (no overlap) | $N_{KX}$ (full) | $N_{KX}$ (no overlap) |
| --- | --- | --- | --- | --- |
| PCA then concat | +0.0629 | +0.0600 | −0.0195 | −0.0214 |
| Linear concat | +0.0228 | +0.0168 | −0.0049 | −0.0047 |
| PCA add | −0.0067 | +0.0041 | +0.0034 | +0.0019 |
| Interaction MLP | +0.0169 | +0.0108 | +0.0022 | +0.0016 |
| Low-rank higher-order | +0.0227 | +0.0182 | −0.0051 | −0.0030 |
| Reliability experts | +0.0179 | +0.0169 | +0.0030 | −0.0012 |

#### 11.1 Definitions under the shared contract

##### Cross-attention over views

Stages 1–3 treat the view stack as an unordered feature vector or as a set of branch outputs. The cross-attention operator instead treats each view as a query source over the others, so that the fused representation depends on which view is being read at each step. For view *v* ∈ *V*_*c*_, let 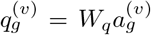 and let the keys and values be formed from the remaining views *u*≠ *v*:

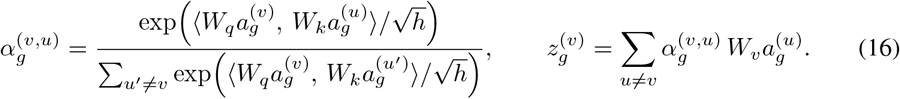

The operator output mean-pools the per-view reads with a residual connection and a shared feed-forward map,

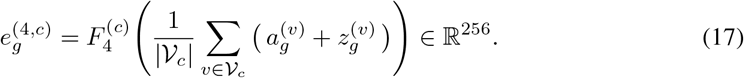

The attention operators use the same adapted view coordinates and output budget as the learned experts, so the comparison holds the available view information and export dimensionality fixed. It does not isolate fusion bias from operator-specific capacity or the shared training budget; those asymmetries remain part of the stated contract limitations.

##### Transformer encoder over the view stack

The transformer encoder processes the view stack as a sequence. Let 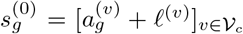 where *ℓ*^(*v*)^ is a learnable view-position embedding under a fixed view ordering (with |V_*c*_ | ≤ 5, these embeddings act as view identifiers rather than sequence positions). The stack is updated by *L* self-attention blocks in the standard pre-norm form with both residual connections,

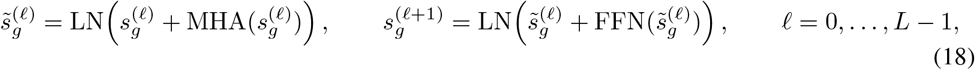

and the representation is the mean-pooled output followed by a projection,

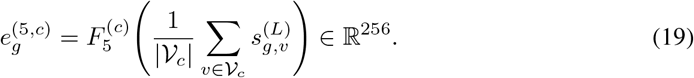

Both operators use the same adapters, the same 256-dimensional output, the same outer-training-only fitting boundary, the same label-free objective, and the same probe as every other operator. For the transformer encoder, the block count *L*, the attention-head configuration, and the feed-forward width were not frozen into the released configuration snapshot, so its exact learned-parameter count cannot be reconstructed from the released artifacts; we record this as a reproducibility gap (Appendix 21) rather than reconstructing a count post hoc, and the operator is therefore excluded from any capacity-matched comparison.

**Table 19:** Matched condition deltas relative to Core on the binary sub-register (nine tasks), computed identically to the full-register table in the main text.

| Method | $\Delta_K$ | $\Delta_X$ | $\Delta_{KX}$ | $N_{KX}$ |
| --- | --- | --- | --- | --- |
| PCA then concat | +0.0539 | +0.0672 | +0.0908 | −0.0303 |
| Linear concat | +0.0135 | +0.0296 | +0.0365 | −0.0066 |
| PCA add | −0.0186 | +0.0024 | −0.0107 | +0.0055 |
| Interaction MLP | +0.0120 | +0.0060 | +0.0236 | +0.0055 |
| Low-rank higher-order | +0.0243 | +0.0181 | +0.0335 | −0.0089 |
| Reliability experts | +0.0127 | +0.0082 | +0.0250 | +0.0041 |

#### 11.2 Results under the identical contract

Table 20 summarizes the two operators on all three evaluation tracks on which they were run. Internally, both sit below the PCA-then-concat baseline (0.7238) and Reliability experts (0.7195). Externally, both sit below all three fixed RepGene experts and the PCA-then-concat baseline. On the cell-level boundary diagnostic of Section 4.4, Cross-attention (0.9026) and the transformer encoder (0.8956) bracket the dimension-matched random control (0.8987), i.e., neither separates from it. Per-task Core+K+X scores for both operators appear in Table 14. The two operators were evaluated under Core+K+X only (9 × 2 × 3 × 3 = 162 task–operator–seed–fold records), so their condition-response profile is not estimated; this coverage restriction is recorded wherever their absence could otherwise be misread.

**Table 20:** Attention-based operators across evaluation tracks (task-equal Macro-F1). Internal and external entries carry descriptive 95% task-bootstrap intervals; cell-level entries carry seed standard deviations over five seeds. Reference rows reproduce the strongest non-attention entries for calibration.

| Operator | Internal Core+K+X | External Core+K+X | Cell level |
| --- | --- | --- | --- |
| Cross-attention | 0.7152 [0.6212, 0.8062] | 0.7177 [0.6228, 0.8095] | 0.9026 ± 0.0145 |
| Transformer encoder | 0.7115 [0.6192, 0.8023] | 0.7126 [0.6164, 0.8062] | 0.8956 ± 0.0149 |
| *Reference* |  |  |  |
| PCA then concat | 0.7238 [0.6224, 0.8157] | 0.7221 [0.6209, 0.8140] | 0.9091 ± 0.0104 |
| Reliability experts | 0.7195 [0.6256, 0.8107] | 0.7227 [0.6250, 0.8159] | 0.8889 ± 0.0246 |
| Low-rank higher-order | 0.7176 [0.6242, 0.8081] | 0.7251 [0.6335, 0.8143] | 0.9024 ± 0.0129 |
| Random 256-D control | – | 0.4240 [0.4071, 0.4457] | 0.8987 ± 0.0100 |

#### 11.3 Why the mismatch is structural: one gene, one vector per view

The negative result above admits a simple structural explanation, which we put forward as a hypothesis consistent with the evidence rather than an established causal claim. In this task family, every modality supplies *exactly one vector per gene*: DNA is distilled to one sequence-derived feature vector, RNA to one sequence-derived vector, protein to one protein language-model vector, knowledge to one text vector, and single-cell expression to one summary vector. At no point do we obtain embeddings of the individual tokens *inside* a gene—there are no per-base embeddings along the DNA sequence, no per-residue embeddings along the protein, and no per-transcript embeddings. Token-wise attention mechanisms, however, derive their expressive advantage from computing interactions *between many fine-grained tokens*: a transformer encoder earns its keep on sequences of hundreds to thousands of tokens, and cross-attention earns its keep by reading one token stream against another.

Under the matched contract, the “sequence” available to these operators has length |*V*_*c*_ | ≤5, and each “token” is an entire view vector. Two consequences follow. First, there is no fine-grained interaction surface to exploit: every coordinate of a view vector is already a global summary of that view, so attending between whole-view vectors cannot recover anything finer than a data-dependent weighted mixture of at most five vectors—a function class already covered by the reliability-conditioned weighting of Stage 3 and by the explicit pairwise features of Stage 1. Second, the operators pay for the capacity they cannot use: query/key/value projections, position embeddings, and multi-layer feed-forward blocks add parameters and estimation variance under an eight-epoch label-free budget, without any token-level signal to amortize them. Under this contract’s label-free objective and fixed eight-epoch budget—8–16 gradient updates per representation fit at batch size 1,024, given the registered tasks’ outer-training partitions of 84–1,768 genes—attention is therefore unsuited to this granularity; the argument is structural, and the budget extension of Appendix 28.2 bounds the optimization confound for cross-attention—it improves across budgets (0.6936 → 0.7198 → 0.7318) yet never attains an advantage—but granularity cannot be fully separated from capacity confounds (Section 5), because the transformer encoder cannot be re-executed at higher budgets. The measured ordering in Table 20 is exactly what this argument predicts: both operators land in the middle of the operators that mix whole-view vectors, below those whose bias is explicit about whole-view interactions, and they never separate from the random control at cell level.

We stress the boundary of this claim. The argument concerns *this* data granularity: given encoders that expose per-base, per-residue, or otherwise token-level embeddings (e.g., nucleotide-level language-model hidden states), attention would again have a token sequence to exploit, and the same matched contract could re-evaluate it fairly. What the present evidence rules out is the default assumption that attention is the obviously correct fusion operator for one-vector-per-gene multi-view representations.

### 12 External Benchmark Details

The external RF track evaluates the same nine registered binary tasks in the Core+K+X state with 256-D representations, three seeds, and three folds. Its probe uses 200 estimators, maximum depth 12, minimum leaf size 2, and `balanced_subsample` class weighting. This configuration was frozen when the external track was designed, before the internal condition results were inspected, and was never revised afterwards (Appendix 18); the external track is thus an independently configured replication, not a re-fit on seen results. Compared entries are: a random 256-D control, GenePT as a text-only source baseline, a PRISME+scRNA five-view protocol extension (Zheng et al., 2025) (not the unmodified original PRISME method), the three baselines, and the five RepGene operators. The native PRISME autoencoder is not an entry of the primary contract: it integrates its own modality set under its own reconstruction objective, whereas the contract requires each entry to export 256-D representations for the registered universe under the shared label-free objective and frozen probe; adapting native PRISME into the primary grid is an open extension (Appendix 21). Task-equal Macro-F1 is the primary external metric.

### 13 Gene-TO-Cell Aggregation and Controls

For cell *c* and shared gene set *G*_*D*_, expression-weighted aggregation is

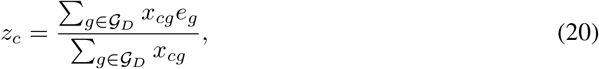

where *x*_*cg*_ is the aorta cell-by-gene expression value used as the aggregation weight and *e*_*g*_ is a gene representation. This downstream aggregation is distinct from the X view, which is a single-cell expression input to the gene-level fusion contract. Equation 20 introduces at least four additional design choices that are absent from the gene-level estimand: the aggregation operator, the expression weighting and normalization, the cell annotation granularity, and the cell-level classifier. Each can change the ranking independently of the fusion operator. Required controls are an expression-only reference, a projection-only reference, a dimension-matched random control, and a label permutation. With those controls in place (Section 4.4), the fused representations do not separate from a random 256-D control, and the expression-only reference leads. We therefore report cell-level results as a boundary diagnostic.

**Table 21:** Secondary metrics on the fifteen-task register: task-equal balanced accuracy (BAcc) and ROC-AUC under the four protocol-defined conditions. Macro-F1 remains the prespecified primary metric; the AUC ordering inverts the primary ordering in favour of PCA then concat.

| Method | Core |  | Core+K |  | Core+X |  | Core+K+X |  |
| --- | --- | --- | --- | --- | --- | --- | --- | --- |
|  | BAcc | AUC | BAcc | AUC | BAcc | AUC | BAcc | AUC |
| Reliability experts | 0.5046 | 0.7170 | 0.5138 | 0.7310 | 0.5116 | 0.7311 | 0.5224 | 0.7441 |
| Interaction MLP | 0.5029 | 0.7157 | 0.5115 | 0.7328 | 0.5080 | 0.7274 | 0.5184 | 0.7462 |
| Low-rank higher-order | 0.4988 | 0.7116 | 0.5127 | 0.7350 | 0.5104 | 0.7325 | 0.5196 | 0.7467 |
| PCA then concat | 0.4766 | 0.7299 | 0.5013 | 0.7492 | 0.5150 | 0.7823 | 0.5269 | 0.7894 |
| Linear concat | 0.4563 | 0.7035 | 0.4619 | 0.7186 | 0.4691 | 0.7234 | 0.4707 | 0.7365 |
| PCA add | 0.4133 | 0.6671 | 0.4061 | 0.6631 | 0.4141 | 0.6666 | 0.4092 | 0.6648 |

#### Derivation of the dilution bound (Eq. 6)

Write *p*_*c*_ (*g*) = *x*_*cg*_ */*∑_*j*_*x*_*cj*_, so that *p*_c_ (*g*) ≥ 0 and ∑_*g*_ *p*_*c*_(*g*) = 1, and decompose the gene representation as *e*_*g*_ = *µ* + *b*_*t*(*g*)_ + *ε*_*g*_ with a global mean *µ*, a cell-category term *b*_*t*(*g*)_, and a fusion-specific residual *ε*_*g*_. Linearity of the weighted average gives the decomposition in Eq. 6 exactly.

##### Deterministic part

By the triangle inequality and ∑_*g*_ *p*_*c*_(*g*) = 1,

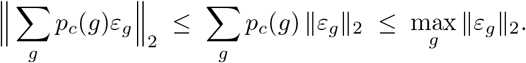

No 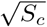 attenuation is available deterministically: if the residuals are perfectly collinear, *ε*_*g*_ = **u**≠ 0 for all *g*, then ∥ ∑_*g*_*p*_*c*_ (*g*)**u**∥_2_ = ∥**u**∥_2_ while 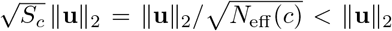 for any non-degenerate weighting (for uniform weights over |*G*| genes the violation factor is exactly 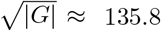 at |*G*| = 18,460). A deterministic 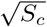 bound would require exact pairwise orthogonality ⟨*ε*_*g*_, *ε*_*g*_*′* ⟩ = 0 for all *g*≠ *g*^*′*^, which cannot hold for 18,460 nonzero vectors in ℝ^256^.

*Statistical part*. Assume instead that the residuals are zero-mean and mutually uncorrelated across genes: E[*ε*_*g*_] = 0 and 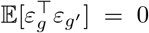 for *g* ≠ *g*^*′*^. Expanding the squared norm and applying uncorrelatedness,

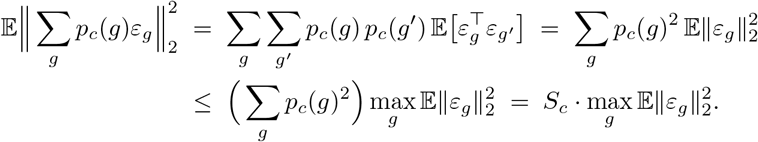

Taking square roots yields Eq. 6: the root-mean-square norm of the fusion-specific component is bounded by 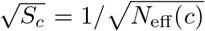 times the largest per-gene residual root-mean-square norm, while the category-barycenter term ∑_*g*_ *p*_*c*_(*g*)*b*_*t*(*g*)_ passes through the aggregation unchanged. The uncorrelatedness premise is a modeling assumption about residual structure, not a theorem about arbitrary representations; the observed attenuation toward the dimension-matched random control (Section 4.4) is the empirical content, and the bound is the quantitative bridge that predicts its order of magnitude.

### 14 Secondary Metrics in Full

Macro-F1 is the prespecified primary metric. Table 21 reports task-equal balanced accuracy and ROC-AUC for the six operators under all four conditions on the fifteen-task register. The operator ordering is *not* preserved across metrics: under ROC-AUC, PCA then concat is numerically highest in all four conditions (0.7299, 0.7492, 0.7823, 0.7894), ahead of every learned expert, while under balanced accuracy the learned experts lead throughout. The binary sub-register shows the same inversion (AUC for PCA then concat: Core 0.7905, Core+K 0.8111, Core+X 0.8231, Core+K+X 0.8347). Section 4.3 discusses the threshold-sensitivity interpretation of this divergence. We therefore state plainly that the register-level conclusions are metric-dependent, that Macro-F1 was prespecified, and that no superiority claim under AUC is made.

**Table 22:** Evidence boundary. The left column lists statements that the current result package does *not* license; the right column gives the replacement wording used throughout the paper.

| Not licensed | Required wording |
| --- | --- |
| “Operator $A$ beats operator $B$” | “$A$ attains a larger task-equal mean in condition $c$; intervals overlap” |
| “K and X views cause the improvement” | “Changing the protocol-defined input state from Core to Core+K+X is associated with a mean change of $\Delta$” |
| “K and X interact synergistically” | “The performance decomposition $N_{KX}$ is negative/positive” |
| “The method is robust to missing views” | “The method was evaluated under controlled protocol-defined conditions” |
| “Good gene representations give good cell representations” | “Gene-level and cell-level utility are separate estimands; the transfer is unmeasured” |
| “Dynamic routing selects the best expert” | “Dynamic routing is prespecified and unvalidated” |
| “The 95% interval shows significance” | “The interval is a descriptive task-bootstrap summary of register heterogeneity” |
| “The ordering is metric-robust” | “BACC preserves the leading trio; AUC numerically favours PCA then concat in all four conditions” |
| “The knowledge view is leak-free” | “The knowledge view shares curation resources with task-label families; a semantic-leakage path is open, and a registered removal control bounds its influence on the reported contrasts (Section 4.2)” |
| “The fold/seed layer supports the swap” | “Task-equal aggregation absorbs fold/seed variance before the task bootstrap; swaps are not verified per fold or seed” |

### 15 Claim–Evidence Ledger

The claim–evidence ledger is the paper’s wording-control layer: every register-level statement in the main text is required to use the replacement wording in Table 22, and statements in the left column are explicitly disclaimed. The machine-readable form of this ledger is maintained as `claim_evidence_ledger.csv` in the released artifact bundle; Table 22 reproduces its current contents.

### 16 Artifacts and Reproduction Order

The released bundle contains: (i) immutable resolved configuration files; (ii) the per-task, per-condition, per-method summary tables; (iii) paired condition-effect tables; (iv) the external track statistics; (v) the figure-generation script with its source manifest and SHA-256 checksums; and (vi) this ledger. Reproduction order is: build the aligned universe → fit adapters per outer fold → fit each fusion operator → export 256-D representations → run the probe → aggregate task-equal → regenerate figures from the manifest. Every figure can be regenerated from `source_data/` without re-running any model.

### 17 Figure-TO-Claim Map

Table 23 is the audit trail between the released artifacts and the statements in the main text: every figure is listed with the claim it supports, its source table, and the reason it does not support a stronger claim. The machine-readable form is maintained as `figure_to_claim_map.json` in the released artifact bundle.

### 18 Protocol Design Decisions

The last two rows deserve emphasis. Reporting the attention operators required adding a coverage limitation to the paper; suppressing them would have produced a cleaner story and a weaker one.

**Table 23:**
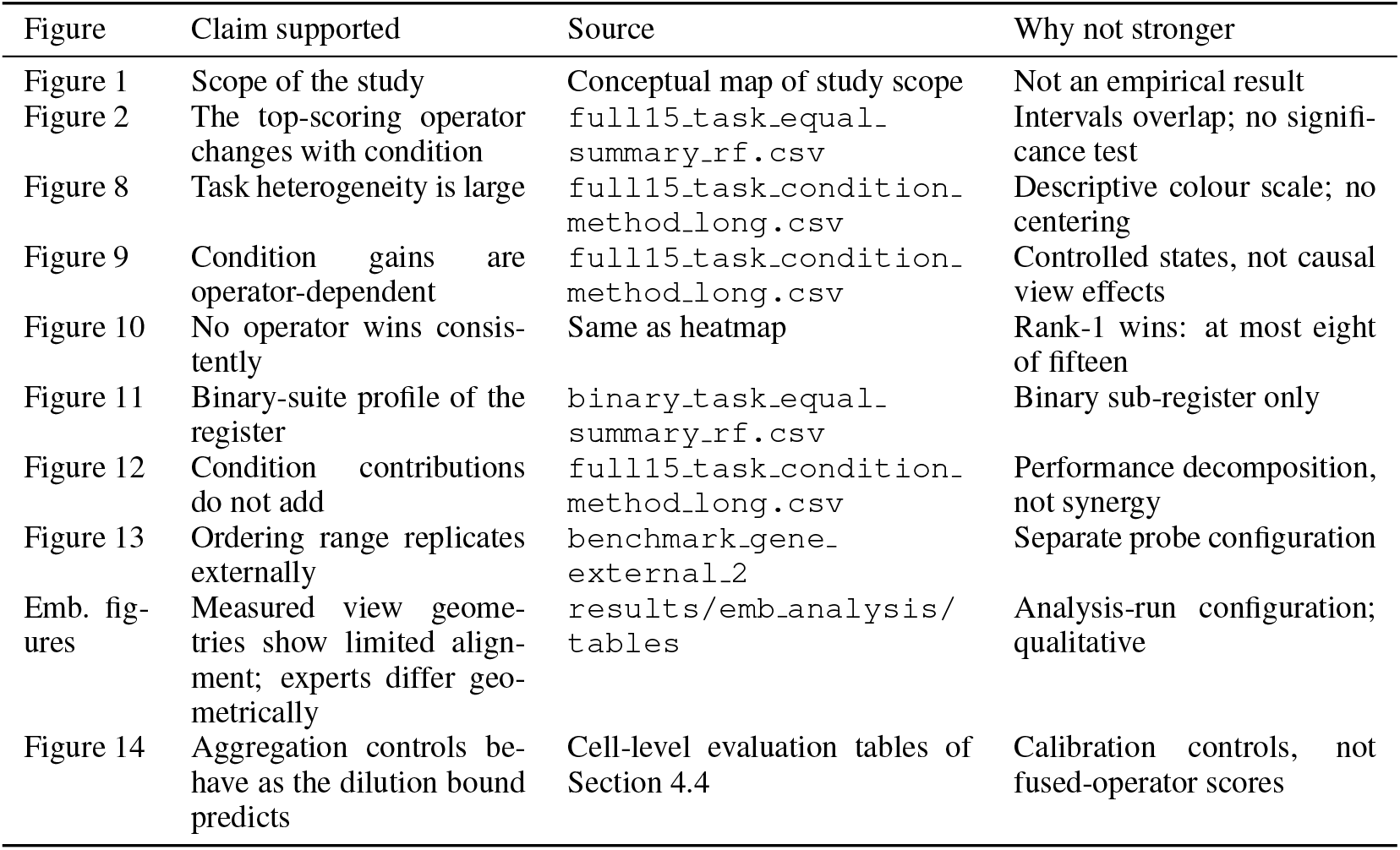
Every figure with the claim it supports, the source table, and the reason it does not support a stronger claim.

Keeping the external probe separate means the external track is a genuine replication of the ordering rather than a re-measurement of the same configuration.

### 19 Protocol Card

#### RepGene fusion-bias contract: protocol card

**Entry requirements.** (i) Export a 256-D representation for the same 18,460-gene universe, per condition, using adapters fitted on outer-training features only. (ii) Re-run the frozen probe (internal: RF, 300 trees, depth 16, leaf 2; external: RF, 200 trees, depth 12, balanced subsample). (iii) Report task-equal Macro-F1 with the descriptive task-bootstrap of Appendix 9.

**Prohibited.** Re-tuning the probe, folds, seeds, metric, or aggregation after seeing scores; reporting the full-input column alone (it inverts the condition-response conclusion); reading intervals as significance tests; interpreting view weights as gene importance or assay quality.

**Registered extensions (separate tracks, reported in the appendix).** The probe family, the training budget, the capacity-matched pairing control, and the calibration/threshold treatment are reported as registered tracks evaluated on identical exports (Appendix 28); they document the sensitivity of the primary grid and do not modify the entry requirements above.

**Known asymmetries (accepted on entry).** The trained adapters are shared by the learned experts while the three baselines operate on fold-standardized raw views, and the fixed eight-epoch label-free budget favours parameter-light operators; baselines carry no learnable interaction terms while experts do (Table 8); the register is fifteen tasks across four correlated annotation families; intervals describe the register, not a task population.

**Scope of conclusions.** Protocol-defined conditions only—not natural missingness, not deployment robustness, not causal view effects, not cell-level utility. Attention operators are covered under Core+K+X only.

**Adding a task** requires recomputing *every* operator, because task-equal aggregation changes with the register.

**Table 24:** Design decisions, the alternative that was rejected, and the reason. Recording the rejected alternative is what distinguishes a protocol from a set of convenient choices.

| Decision | Rejected alternative | Reason |
| --- | --- | --- |
| Fixed 256-D export | Per-method best dimension | Dimension alone changes the probe’s bias–variance point |
| One probe for all operators | Per-operator probe tuning | Tuning couples the probe to the operator and destroys matching |
| Task-equal aggregation | Gene-weighted or fold-weighted | Large tasks would dominate the register |
| Outer-training-only fitting | Fit on all features | Leaks test-fold statistics into the representation |
| Descriptive bootstrap intervals | Paired significance tests | Fifteen tasks are a register, not a population sample |
| Four controlled conditions | Natural missingness simulation | Natural availability is unobserved here and answers a different estimand |
| Report attention results despite being negative | Drop the two operators | A negative result under a matched contract is informative |
| Separate external probe configuration | Reuse the internal probe | Would be tuned on the internal register |
| Freeze external probe before comparing tracks | Iterate probe design after seeing primary scores | Independence of the comparison would be destroyed |
| Report AUC inversion (Appendix 14) | Report only the primary metric | The register picture is metric-dependent; hiding the inversion would overstate robustness |
| Report the completed fifteen-task register | Wait for the full 54-task eligibility set | Completion status, not result-dependent selection, fixes the register |
| Semantic-leakage removal control (Section 4.2) | Leave the K-view leakage path unquantified | A registered removal control bounds the pathway without new training |

**Table 25:**
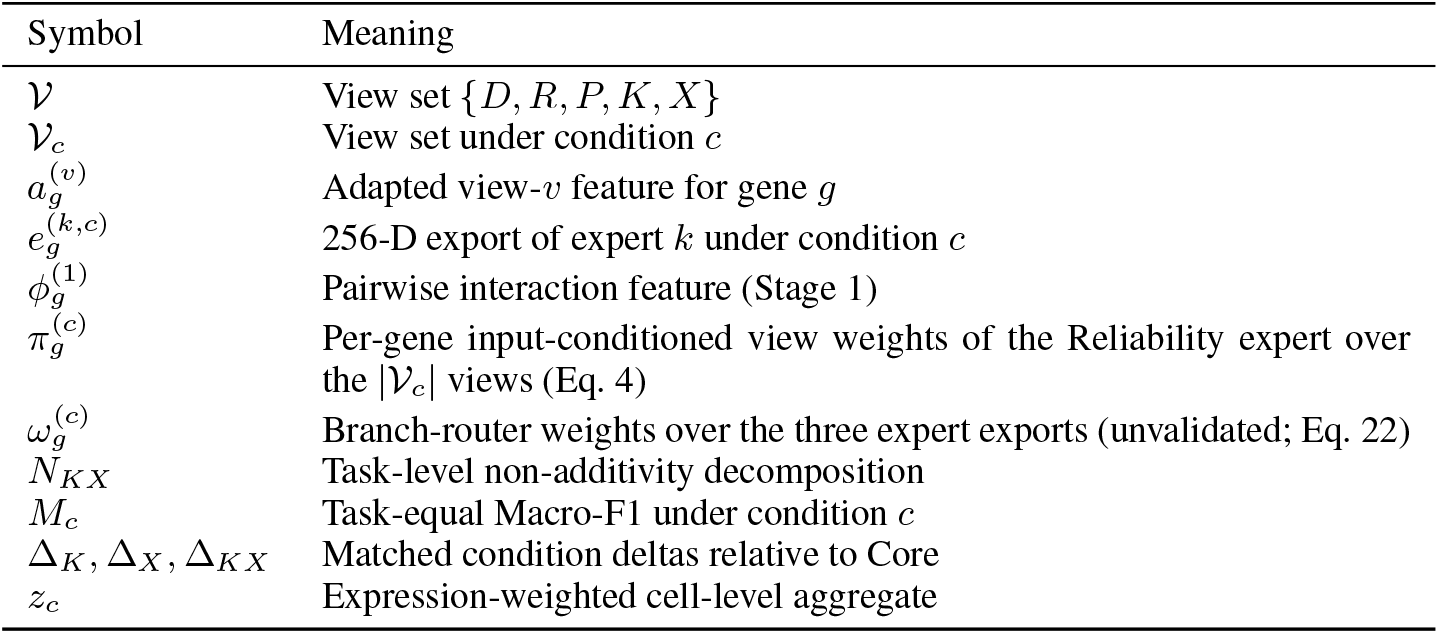
Symbols used in the paper. Gene-level vectors are set in italics throughout; no separate boldface convention is used.

| Symbol | Meaning |
| --- | --- |
| $\mathcal{V}$ | View set $\{D, R, P, K, X\}$ |
| $\mathcal{V}_c$ | View set under condition $c$ |
| $a_g^{(v)}$ | Adapted view-$v$ feature for gene $g$ |
| $e_g^{(k,c)}$ | 256-D export of expert $k$ under condition $c$ |
| $\phi_g^{(1)}$ | Pairwise interaction feature (Stage 1) |
| $\pi_g^{(c)}$ | Per-gene input-conditioned view weights of the Reliability expert over the $\vert \mathcal{V}_c\vert $ views (Eq. 4) |
| $\omega_g^{(c)}$ | Branch-router weights over the three expert exports (unvalidated; Eq. 22) |
| $N_{KX}$ | Task-level non-additivity decomposition |
| $M_c$ | Task-equal Macro-F1 under condition $c$ |
| $\Delta_K, \Delta_X, \Delta_{KX}$ | Matched condition deltas relative to Core |
| $z_c$ | Expression-weighted cell-level aggregate |

### 20 Notation

### 21 Limitations

- **Register size and label-family correlation**. Fifteen tasks are reported (nine binary, six multiclass) of 54 eligible; the remaining 36 multiclass and 3 multilabel tasks were not run to completion and contribute no numbers. Tasks share functional-annotation families (complex/network, TF/chromatin, expression cluster, pathology), so task-equal averaging re-weights correlated label structures rather than independent evidence; the family stratification of Section 4.2 makes this composition explicit. The pathology family sits at the constant-prediction floor, so it contributes register coverage rather than condition signal. Completing the full eligibility-filtered set, and cross-checking operator behaviour against open multi-task suites (Zhong et al., 2025; Brechtmann et al., 2023), remains the primary scope extension.
- **Gene-intersection composition**. The 18,460-gene five-view intersection over-represents well-studied, well-annotated genes; conclusions do not automatically extend to poorly annotated genes, other organisms, or other view sets.
- **Text-view semantic-leakage caution**. The knowledge view embeds curated functional text that shares curation resources with several task-label families. This pathway is now bounded by a registered control (Section 4.2): removing all five tasks that share curation resources with K leaves every qualitative conclusion unchanged, and the K-condition gain is not concentrated in the overlapping tasks. The control bounds the pathway’s influence on the reported contrasts; it does not certify the K view as leak-free, and K-condition gains must not be read as mechanism. The same curation process induces a publication-density (ascertainment) bias: well-studied genes carry richer text, which can act as an artificial classification shortcut on curated functional tasks while obscuring generalization to unannotated genes (Brechtmann et al., 2023). The evidence boundary is stated in the ledger (Appendix 15).
- **External platform coverage**. The primary contract does not include a native multi-modal autoencoder entry such as PRISME (Zheng et al., 2025), whose own reconstruction objective and modality set do not satisfy the contract’s shared-objective entry requirement; the external track reports a five-view PRISME+scRNA protocol extension instead (Appendix 12), and a native entry is an open extension.
- **No bias-versus-capacity decomposition**. Baselines carry no learnable interaction terms while experts do (Table 8); a same-budget, interaction-free expert control does not exist in the primary register. The capacity-matched pairing control (Appendix 28.3) isolates the contribution of gene-wise cross-view pairing within the interaction expert, but it does not substitute for an interaction-free expert at matched budget. The supplementary sweep (Appendix 24) includes an equal-capacity MLP-on-concat control on synthetic and subset tasks, where explicit interaction terms add at most +0.009 task-equal Macro-F1; adding this control to the primary fifteen-task register is a defined extension of the contract. We record differences between function classes and deliberately do not attribute them to “bias rather than capacity”.
- **Training regime partially varied**. One label-free objective is used throughout, and the primary register runs at one eight-epoch budget: at batch size 1,024 that budget amounts to 8–16 gradient updates per representation fit (outer-training partitions of 84–1,768 genes), so the attention result and the experts’ margins are conditional on a heavily constrained optimization regime. A bounded epoch sweep at 32 and 64 epochs is reported for the four learned operators on the binary sub-register (Appendix 28.2); coefficient sensitivity (*λ*), training curves, and epoch sweeps on the multiclass tasks remain outside this package.
- **Controls partially instantiated**. Three controls are executed and reported: the semantic-leakage removal control (Section 4.2), the capacity-matched view-pairing shuffle control (Appendix 28.3), and the probe-sensitivity and calibration extensions (Appendices 28.1, 28.4). The staged P0–P4 ablation (parameter/FLOP/time/peak-memory accounting per stage) is *not* executed, peak memory is not measured, and the budget sweep does not cover the multiclass tasks; these are recorded as gaps, not promised as future deliveries within this submission.
- **Fold/seed layer absorbed**. Task-equal aggregation pools seeds and folds before the task bootstrap; condition swaps are not verified per fold or per seed, and the bootstrap interval does not contain a fold/seed variance component.
- **Multiplicity**. The 6-operator × 4-condition grid produces frequent top-score swaps under noise; *N*_*KX*_ extrema are within the same magnitude and are read descriptively (Section 4.2).
- **Metric and threshold dependence**. Under ROC-AUC the ordering inverts in favour of PCA then concat in all four conditions (Appendix 14); a registered calibration analysis (Appendix 28.4) shows this inversion is a threshold/calibration effect—PCA then concat ranks best and is calibrated worst, so the ordering flips again once thresholds are tuned— and no cross-metric or cross-threshold superiority is claimed.
- **Coverage asymmetry**. Cross-attention and the transformer encoder were evaluated in Core+K+X only, so their condition response is not estimated; Appendix 11 gives the granularity argument, with confounds stated.
- **Primary probe family**. The primary register uses random forests only (two configurations). A registered probe extension reports a frozen logistic probe and a supervised MLP-head reference evaluated on the *same* exported representations (Appendices 28.1 and 28.4): the structural findings survive, while the identity of the leading operator changes with the probe and with the decision threshold. Gradient-boosted probes, fine-tuned heads, and probe sweeps on the multiclass register remain unmeasured.
- **No homology-aware splitting**. Outer folds are stratified randomly within each task’s labeled genes; sequence homology, shared protein domains, and paralogous gene families can place near-duplicate genes on both sides of a split. The outer-training-only boundary removes statistical and transductive leakage of preprocessing and fitting, but not this biological leakage channel; homology-aware split construction is outside the current package.
- **Transformer-encoder hyperparameters not frozen**. The block count, attention-head configuration, and feed-forward width of the transformer-encoder operator were not frozen into the released configuration snapshot, so its exact parameter count is not recoverable from the released artifacts and the operator cannot be re-executed (Appendix 5); this is recorded as a gap rather than reconstructed post hoc, and the budget statement of Appendix 28.2 is consequently restricted to cross-attention.
- **Single cell-level design point**. One tissue (aorta), linear expression-weighted aggregation, one annotation granularity, logistic-regression classifier, five seeds. The dilution result is a boundary at this point, not a general cell-level law.
- **No natural missingness**. All conditions are controlled protocol states.
- **Unvalidated routing**. The branch-level router is prespecified, not tested (Appendix 25).
- **Supplementary multiplicative-interaction evidence**. The results in Appendix 24 provide summary-level values without fold-level records or interval endpoints; they are therefore interpreted as magnitude-only supplementary evidence.

### 22 Reproducibility Checklist

1. **Gene universe**. 18,460 genes present in all five views; identifier mapping and source versions recorded in the manifest; checksums published with the snapshot.
2. **Splits and seeds**. Three outer folds; seeds {41, 42, 43}; split hashes recorded. Folds are shared by every operator and every condition.
3. **Adapters**. Per-view standardization and dimensionality reduction fitted on outer-training features only, applied unchanged to validation and test folds.
4. **Operators**. Eight fusion maps; shared hidden width 96, dropout 0.1, eight epochs, batch size 1024, Adam with learning rate 10^−3^ and weight decay 10^−5^; label-free weights (0.1, 0.01, 0.05, 0.1) on top of a unit-weight reconstruction term whose decoder branch is training-only.
5. **Exports**. Every operator exports exactly 256 dimensions for every condition, including Core+K+X.
6. **Probe**. Internal: random forest, 300 trees, maximum depth 16, minimum leaf size 2, identical across operators. External: 200 trees, depth 12, balanced subsample, configured independently.
7. **Aggregation**. Fold → seed → task → task-equal mean. Confidence summaries are 10,000-resample task bootstraps with a fixed seed.
8. **Figures**. Regenerated from `source_data/` alone; each figure records the source file and its SHA-256 in `source_manifest.json`.
9. **Deviations**. Cross-attention and the transformer encoder were run under Core+K+X only. This is the only deviation from a complete operator × condition grid, and it is recorded in every table that would otherwise be affected.

### 23 Extended Discussion of Threats to Validity

#### Construct validity

Task-equal Macro-F1 measures performance on a fixed register of fifteen tasks (nine binary, six multiclass). It is not a measure of representation quality in the abstract, and two representations with identical task-equal scores may differ substantially on any single task. Figure 8 is provided precisely so that the register-level average is never read without the per-task spread.

#### Internal validity

The matching contract controls the gene universe, dimension, folds, seeds, probe, and aggregation. It does not control the optimizer trajectory, operator-specific capacity, or the interaction between capacity and the eight-epoch budget. An operator that is under-trained relative to its capacity would be penalized; the budget remains fixed because varying it per operator would break matching in a different place.

#### External validity

Five molecular and cellular views of human genes are used, with one external benchmark configuration. Nothing here supports a claim about other organisms, other view sets, tasks outside the register, or end-to-end supervised training.

#### Statistical conclusion validity

No significance test is reported. The fifteen tasks are a register, not a sample from a defined population of tasks, so a *p*-value would not have the interpretation a reader would expect. The reported intervals describe how much the task-equal mean would move if the register were resampled; they are not confidence intervals for a population parameter and they do not license a superiority claim.

### 24 Multiplicative-Interaction Results: Magnitude-Only Supplementary Evidence

A supplementary analysis evaluated whether explicit cross-view *multiplicative* interaction can raise the ceiling over additive (concatenation-based) fusion. Four experiments, conducted on 2026-08-06/07 and summarized in `results/multiplicative_sweep_recovered_20260911/`, span a synthetic case and an exhaustive real-task enumeration. The results bound the gains observed for interaction-based operators and provide magnitude-only supplementary evidence for the claims in the main text.

#### Evidence boundary

The analysis provides summary-level means, aggregate curves, and decision counts; fold-level values and interval endpoints are unavailable. Accordingly, the results are interpreted as magnitude evidence only and are not combined with the matched condition study. Where two metric conventions are possible, both readings are shown (`T1_synthetic_5seed_macro_f1.csv` versus `T1b_synthetic_alt_reading.csv`). Identifiers of the form `T1`–`T11` (e.g., `T2`, `T3`, `T4b`, `T5`, `T8`, `T11` below) are log-table filenames inside that run directory, not manuscript table numbers.

#### A. Synthetic existence proof

On an 800-gene, four-class synthetic task whose label is a function of cross-view latent products (score = *z*_*d*_*z*_*p*_ + 0.5 *z*_*r*_*z*_*p*_ + *ε*; random baseline Macro-F1 = 0.25), all four non-linear operators exceed both baselines in every one of five seeds (Table 26). The capacity-matched contrast isolates the structure itself: explicit products add +0.009 over an equal-capacity MLP on concatenated inputs (0.3340 vs. 0.3249), and soft-gated interaction adds +0.0946 over linear concatenation. Multiplicative interaction is therefore *detectable* when the ground truth is multiplicative.

#### B. Real-task scan

Across a 79-task inventory, the nine small/medium tasks evaluated with three folds and seed 42 show a positive multimodal gain in 61.1% of task × method combinations. The multiplicative operator exceeds linear fusion in only 9 of 21 bi/tri-modal effects (42.9%), indicating that the interaction term is unstable even when the multimodal trend is positive (`T2`, `T3`).

**Table 26:** Experiment A: five-seed mean Macro-F1 on the synthetic multiplicative-interaction task. Data file: `T1_synthetic_5seed_macro_f1.csv`.

| Operator | Family | Macro-F1 |
| --- | --- | --- |
| PCA add | baseline | 0.2471 |
| Linear concat | baseline | 0.2506 |
| Bilinear (products only) | non-linear | 0.3070 |
| MLP on concat (capacity control) | non-linear | 0.3249 |
| MLP + explicit interactions | non-linear | 0.3340 |
| Gated interaction | non-linear | **0.3452** |

#### C. Five-seed confirmation

Among the three task × combination cells with positive initial gains, no candidate simultaneously showed a positive gain, a paired-bootstrap interval excluding zero, and BH-FDR *q <* 0.05 against the capacity-matched MLP (`T4b`). The closest cell (bivalent versus non-methylated, DNA+protein) reached 0.7968 versus 0.7899; only 3/5 seeds were positive and the interval crossed zero.

#### D. Four-modality exhaustive enumeration

Enumerating all 15 non-empty subsets of DNA/RNA/protein/Text on two tasks (5 seeds × 3 folds) gives the ceiling directly (Table 27). Scores rise monotonically with the number of views—consistent with complementarity—but the multiplicative lead at four views (+0.0038 over the capacity-matched MLP) is six times *smaller* than its own seed standard deviation (0.0226). One task (long vs short range TF) is best solved by a *single* view with linear fusion (0.6226), a direct counterexample to “more views, more interaction, better”. A follow-up iteration (2026-08-07) with the evolved operator family found the low-rank higher-order expert winning 10 of 15 combinations yet still trailing the MLP by 0.000189 on the task-level mean (`T11`).

**Table 27:**
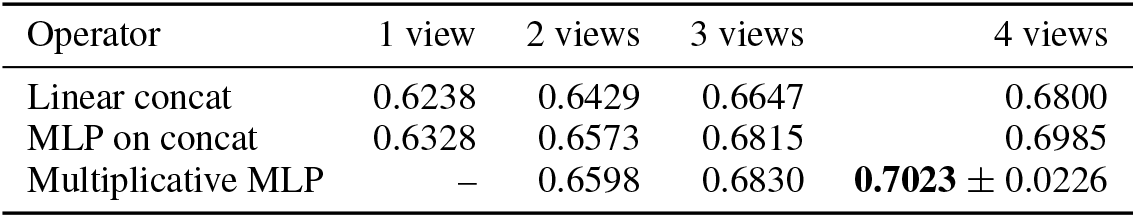
Experiment D: task-equal Macro-F1 by number of input views (mean over two tasks and 15 combinations), and the four-view full-input contrast. The multiplicative lead over the capacity-matched MLP (+0.0038) is below its seed SD (0.0226). Data file: `T5`, `T8`.

| Operator | 1 view | 2 views | 3 views | 4 views |
| --- | --- | --- | --- | --- |
| Linear concat | 0.6238 | 0.6429 | 0.6647 | 0.6800 |
| MLP on concat | 0.6328 | 0.6573 | 0.6815 | 0.6985 |
| Multiplicative MLP | – | 0.6598 | 0.6830 | **0.7023** ± 0.0226 |

#### What this bounds

The results show that multiplicative interaction is detectable on the synthetic task (A), appears inconsistently on real tasks (B), does not achieve multiplicity-corrected confirmation against a capacity-matched baseline (C), and remains below seed variability at full input (D). This is the regime in which the main-text *N*_*KX*_ decomposition falls: condition gains occur for some operators (Table 3), whereas the interaction component does not exceed its capacity-matched ceiling. The controlled condition contrasts and this supplementary analysis agree qualitatively; because fold-level records are unavailable, the latter contributes magnitude-only evidence and the confirmatory interpretation rests on the matched condition study. Potential semantic leakage in the text view, view occlusion, an independent held-out set, and strict parameter matching remain outside the present evidence.

### 25 Dynamic-Routing Evaluation Criteria

The branch-level router below defines an evaluation target suggested by the progressive construction. Its equations, diagnostics, and falsifiable predictions specify the comparisons required to assess conditional selection; the main-text results do not depend on this component.

#### Router equations

For each gene and condition, a label-free diagnostic vector

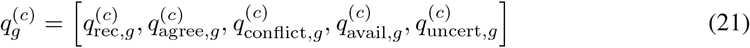

collects a self-supervised reconstruction residual, cross-view agreement and conflict statistics, the protocol availability statistic, and a representation-level uncertainty proxy, each computed without downstream labels. Writing 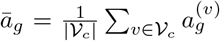 for the view centroid, 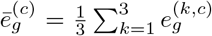 for the mean expert export, and *D*_*v*_ for the training-only decoders of Appendix 6, the components are defined as

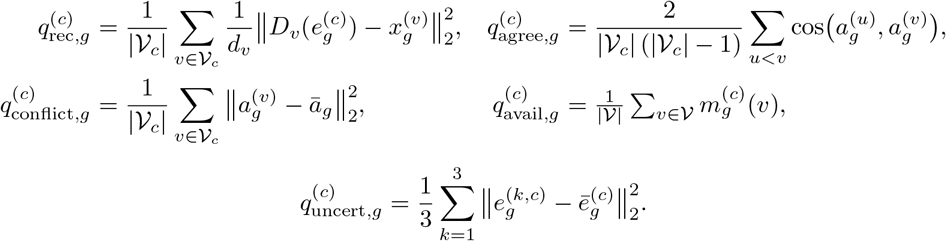

where cos(·, ·) denotes cosine similarity between adapted view vectors and 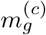 is the registered condition mask, so *q*_avail_ equals the constant |*V*_*c*_|*/*5 within a fixed condition. Under the resolved objective the reconstruction term is active, so all five statistics are defined. The router forms branch logits and weights

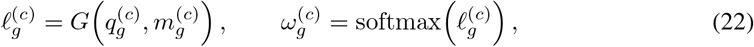

where 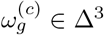, and mix the three expert exports,

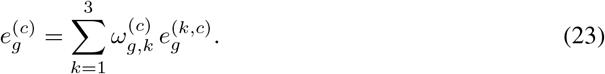

These weights describe a conditional choice among representation branches; they are not gene importance scores, assay-quality calibration, or causal effects. This branch-level router must not be confused with the per-view weighting of the Reliability expert (Eq. 4), which is part of the primary evaluation.

#### Four falsifiable predictions (prespecified)

(P1) If a task carries information no single view captures, the Interaction or LRHO expert exceeds the strongest single-view and concatenation references on that task. (P2) The LRHO gain varies with measurable view complementarity or residual dependence rather than remaining constant across tasks. (P3) Under controlled noise, conflict, or structured view removal, the router down-weights branches whose assumptions are degraded. (P4) Dynamic routing exceeds the best fixed expert only when expert risks are genuinely complementary and the label-free diagnostics predict that complementarity. The decisive comparisons are paired within task, seed, fold, condition, probe, and metric. An oracle advantage without a router advantage indicates complementarity with routing failure; the converse pattern supports conditional selection; no dynamic advantage requires downgrading the method claim to the fixed-expert comparison this paper reports.

The routing hypothesis is confirmed only if all of the following hold on a held-out target-task split: (i) the label-free router exceeds the best fixed expert by more than its task-bootstrap interval width; (ii) it exceeds a globally learned static mixture and random routing; (iii) the advantage persists under controlled noise and structured view removal; and (iv) an oracle that selects the best expert per task shows a materially larger advantage, establishing that complementarity exists. If (i)–(iii) hold but (iv) fails, the correct conclusion is conditional bias rather than expert complementarity. If (i) fails while the oracle advantage is large, the correct conclusion is complementarity with routing failure. This paper reports neither case; it supplies the fixed-expert baseline against which the test should be run.

### 26 Embedding-Geometry Analysis of Views and Fused Representations

This appendix examines the geometry of the five native views and of the fused representations directly, complementing the probe-based audit with probe-free evidence. It answers three questions the main text cannot: Are the five views actually complementary? Do the learned experts produce differently-structured embeddings from the baselines? And does the label-free objective—variance/covariance regularisation in the spirit of VICReg (Bardes et al., 2022)—leave a measurable geometric signature? All numbers below come from the recorded analysis run in `results/emb_analysis/` (scripts and CSV tables released with the artifact). **Scope caution:** the analysis run uses PCA-128 pre-reduction, hidden width 32, and twenty epochs for memory reasons, so its geometries are qualitative evidence about operator biases, not numbers from the benchmark configuration of Section 3; downstream scores are never re-derived here.

#### Native views show limited alignment under the reported geometry diagnostics

Pairwise linear CKA (Kornblith et al., 2019) between the five native views is uniformly below 0.04 (DNA–protein 0.027, DNA–RNA 0.015, protein–knowledge 0.003, knowledge–single-cell expression 0.001), and *k*-NN overlap at *k*=10 is at most 0.101 (Figure 4). These diagnostics indicate limited alignment of the measured geometries under this analysis configuration. They do not identify the biological source of downstream gains, exclude shared information at other scales, or establish that any fusion gain must arise from complementarity rather than shared-signal denoising. PCA spectra of the full-dimensional native views (Figure 3; `native_pca_statistics.csv`) show the views occupy different complexity regimes: DNA is nearly low-rank (effective rank 6.4; PC1 49%), whereas knowledge and single-cell expression are scattered across dimensions (effective ranks 42.6 and 35.5). These full-spectrum values describe the native input spaces; Table 28 instead reports effective ranks computed in the PCA-128 pre-reduced analysis space (native DNA 38.3, knowledge 118.9, single-cell expression 121.3), so the two sets of numbers are not directly comparable and are reported on their own bases.

**Figure 3:**
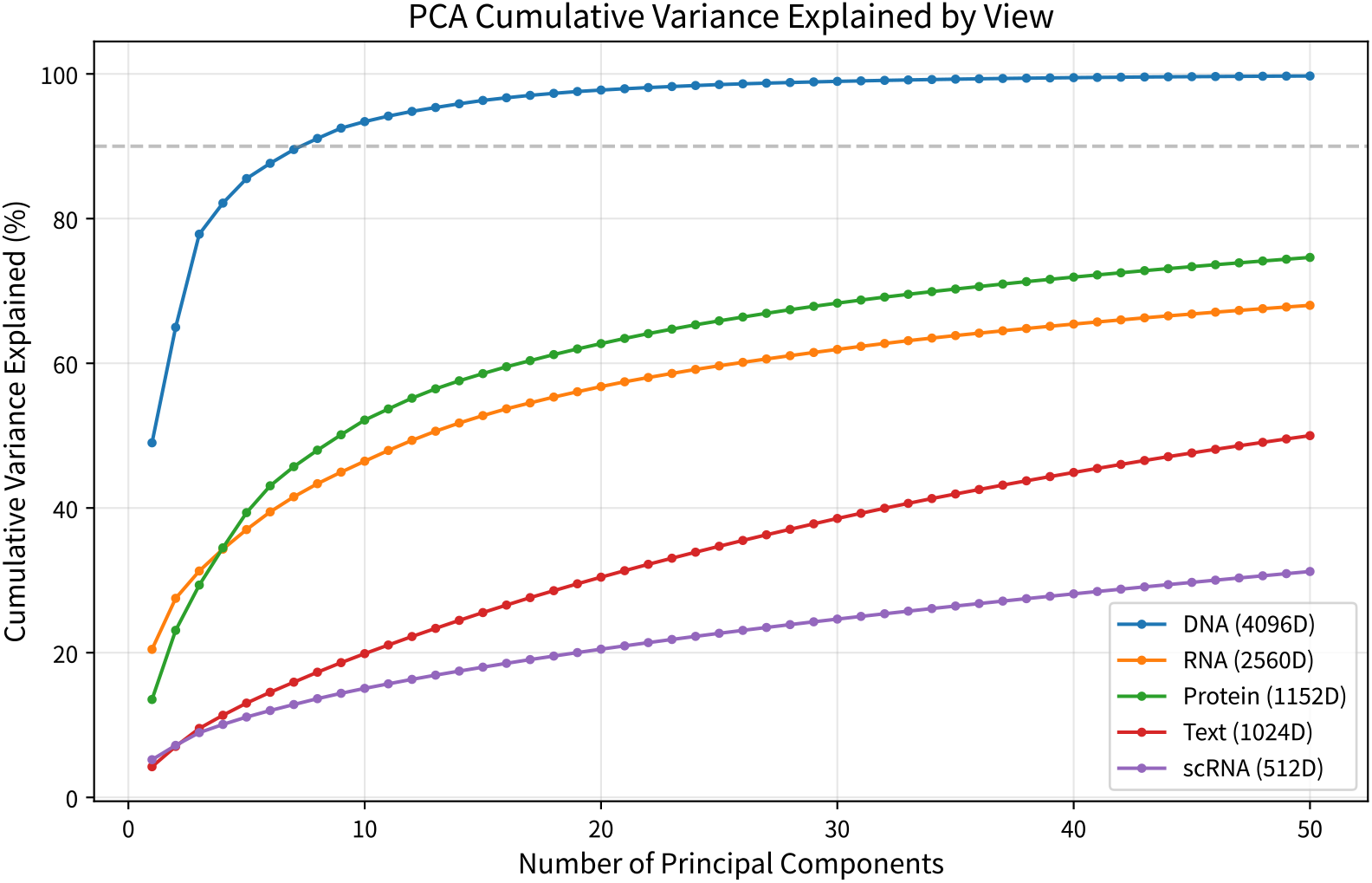
PCA variance explained for the five native views (first 50 components). DNA concentrates nearly all variance in a few components; knowledge and single-cell expression spread information across dimensions. Data file: `results/emb_analysis/tables/native_pca_statistics.csv`; method: scikit-learn PCA (Pedregosa et al., 2011).

**Figure 4:**
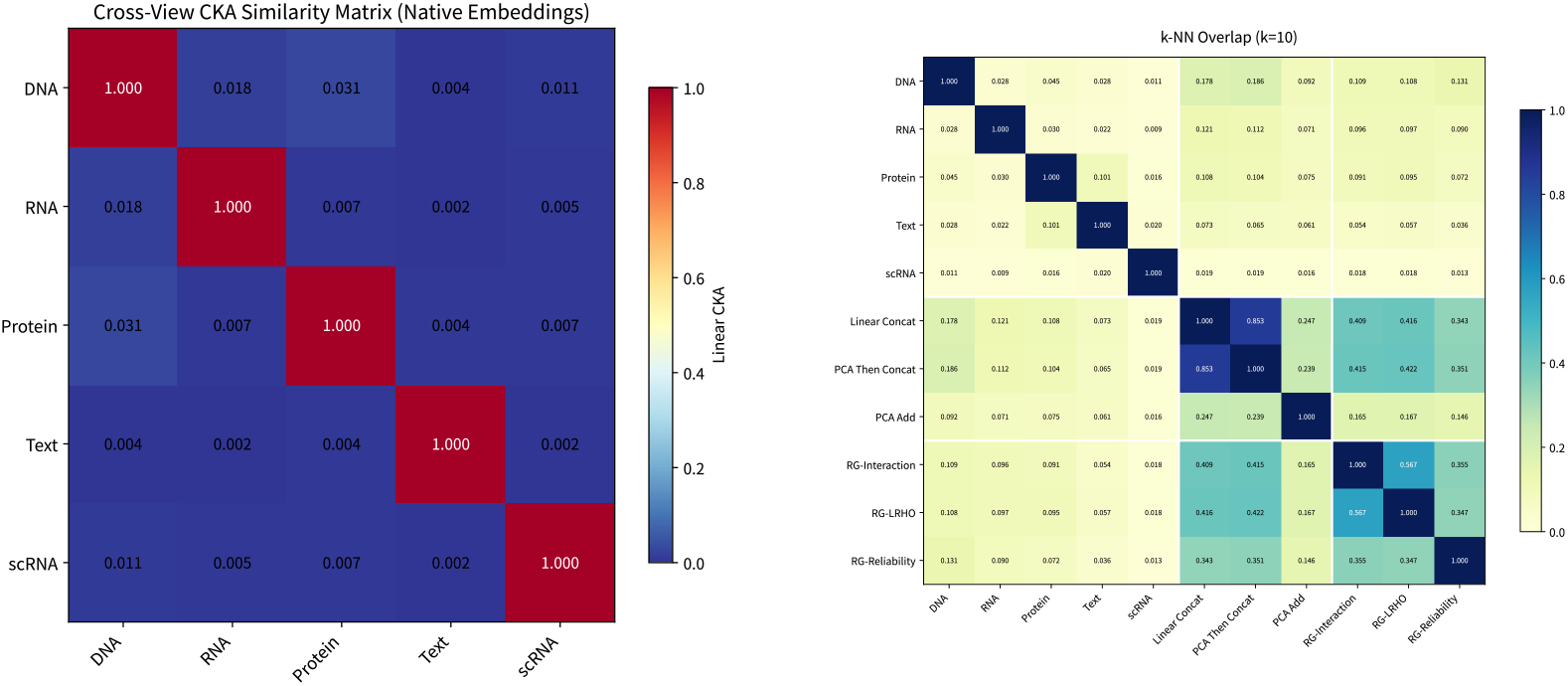
Native-view complementarity. Left: pairwise linear CKA between the five native views (all *<* 0.04). Right: *k*-NN (*k*=10) neighbour-set overlap between native views (at most 0.101). Sources: `cka_matrix_native.csv`, `knn_overlap_k10_native.csv`.

**Figure 5:**
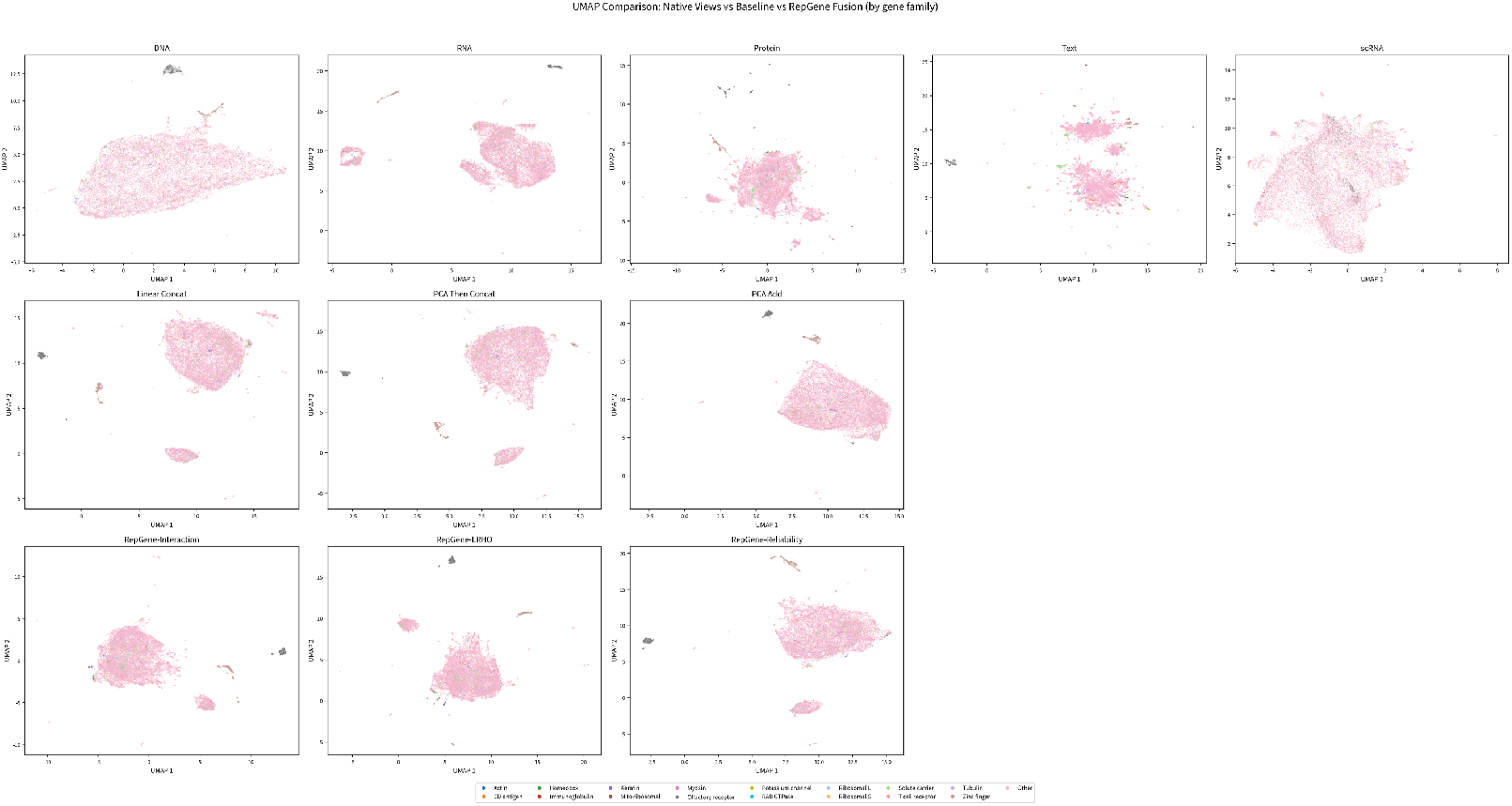
UMAP (McInnes et al., 2018) master panel of all eleven representations (five native views, three baselines, three RepGene experts), coloured by gene-family prefix labels. DNA-dominated structure is visible in the concatenation baselines; the learned experts produce visibly different, more uniform manifolds. A t-SNE (van der Maaten & Hinton, 2008) panel with identical layout is released with the artifact (`emb_tsne_master_panel.pdf`).

#### Learned experts produce different geometry from baselines

Three quantitative contrasts stand out (Table 28; Figures 6, 7). First, the learned experts are much more *compact*: effective rank 47.9– 74.3 versus 171.3–188.1 for the baselines, i.e., the label-free objective with variance/covariance regularisation concentrates representations onto fewer usable directions rather than preserving input dimensionality. Second, their norms are more *uniform* across genes than the two concatenation baselines (per-gene L2-norm spread 2.31–2.55 versus 5.42–5.45; Table 28), a geometric pattern consistent with the variance term, without isolating a single loss-term effect. PCA add is the most uniform representation overall (1.17), so norm uniformity separates the learned experts from the concatenation baselines but not from the additive baseline. Third, the three experts are more diverse *among themselves* than the baselines are (inter-method CKA 0.027–0.032 versus 0.091–0.103), consistent with the inductive-bias differences the audit is designed to expose; meanwhile every fused representation remains correlated with DNA (CKA up to 0.168), reflecting DNA’s outsized norm and low-rank concentration. The Reliability expert is the most compact (effective rank 47.9) and the least similar to Linear concat (*k*-NN overlap 0.343 at *k*=10), matching its input-conditioned view weighting. Silhouette scores against gene-family prefix labels are negative for *all* representations (Table 28), so no operator organises the space by these name-based families; we report this rather than claim cluster structure.

**Figure 6:**
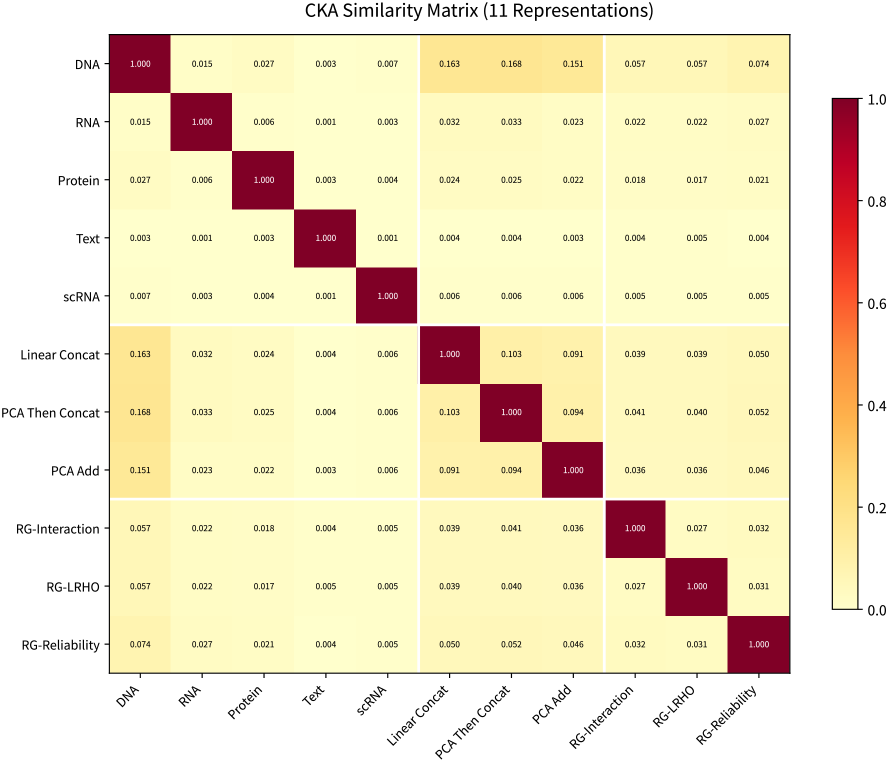
Linear CKA between all eleven representations. Baselines are mutually similar (0.09–0.10); learned experts are both less similar to baselines (0.036– 0.052) and more diverse among themselves (0.027– 0.032). Data file: `cka_matrix_all.csv`.

**Figure 7:**
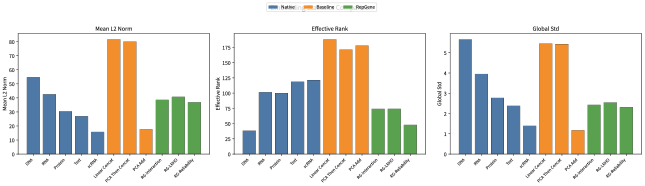
Geometry statistics for all representations: learned experts trade effective rank for norm uniformity under the variance-regularised objective. Data file: `embedding_stats_all.csv`.

**Table 28:** Embedding-geometry statistics (analysis-run configuration; qualitative evidence). Effective rank from the singular-value entropy; L2 spread is the per-gene norm standard deviation; silhouette against gene-family prefix labels (*n*=5,000 subsample (Rousseeuw, 1987)).

| Representation | Effective rank | L2 norm (mean) | L2 spread | Silhouette |
| --- | --- | --- | --- | --- |
| Native DNA | 38.3 | 54.6 | 5.66 | −0.136 |
| Native RNA | 101.4 | 42.5 | 3.95 | −0.055 |
| Native protein | 99.9 | 30.3 | 2.77 | −0.042 |
| Native knowledge | 118.9 | 26.9 | 2.38 | −0.097 |
| Native single-cell expression | 121.3 | 15.7 | 1.40 | −0.059 |
| Linear concat | 188.1 | 81.6 | 5.45 | −0.058 |
| PCA then concat | 171.3 | 80.0 | 5.42 | −0.048 |
| PCA add | 177.9 | 17.6 | 1.17 | −0.043 |
| RG-Interaction | 74.0 | 38.6 | 2.43 | −0.060 |
| RG-LRHO | 74.3 | 40.8 | 2.55 | −0.057 |
| RG-Reliability | 47.9 | 36.8 | 2.31 | −0.081 |

#### What this adds to the audit

The geometry line is independent of the probe: it shows (i) the premise of fusion (view complementarity) holds in this universe; (ii) the learned experts are not “the same embeddings with extra steps”—their induced geometry differs measurably and in the direction the objective predicts; and (iii) no representation, learned or not, organises genes by name-based families, which tempers biological-interpretation claims for *every* operator, including ours. These observations use distribution-level diagnostics (UMAP/t-SNE, CKA, neighbourhood overlap, rank and norm statistics) whose estimation does not touch task labels at any point.

### 27 Additional Evaluation Figures

The figures in this section provide additional evaluation views grouped by scientific function. Each caption identifies the displayed object, data source, and evidence boundary.

**Figure 8:**
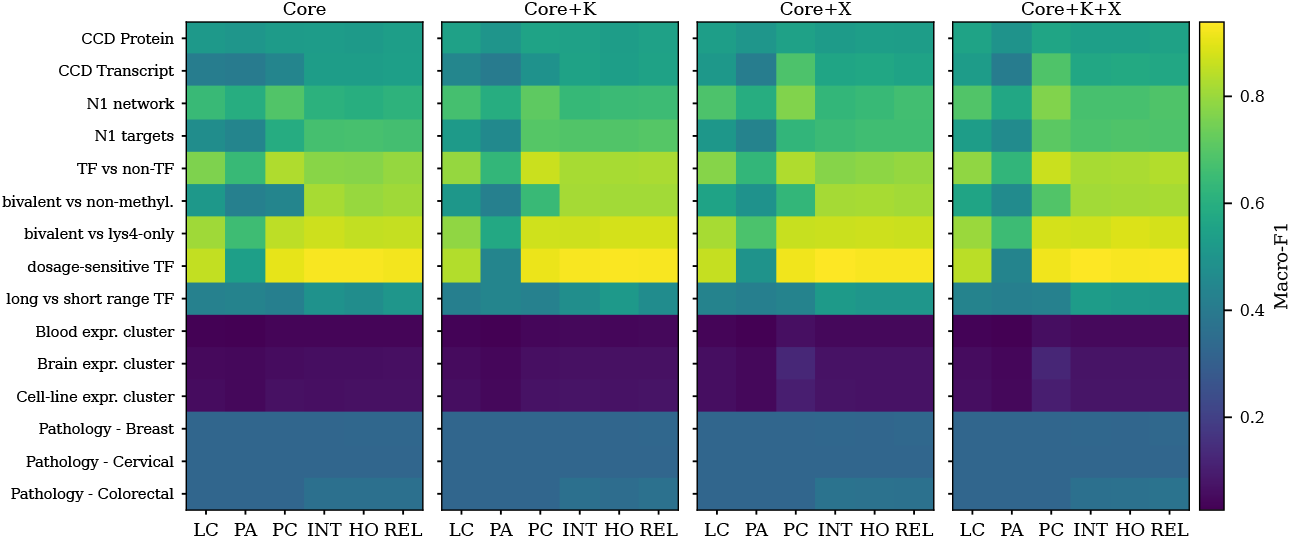
Task-level Macro-F1 for the fifteen-task register, six methods, and four protocol-defined conditions. The common colour scale keeps task heterogeneity visible without task filtering or centering; rows are grouped by task family (binary families above, multiclass families below). Data file: `full15_task_condition_method_long.csv`.

**Figure 9:**
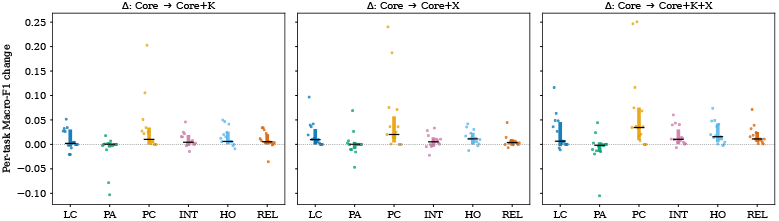
Matched condition differences relative to Core, one point per register task (fifteen per method). Thick bars show descriptive interquartile ranges. Data file: `full15_task_condition_method_long.csv`.

**Figure 10:**
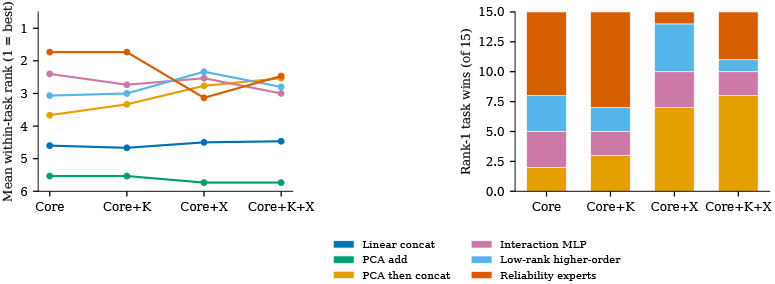
Descriptive within-condition mean ranks and rank-1 task wins on the fifteen-task register. No operator wins more than eight of fifteen tasks in any condition—the cleanest single no-dominance observation. Data file: `full15_task_condition_method_long.csv`.

**Figure 11:**
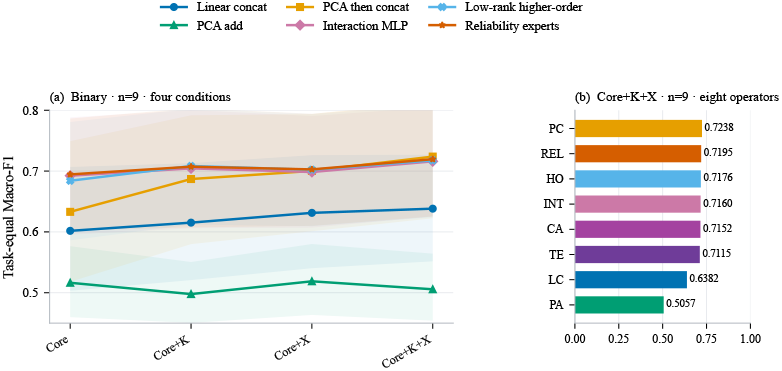
Binary-suite profile: task-equal Macro-F1 for the registered nine-task binary sub-register. Descriptive only; the primary fifteen-task register is reported in the main text and Appendix 10; Figure 2 is the binary continuity diagnostic. Data file: `binary_task_equal_summary_rf.csv`.

**Figure 12:**
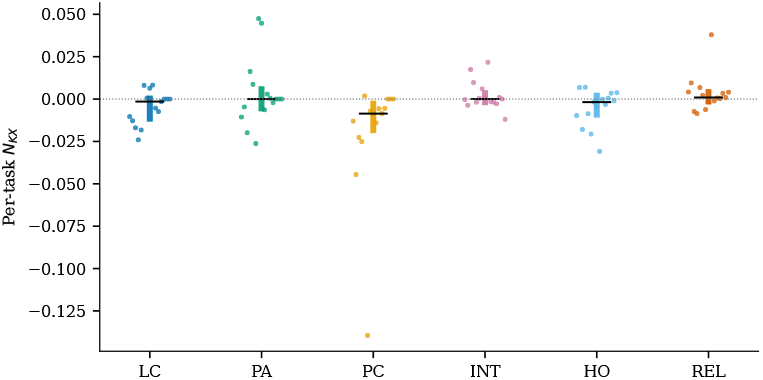
K/X non-additivity across fifteen paired register tasks per method. An arithmetic decomposition of the condition response, not biological synergy or causal interaction. Data file: `full15_task_condition_method_long.csv`.

### 28 Protocol-Extension Evidence: Probes, Training Budget, Pairing, and Calibration

The primary register of Section 4 is reported under one frozen configuration: a random-forest probe, an eight-epoch label-free budget, and the unmodified 0.5 decision threshold. This appendix reports four protocol extensions that vary one factor at a time while holding the rest of the contract fixed: the probe family (random forest, logistic regression, and a supervised MLP head), the training budget (8, 32, and 64 epochs), the gene-wise pairing of the interaction expert, and the calibration/threshold treatment of the stored probabilities. All extensions were executed on the same task files and the same embedding universe as the frozen register.

**Figure 13:**
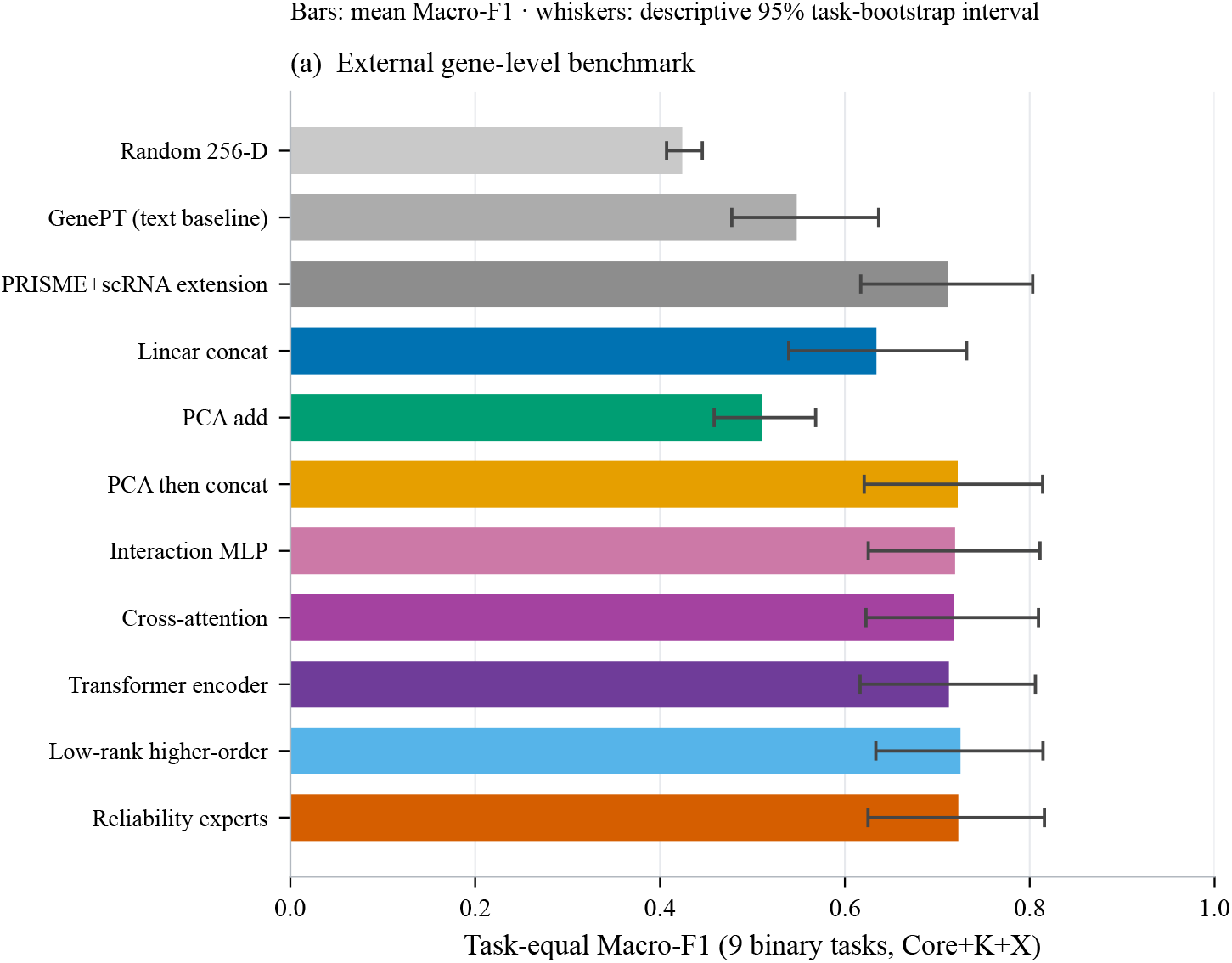
External Core+K+X gene-level track on nine binary tasks with the independently configured RF probe. The reported RepGene entries reproduce the numerical ordering range observed under the primary track; descriptive intervals overlap throughout. PRISME+scRNA denotes a five-view protocol extension rather than the original PRISME method. Data file: `benchmark_gene_external 2`.

**Figure 14:**
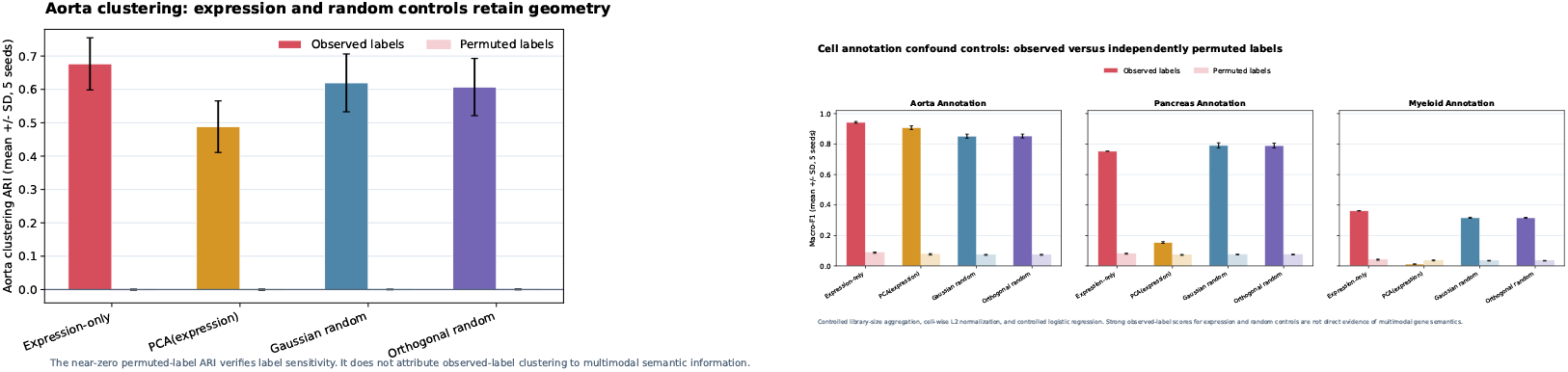
Cell-level boundary diagnostic, calibration-control view. Left: clustering (ARI) on the aorta annotation task; right: annotation Macro-F1 on three tasks (aorta; pancreas and myeloid as supplementary control-only panels whose dataset documentation is outside the reported aorta design point). Each panel plots the four label-permutation calibration controls (Expression-only, PCA(expression), Gaussian random, Orthogonal random) under observed versus permuted labels; the six fused representations of the matched contract are tabulated separately in Table 5 and are not drawn here. The panels show that observed-label scores remain high for all controls while permuted labels collapse, i.e. the aggregation channel itself carries the annotation signal—consistent with the dilution bound of Eq. 6—and Section 4.4 shows the fused operators do not separate from the dimension-matched random control under this aggregation. Sources: cell-level evaluation tables of Section 4.4.

**Table 29:** Summary of the four registered protocol extensions, each varying one factor of the frozen contract while the rest stays fixed. The following subsections give the full tables, the provenance of the re-execution, and the per-track caveats.

| Extension | Changed factor | Outcome |
| --- | --- | --- |
| Probe family | RF → logistic probe, identical exports (15 tasks × 4 conditions) | Structure survives: no operator leads in all four conditions and PCA then concat stays least decomposable ($N_{KX} = -0.0204$ RF, $-0.0116$ linear), but the leader changes with the probe and the experts’ edge over Linear concat does not survive (0.4929 vs. 0.4366 → 0.4964 vs. 0.5057) |
| Supervised head | RF → supervised MLP head (binary9 × Core+K+X) | Linear concat leads (0.7523); the learned experts follow (0.7257–0.7326) |
| Training budget | 8 → 32 → 64 epochs, four learned operators (binary9 × Core+K+X) | Every operator improves ($\approx +0.022/+0.031$), yet cross-attention never leads (residual gap $+0.0057/+0.0020/+0.0036$; at most two of nine per-task wins) |
| Pairing control | real pairing vs. capacity-matched shuffled twin | Real pairing wins in all four conditions: paired mean $+0.0259$, median $+0.0212$, 94% of 32 cells positive |
| Calibration | Brier/ECE and threshold sweeps from stored probabilities | PCA then concat ranks best (AUC 0.9387) and is calibrated worst (ECE 0.1270): it loses at the frozen 0.5 threshold (0.6866 vs. 0.7045) and wins once thresholds are tuned (0.7708 vs. 0.7372) |

#### Provenance and re-measurement fidelity

The extensions were produced by a re-execution of the released pipeline on a second machine (2026-09-18; `scikit-learn` 1.9.0, `numpy` 2.x, `torch` 2.13.0+cpu, against `numpy` 1.26.4 and `torch` 2.1.0+cu121 at freeze time), with the released configuration files as the only input. Two checks bound the comparison: (i) the per-fold train/test index hashes and the sample counts of the re-executed cells are *identical* to the frozen records, so the task files, label sets, and splits are unchanged; (ii) on the binary sub-register under Core+K+X, the re-executed epoch-8 task-equal means agree with the frozen values to four decimals (Reliability experts 0.6995 vs. 0.6993; Interaction MLP 0.6965 vs. 0.6963; PCA then concat 0.7037 vs. 0.7042), i.e. library-level randomness does not perturb the aggregate. Comparisons inside a track below are therefore matched, and cross-track comparisons with the frozen register inherit this verified agreement.

#### 28.1 Probe sensitivity: the pattern survives, the ordering does not

Table 30 reports the fifteen-task register under the frozen random-forest probe and under a logistic-regression probe, both evaluated on the *same* exported representations, so probe comparisons are fully matched within a cell. Three observations follow.

First, the qualitative claim of the main text is probe-robust: no operator leads in all four conditions under either probe, and the per-condition leaders differ between the two probes (boldface marks the per-condition leader). Second, one specific claim is also probe-robust: PCA then concat remains the least decomposable operator, with a negative non-additivity term under both probes (*N*_*KX*_ = −0.0204 RF, −0.0116 linear). Third, one claim is *not* probe-robust: the numerical advantage of the learned experts over Linear concat under the RF probe (0.4929 vs. 0.4366 at Core) disappears under the linear probe (0.4964 vs. 0.5057), and the supervised MLP-head reference of Table 31 likewise places Linear concat highest (0.7523) with the experts below it (0.7257–0.7326). A trained head therefore does not by itself rescue the learned operators. This is the concrete form of the protocol-dependence claim: the register’s *structure* is stable across probes, while which operator “wins” is not.

**Table 30:** Fifteen-task task-equal Macro-F1 under two probes evaluated on identical exported representations (three seeds, three folds, four conditions). Boldface marks the per-condition leader. Bootstrap intervals of width ≈ ±0.13 overlap throughout, as in the main register.

| Method | Core | Core+K | Core+X | Core+K+X |
| --- | --- | --- | --- | --- |
| *RF probe (frozen primary)* |  |  |  |  |
| Linear concat | 0.4366 | 0.4448 | 0.4544 | 0.4588 |
| PCA add | 0.3838 | 0.3711 | 0.3853 | 0.3765 |
| PCA then concat | 0.4565 | 0.4898 | **0.5058** | **0.5188** |
| Interaction MLP | 0.4929 | 0.5053 | 0.5021 | 0.5118 |
| Low-rank higher-order | 0.4920 | **0.5056** | 0.5020 | 0.5109 |
| Reliability experts | **0.4951** | 0.5034 | 0.5019 | 0.5135 |
| *linear (logistic) probe* |  |  |  |  |
| Linear concat | **0.5057** | 0.5190 | 0.5326 | 0.5355 |
| PCA add | 0.4668 | 0.4598 | 0.4602 | 0.4605 |
| PCA then concat | 0.5019 | **0.5201** | **0.5357** | **0.5423** |
| Interaction MLP | 0.4964 | 0.5056 | 0.5060 | 0.5179 |
| Low-rank higher-order | 0.4902 | 0.5076 | 0.5057 | 0.5185 |
| Reliability experts | 0.4978 | 0.5060 | 0.5094 | 0.5199 |

**Table 31:** Supervised MLP-head reference (B2) on the nine binary tasks under Core+K+X, alongside the frozen random-forest probe on the same exported representations. The trained head does not restore the learned experts’ advantage over the concatenation baselines.

| Method | Supervised MLP head | RF probe |
| --- | --- | --- |
| Linear concat | 0.7523 | 0.6385 |
| PCA then concat | 0.7483 | 0.7236 |
| Low-rank higher-order | 0.7326 | 0.7206 |
| Interaction MLP | 0.7261 | 0.7184 |
| Reliability experts | 0.7257 | 0.7169 |
| PCA add | 0.6707 | 0.5059 |

#### 28.2 Training budget: more epochs raise every operator and do not rescue attention

Table 32 repeats the nine binary tasks under Core+K+X at 32 and 64 epochs, alongside the same run’s 8-epoch values and the frozen epoch-8 reference. Raising the budget helps every family: the task-equal mean rises by roughly 0.022 at 32 epochs and 0.031 at 64 epochs relative to the matched 8-epoch row, so the register’s absolute levels are budget-limited. The ordering nevertheless does not change: the best learned operator at every budget is a reliability- or low-rank-style expert, and cross-attention stays below it (0.6936 → 0.7198 → 0.7318 against 0.6993 → 0.7217 → 0.7354), with a residual gap of +0.0057/+0.0020/+0.0036 and 0–2 per-task wins out of nine. Because that residual gap is of the same order as the seed noise (± 0.005), the accurate reading of the attention result is that it *never attains an advantage at any tested budget*, not that it is reliably worse. The transformer encoder was never frozen into the released snapshot and could not be re-executed, so the budget statement is restricted to cross-attention.

#### 28.3 View-pair shuffle control: the interaction gain is genuine pairing

The interaction expert adds pairwise products and absolute differences between views. A capacity-matched twin (`mlp_interaction_shuffled`) keeps the identical parameter count (20,168 in a unit test), the identical feature dimension, and the identical concatenation block, but forms each pair term against a row-permuted copy of the second view, so gene-wise cross-view pairing is destroyed by construction while every marginal distribution is preserved. Table 33 shows that real pairing is better in all four conditions, by +0.0173 to +0.0363 task-equal Macro-F1 (paired *n* = 32 task×condition cells: mean +0.0259, median +0.0212, 94% positive). The interaction advantage is thus attributable to genuine cross-view pairing rather than to generic nonlinear capacity or added parameters.

**Table 32:** Nine binary tasks under Core+K+X, task-equal Macro-F1 against the label-free training budget. Columns 2–3 are the matched re-execution; the last column is the frozen epoch-8 reference for drift comparison. PCA then concat and Linear concat were not re-run at higher budgets (the budget question concerns the learned operators) and are shown for the epoch-8 reference only.

| Method | 8 epochs | 32 epochs | 64 epochs | 8 epochs (frozen ref.) |
| --- | --- | --- | --- | --- |
| Reliability experts | 0.6995 | 0.7217 | 0.7354 | 0.6993 |
| Interaction MLP | 0.6965 | 0.7208 | 0.7308 | 0.6963 |
| Low-rank higher-order | 0.6957 | 0.7182 | 0.7279 | 0.6958 |
| Cross-attention | – | 0.7198 | 0.7318 | 0.6936 |
| PCA then concat | – | – | – | – |
| Linear concat | – | – | – | – |

**Table 33:** View-pair shuffle control on the nine binary tasks (three seeds, four conditions, random-forest probe). The shuffled twin is capacity-matched; only the gene-wise cross-view pairing is destroyed.

| Variant | Core | Core+K | Core+X | Core+K+X |
| --- | --- | --- | --- | --- |
| Real pairing | 0.6702 | 0.6888 | 0.6768 | 0.6974 |
| Pair-shuffled twin | 0.6529 | 0.6619 | 0.6538 | 0.6611 |
| Difference | +0.0173 | +0.0269 | +0.0230 | +0.0363 |

#### 28.4 Calibration and threshold sensitivity explain the metric inversions

Section 4.3 reports that Macro-F1 and ROC-AUC disagree about the operator ordering. Table 34 supplies the deciding analysis on the binary sub-register, computed from the stored per-gene probabilities. Under the RF probe, PCA then concat has the highest rank discrimination (AUC 0.9387) and the *worst* calibration (ECE 0.1270, Brier 0.3291), while the learned experts are the best calibrated (ECE 0.0730–0.0761) and lead at the frozen 0.5 threshold (0.7045 vs. 0.6866). Re-tuning the threshold per operator reverses the Macro-F1 ordering: PCA then concat reaches 0.7708 against 0.7347–0.7372 for the experts. The inversion is therefore a threshold/calibration effect rather than a change in ranking quality, and the fixed 0.5 threshold is not a neutral comparison point. Under the linear probe the ordering changes once more (Linear concat 0.7318, PCA then concat 0.7291, experts 0.7084–0.7146), so the operator ordering is jointly sensitive to probe *and* threshold, while the calibration advantage of the learned experts persists in every probe tested. All values are task-equal means over the nine binary tasks.

**Table 34:**
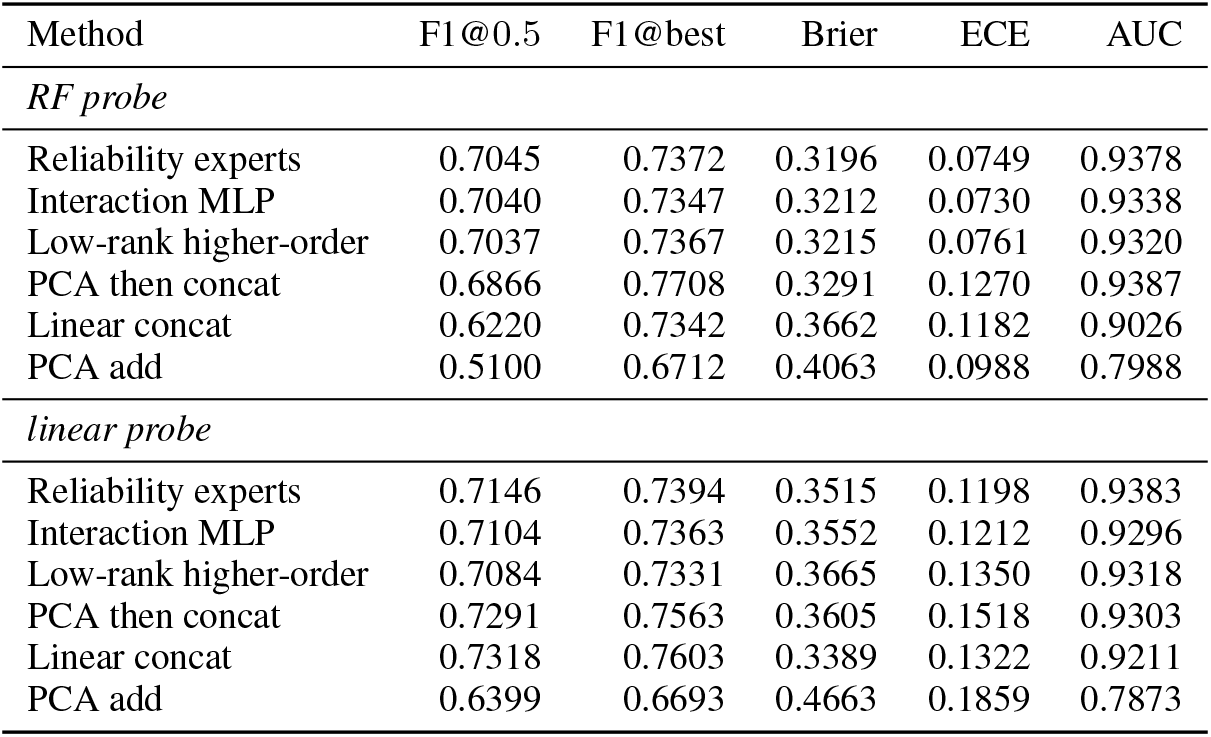
Calibration and threshold sensitivity on the nine binary tasks, task-equal means. Macro-F1 is reported at the frozen 0.5 threshold and at the best swept threshold of the same probabilities; Brier and ECE quantify calibration; ROC-AUC quantifies rank discrimination.

| Method | F1@0.5 | F1@best | Brier | ECE | AUC |
| --- | --- | --- | --- | --- | --- |
| *RF probe* |  |  |  |  |  |
| Reliability experts | 0.7045 | 0.7372 | 0.3196 | 0.0749 | 0.9378 |
| Interaction MLP | 0.7040 | 0.7347 | 0.3212 | 0.0730 | 0.9338 |
| Low-rank higher-order | 0.7037 | 0.7367 | 0.3215 | 0.0761 | 0.9320 |
| PCA then concat | 0.6866 | 0.7708 | 0.3291 | 0.1270 | 0.9387 |
| Linear concat | 0.6220 | 0.7342 | 0.3662 | 0.1182 | 0.9026 |
| PCA add | 0.5100 | 0.6712 | 0.4063 | 0.0988 | 0.7988 |
| *linear probe* |  |  |  |  |  |
| Reliability experts | 0.7146 | 0.7394 | 0.3515 | 0.1198 | 0.9383 |
| Interaction MLP | 0.7104 | 0.7363 | 0.3552 | 0.1212 | 0.9296 |
| Low-rank higher-order | 0.7084 | 0.7331 | 0.3665 | 0.1350 | 0.9318 |
| PCA then concat | 0.7291 | 0.7563 | 0.3605 | 0.1518 | 0.9303 |
| Linear concat | 0.7318 | 0.7603 | 0.3389 | 0.1322 | 0.9211 |
| PCA add | 0.6399 | 0.6693 | 0.4663 | 0.1859 | 0.7873 |

##### Scope of the extensions

The probe-sensitivity sweep covers the full fifteen-task register and four conditions with three seeds; the budget scan covers the nine binary tasks under Core+K+X; the shuffle control covers the nine binary tasks under all four conditions; the supervised reference covers the nine binary tasks under Core+K+X. The primary register of Section 4 remains the frozen run; the re-executed tracks are used for matched within-track contrasts and for the documented agreement check above, and no extension upgrades a claim from descriptive to inferential. The homogeneous input-condition states, the label-free training objective, and the shared annotation families of the register are unchanged by these extensions.

