## Supplementary_Material for "RepGene: Auditing Condition-Specific Fusion Biases for Multi-View Gene Representations": analysis_budget_shuffle.py

```
"""Training-budget scan (Q3) and view-pair shuffle control (Q5).

Budget scan: binary9 x Core+K+X, learned operators at epochs {8 (frozen), 32, 64}.
  - epoch 8 baselines come from the frozen runs: the four main operators from the
    15-task register and cross-attention from the binary new-methods run.
  - epochs 32/64 come from the Pod re-runs produced here.
Reports task-equal Macro-F1, the attention gap, and per-task win counts.

Shuffle control: Interaction MLP vs its pair-shuffled twin across the four
conditions on binary9, task-equal means with paired per-task deltas.
"""
from __future__ import annotations

import glob
import sys
from pathlib import Path

import numpy as np
import pandas as pd

ROOT = Path("${REPO_ROOT}/results")
BINARY9 = ["CCD Protein", "CCD Transcript", "N1 network", "N1 targets", "TF vs non-TF",
           "bivalent vs non-methylated", "bivalent_vs_lys4_only",
           "dosage sensitive vs insensitive TF", "long vs short range TF"]
RENAME = {"bivalent_vs_lys4_only": "bivalent vs lys4-only"}
METHOD_MAP = {
    "fusion_mlp_interaction_unsup": "Interaction MLP",
    "fusion_lowrank_higher_order_unsup": "Low-rank higher-order",
    "fusion_reliability_experts_unsup": "Reliability experts",
    "fusion_cross_attention_unsup": "Cross-attention",
    "fusion_mlp_interaction_unsup_shuffled": "Interaction MLP (pair-shuffled)",
}
COND_MAP = {"DNA_RNA_protein": "Core", "DNA_RNA_protein_text": "Core+K",
            "DNA_RNA_protein_scRNA": "Core+X", "DNA_RNA_protein_text_scRNA": "Core+K+X"}


def read_metrics(run_subdir: str, methods: set[str] | None = None) -> pd.DataFrame:
    rows = []
    for f in glob.glob(f"{ROOT}/{run_subdir}/task=*/modalities=*/methods=*/seed=*/metrics_long.csv"):
        p = Path(f).parts
        task = [x for x in p if x.startswith("task=")][0].split("=", 1)[1]
        cond = [x for x in p if x.startswith("modalities=")][0].split("=", 1)[1]
        seed = [x for x in p if x.startswith("seed=")][0].split("=", 1)[1]
        df = pd.read_csv(f)
        df["task"] = df["task"].replace({"bivalent_vs_lys4_only": "bivalent vs lys4-only"})
        df["condition"] = COND_MAP.get(cond, cond)
        df["seed"] = int(seed)
        rows.append(df)
    d = pd.concat(rows, ignore_index=True)
    d = d[d["task"].isin(BINARY9)]
    if methods:
        d = d[d["representation_method"].isin(methods)]
    return d


def cell_means(d: pd.DataFrame, probe: str = "rf") -> pd.DataFrame:
    sub = d[d["probe"] == probe]
    return (sub.groupby(["task", "condition", "representation_method"], as_index=False)
            .agg(macro_f1=("macro_f1", "mean"), roc_auc=("roc_auc", "mean"),
                 balanced_accuracy=("balanced_accuracy", "mean")))


def main():
    out_dir = ROOT / "analysis"
    out_dir.mkdir(exist_ok=True)

    # ---------------- budget scan ----------------
    e8_main = read_metrics("core_k_x_fulltask_20260908/core_k_x_fulltask_20260908")
    e8_attn = read_metrics("core_k_x_fulltask_20260908/core_k_x_new_methods_binary_20260910")
    e32 = read_metrics("budget_scan_epochs32_20260918/budget_scan_epochs32_20260918")
    e64 = read_metrics("budget_scan_epochs64_20260918/budget_scan_epochs64_20260918")

    frames = []
    e8_new = read_metrics("probe_track_20260918/probe_track_20260918")   # same environment as 32/64
    for ep, d, tag in [(8, e8_new, "matched"), (32, e32, "matched"), (64, e64, "matched"),
                       (8, pd.concat([e8_main, e8_attn], ignore_index=True), "frozen-ref")]:
        cm = cell_means(d)
        cm = cm[cm["condition"] == "Core+K+X"]
        cm["epochs"] = ep
        cm["env"] = tag
        frames.append(cm)
    bud = pd.concat(frames, ignore_index=True)
    bud["method"] = bud["representation_method"].map(METHOD_MAP).fillna(bud["representation_method"])
    bud.to_csv(out_dir / "budget_scan_per_task.csv", index=False)

    print("=== Budget scan (binary9 x Core+K+X, RF probe, task-equal Macro-F1) ===")
    matched = bud[bud.env == "matched"]
    piv = matched.pivot_table(index="method", columns="epochs", values="macro_f1", aggfunc="mean")
    print("[same Pod environment across budgets]")
    print(piv.round(4))
    print("\nper-epoch gap = best learned operator - cross-attention:")
    for ep in [8, 32, 64]:
        s = piv[ep].drop("Cross-attention")
        print(f"  epochs={ep:>3}: best={s.idxmax()} {s.max():.4f} | cross-attention {piv.loc['Cross-attention', ep]:.4f} "
              f"| gap {s.max() - piv.loc['Cross-attention', ep]:+.4f}")
    print("\nper-task attention wins (of 9):",
          {ep: int((matched[matched.epochs == ep].pivot_table(index="task", columns="method",
                                                             values="macro_f1").idxmax(axis=1) == "Cross-attention").sum())
           for ep in [8, 32, 64]})
    ref = bud[bud.env == "frozen-ref"].pivot_table(index="method", columns="epochs", values="macro_f1", aggfunc="mean")
    print("\n[frozen-environment epoch-8 reference, for drift comparison]")
    print(ref.round(4))

    # ---------------- shuffle control ----------------
    sh = read_metrics("shuffle_control_20260918/shuffle_control_20260918")
    shm = cell_means(sh)
    shm["method"] = shm["representation_method"].map(METHOD_MAP)
    shm.to_csv(out_dir / "shuffle_control_per_task.csv", index=False)
    print("\n=== View-pair shuffle control (binary9, RF probe, task-equal Macro-F1) ===")
    tab = shm.pivot_table(index="method", columns="condition", values="macro_f1", aggfunc="mean")
    print(tab.round(4))
    wide = shm.pivot_table(index=["task", "condition"], columns="method", values="macro_f1")
    wide["delta"] = wide["Interaction MLP"] - wide["Interaction MLP (pair-shuffled)"]
    print("\nreal - shuffled delta: mean {:+.4f}, median {:+.4f}, share>0 {:.2f}, n={}".format(
        wide["delta"].mean(), wide["delta"].median(), (wide["delta"] > 0).mean(), len(wide)))
    per_cond = wide.reset_index().groupby("condition")["delta"].agg(["mean", "median"])
    print(per_cond.round(4))
    wide.reset_index().to_csv(out_dir / "shuffle_control_paired.csv", index=False)
    print("\nwrote:", out_dir / "budget_scan_per_task.csv", "and shuffle_control_per_task.csv / _paired.csv")


if __name__ == "__main__":
    main()
```
