## Supplementary_Material for "RepGene: Auditing Condition-Specific Fusion Biases for Multi-View Gene Representations": analysis_calibration.py

```
"""Calibration analysis (Q6) from saved per-gene probabilities.

For every (task, condition, method, probe, fold) reads predictions/method=*/probe=*/fold=*.jsonl
and computes:
  - Brier score (multi-class sum-of-squares convention, mean over samples)
  - ECE (expected calibration error on the top-label confidence, 15 equal-width bins)
  - Macro-F1 at the default argmax decision
  - for binary tasks: Macro-F1 at the best swept threshold and the sweep curve summary
Outputs per-cell tables plus task-equal aggregates per probe/method/condition.
"""
from __future__ import annotations

import glob
import json
import sys
from pathlib import Path

import numpy as np
import pandas as pd
from sklearn.metrics import f1_score

COND_MAP = {
    "DNA_RNA_protein": "Core",
    "DNA_RNA_protein_text": "Core+K",
    "DNA_RNA_protein_scRNA": "Core+X",
    "DNA_RNA_protein_text_scRNA": "Core+K+X",
}
METHOD_MAP = {
    "linear_concat": "Linear concat",
    "pca_add": "PCA add",
    "pca_then_concat": "PCA then concat",
    "fusion_mlp_interaction_unsup": "Interaction MLP",
    "fusion_lowrank_higher_order_unsup": "Low-rank higher-order",
    "fusion_reliability_experts_unsup": "Reliability experts",
    "fusion_cross_attention_unsup": "Cross-attention",
    "fusion_mlp_interaction_unsup_shuffled": "Interaction MLP (pair-shuffled)",
}
BINARY_TASKS = {"CCD Protein", "CCD Transcript", "N1 network", "N1 targets",
                "TF vs non-TF", "bivalent vs non-methylated", "bivalent_vs_lys4_only",
                "dosage sensitive vs insensitive TF", "long vs short range TF"}


def brier_ece(y_true: np.ndarray, probs: np.ndarray, n_bins: int = 15):
    n, k = probs.shape
    onehot = np.zeros_like(probs)
    onehot[np.arange(n), y_true] = 1.0
    brier = float(np.mean(np.sum((probs - onehot) ** 2, axis=1)))
    conf = probs.max(axis=1)
    pred = probs.argmax(axis=1)
    correct = (pred == y_true).astype(float)
    edges = np.linspace(0.0, 1.0, n_bins + 1)
    ece = 0.0
    for i in range(n_bins):
        m = (conf > edges[i]) & (conf <= edges[i + 1])
        if m.sum() > 0:
            ece += (m.mean()) * abs(correct[m].mean() - conf[m].mean())
    return brier, float(ece)


def best_threshold_f1(y_true: np.ndarray, probs: np.ndarray):
    """Binary only: Macro-F1 at the best swept threshold on P(class 1).

    Vectorised sweep (no per-threshold sklearn call) so the full 180-cell
    probe track can be processed in seconds.
    """
    p1 = probs[:, 1]
    thresholds = np.arange(0.05, 0.96, 0.02)
    pos = (y_true == 1)
    n_pos, n_neg = pos.sum(), (~pos).sum()
    best = (-1.0, 0.5)
    for t in thresholds:
        pred_pos = p1 >= t
        tp = np.logical_and(pred_pos, pos).sum()
        fp = np.logical_and(pred_pos, ~pos).sum()
        fn = n_pos - tp
        tn = n_neg - fp
        f1_pos = 2 * tp / (2 * tp + fp + fn) if (2 * tp + fp + fn) else 0.0
        f1_neg = 2 * tn / (2 * tn + fn + fp) if (2 * tn + fn + fp) else 0.0
        macro = 0.5 * (f1_pos + f1_neg)
        if macro > best[0]:
            best = (float(macro), float(t))
    return best


def main():
    run_dir = Path(sys.argv[1] if len(sys.argv) > 1 else
                   "${REPO_ROOT}/results/probe_track_20260918/probe_track_20260918")
    out_dir = Path(sys.argv[2]) if len(sys.argv) > 2 else run_dir.parent / "analysis"
    out_dir.mkdir(parents=True, exist_ok=True)

    rows = []
    files = glob.glob(f"{run_dir}/task=*/modalities=*/methods=*/seed=*/predictions/method=*/probe=*/fold=*.jsonl")
    print("prediction files:", len(files))
    for f in files:
        parts = Path(f).parts
        task_dir = [p for p in parts if p.startswith("task=")][0].split("=", 1)[1]
        cond_dir = [p for p in parts if p.startswith("modalities=")][0].split("=", 1)[1]
        seed_dir = [p for p in parts if p.startswith("seed=")][0].split("=", 1)[1]
        method_dir = [p for p in parts if p.startswith("method=")][0].split("=", 1)[1]
        probe_dir = [p for p in parts if p.startswith("probe=")][0].split("=", 1)[1]
        fold = Path(f).stem.split("=", 1)[1]
        recs = [json.loads(l) for l in open(f)]
        if not recs:
            continue
        y_true = np.array([r["y_true"] for r in recs])
        probs = np.array([r["probabilities"] for r in recs], dtype=float)
        if probs.ndim == 1:  # defensive: binary stored as scalar probability
            probs = np.stack([1 - probs, probs], axis=1)
        brier, ece = brier_ece(y_true, probs)
        pred = probs.argmax(axis=1)
        row = {
            "task": task_dir, "condition": COND_MAP.get(cond_dir, cond_dir), "seed": int(seed_dir),
            "fold": int(fold), "method": METHOD_MAP.get(method_dir, method_dir), "probe": probe_dir,
            "brier": brier, "ece": ece,
            "macro_f1_default": float(f1_score(y_true, pred, average="macro", zero_division=0)),
            "n": len(recs),
        }
        # threshold sweeps are only defined for the binary sub-register, where the
        # stored probability vector has exactly two entries
        if probs.shape[1] == 2:
            f1_best, thr = best_threshold_f1(y_true, probs)
            row["macro_f1_best_threshold"] = f1_best
            row["best_threshold"] = thr
        rows.append(row)
    if not rows:
        print("no predictions found")
        return
    df = pd.DataFrame(rows)
    df.to_csv(out_dir / "calibration_per_fold.csv", index=False)

    agg = (df.groupby(["task", "condition", "method", "probe"], as_index=False)
           .agg(brier=("brier", "mean"), ece=("ece", "mean"),
                macro_f1_default=("macro_f1_default", "mean"),
                macro_f1_best_threshold=("macro_f1_best_threshold", "mean")))
    agg.to_csv(out_dir / "calibration_per_task.csv", index=False)

    for probe in sorted(agg["probe"].unique()):
        sub = agg[agg["probe"] == probe]
        te = sub.groupby(["condition", "method"], as_index=False)[["brier", "ece", "macro_f1_default"]].mean()
        print(f"\n=== calibration | probe={probe} (task-equal means) ===")
        print(te.pivot_table(index="method", columns="condition",
                             values=["brier", "ece"]).round(4))
    print("\nwrote:", out_dir / "calibration_per_task.csv", "and calibration_per_fold.csv")


if __name__ == "__main__":
    main()
```
