## Supplementary_Material for "RepGene: Auditing Condition-Specific Fusion Biases for Multi-View Gene Representations": analysis_probe_track.py

```
"""Aggregate the probe-sensitivity track (B1) and the supervised MLP reference (B2).

Reads results/<run>/<run>/task=*/modalities=*/methods=*/seed=*/metrics_long.csv
(written by scripts/run_benchmark.py) and reports, per probe:
  - task-equal Macro-F1 with descriptive task-cluster bootstrap intervals
  - matched condition deltas (Delta_K, Delta_X, Delta_KX) and N_KX
  - the per-condition top operator (does the condition-dependent pattern survive?)
"""
from __future__ import annotations

import glob
import sys
from pathlib import Path

import numpy as np
import pandas as pd

COND_MAP = {
    "DNA_RNA_protein": "Core",
    "DNA_RNA_protein_text": "Core+K",
    "DNA_RNA_protein_scRNA": "Core+X",
    "DNA_RNA_protein_text_scRNA": "Core+K+X",
}
METHOD_MAP = {
    "linear_concat": "Linear concat",
    "pca_add": "PCA add",
    "pca_then_concat": "PCA then concat",
    "fusion_mlp_interaction_unsup": "Interaction MLP",
    "fusion_lowrank_higher_order_unsup": "Low-rank higher-order",
    "fusion_reliability_experts_unsup": "Reliability experts",
    "fusion_cross_attention_unsup": "Cross-attention",
    "fusion_mlp_interaction_unsup_shuffled": "Interaction MLP (pair-shuffled)",
}
PROBE_LABEL = {"rf": "RF (frozen primary)", "linear": "Linear (logistic)",
               "mlp": "Supervised MLP head"}
TASK_ORDER = ["CCD Protein", "CCD Transcript", "N1 network", "N1 targets",
              "TF vs non-TF", "bivalent vs non-methylated", "bivalent vs lys4-only",
              "dosage sensitive vs insensitive TF", "long vs short range TF",
              "Blood expression cluster", "Brain expression cluster",
              "Cell line expression cluster",
              "Pathology prognostics - Breast cancer",
              "Pathology prognostics - Cervical cancer",
              "Pathology prognostics - Colorectal cancer"]
TASK_RENAME = {"bivalent_vs_lys4_only": "bivalent vs lys4-only"}
CONDS = ["Core", "Core+K", "Core+X", "Core+K+X"]


def load_run(run_dir: str) -> pd.DataFrame:
    rows = []
    for f in glob.glob(f"{run_dir}/task=*/modalities=*/methods=*/seed=*/metrics_long.csv"):
        parts = Path(f).parts
        cond_dir = [p for p in parts if p.startswith("modalities=")][0]
        df = pd.read_csv(f)
        df["condition"] = COND_MAP[cond_dir.split("=", 1)[1]]
        rows.append(df)
    if not rows:
        return pd.DataFrame()
    d = pd.concat(rows, ignore_index=True)
    d["method"] = d["representation_method"].map(METHOD_MAP).fillna(d["representation_method"])
    d["task"] = d["task"].replace(TASK_RENAME)
    return d


def task_equal_boot(piv: pd.DataFrame, n_boot: int = 10_000, seed: int = 20260918):
    rng = np.random.default_rng(seed)
    means = piv.mean(axis=0)
    boot = np.empty((n_boot, len(means)))
    for b in range(n_boot):
        idx = rng.integers(0, len(piv), len(piv))
        boot[b] = piv.to_numpy()[idx].mean(axis=0)
    lo, hi = np.percentile(boot, [2.5, 97.5], axis=0)
    out = pd.DataFrame({"mean": means, "lo": lo, "hi": hi})
    out.index = pd.MultiIndex.from_tuples(out.index, names=["condition", "method"])
    return out.reset_index()


def main():
    run_dir = sys.argv[1] if len(sys.argv) > 1 else \
        "${REPO_ROOT}/results/probe_track_20260918/probe_track_20260918"
    out_dir = Path(sys.argv[2]) if len(sys.argv) > 2 else Path(run_dir).parents[1] / "analysis"
    out_dir.mkdir(parents=True, exist_ok=True)

    d = load_run(run_dir)
    if d.empty:
        print("no metrics_long.csv found under", run_dir)
        return
    per_task = (d.groupby(["task", "condition", "method", "probe"], as_index=False)
                .agg(macro_f1=("macro_f1", "mean"),
                     balanced_accuracy=("balanced_accuracy", "mean"),
                     roc_auc=("roc_auc", "mean"),
                     macro_roc_auc=("macro_roc_auc", "mean"),
                     log_loss=("log_loss", "mean")))
    per_task["auc"] = per_task["roc_auc"].fillna(per_task["macro_roc_auc"])
    per_task.to_csv(out_dir / "probe_track_per_task.csv", index=False)

    summary_rows = []
    for probe in sorted(per_task["probe"].unique()):
        sub = per_task[per_task["probe"] == probe]
        tasks = [t for t in TASK_ORDER if t in set(sub["task"])]
        piv = sub[sub["task"].isin(tasks)].pivot_table(
            index="task", columns=["condition", "method"], values="macro_f1")
        te = task_equal_boot(piv)
        te["probe"] = probe
        te["n_tasks"] = len(tasks)
        summary_rows.append(te)
        print(f"\n=== probe={probe} | {len(tasks)} tasks ===")
        show = te.pivot_table(index="method", columns="condition", values="mean")
        print(show.reindex(columns=[c for c in CONDS if c in show.columns]).round(4))
        wins = {c: piv[c].mean(axis=0).idxmax() for c in CONDS if c in piv.columns.get_level_values(0)}
        print("top operator per condition:", wins)
        if "PCA then concat" in set(piv.columns.get_level_values("method")):
            m = piv.mean(axis=0).unstack("condition")
            print("PCA then concat: Delta_KX = {:+.4f}, N_KX = {:+.4f}".format(
                m.loc["PCA then concat", "Core+K+X"] - m.loc["PCA then concat", "Core"],
                m.loc["PCA then concat", "Core+K+X"] - m.loc["PCA then concat", "Core+K"]
                - m.loc["PCA then concat", "Core+X"] + m.loc["PCA then concat", "Core"]))
    pd.concat(summary_rows).to_csv(out_dir / "probe_track_task_equal.csv", index=False)
    print("\nwrote:", out_dir / "probe_track_task_equal.csv", "and probe_track_per_task.csv")


if __name__ == "__main__":
    main()
```
