## Supplementary_Material for "RepGene: Auditing Condition-Specific Fusion Biases for Multi-View Gene Representations": probe_track.yaml

```
project:
  name: RepGene
  version: '2026-09-18'
  run_id: probe_track_20260918
  protocol: probe-sensitivity-15task-4condition
data:
  task_root: ${REPO_ROOT}/data/gene-benchmark/tasks
  embedding_root: ${REPO_ROOT}/data/processed/embeddings/Homo_sapiens
  view_file_stems:
    DNA: DNA
    RNA: RNA
    protein: protein
    text: text
    scRNA: singlecell
benchmark:
  workflow: staged_unsupervised
  task_type: auto
  seeds:
  - 41
  - 42
  - 43
  cv_folds: 3
  tasks:
    include:
    - CCD Protein
    - CCD Transcript
    - N1 network
    - N1 targets
    - TF vs non-TF
    - bivalent vs non-methylated
    - bivalent_vs_lys4_only
    - dosage sensitive vs insensitive TF
    - long vs short range TF
    - Blood expression cluster
    - Brain expression cluster
    - Cell line expression cluster
    - Pathology prognostics - Breast cancer
    - Pathology prognostics - Cervical cancer
    - Pathology prognostics - Colorectal cancer
  modalities:
    available:
    - DNA
    - RNA
    - protein
    - text
    - scRNA
    groups:
      core:
      - DNA
      - RNA
      - protein
      core_plus_K:
      - DNA
      - RNA
      - protein
      - text
      core_plus_X:
      - DNA
      - RNA
      - protein
      - scRNA
      core_plus_K_plus_X:
      - DNA
      - RNA
      - protein
      - text
      - scRNA
    sets:
    - core
    - core_plus_K
    - core_plus_X
    - core_plus_K_plus_X
  methods:
    available:
    - linear_concat
    - pca_then_concat
    - pca_add
    - fusion_mlp_interaction_unsup
    - fusion_lowrank_higher_order_unsup
    - fusion_reliability_experts_unsup
    sets:
    - - linear_concat
      - pca_then_concat
      - pca_add
      - fusion_mlp_interaction_unsup
      - fusion_lowrank_higher_order_unsup
      - fusion_reliability_experts_unsup
  representation:
    protocol: fold_inductive
    target_dim: 256
    methods:
    - - linear_concat
      - pca_then_concat
      - pca_add
      - fusion_mlp_interaction_unsup
      - fusion_lowrank_higher_order_unsup
      - fusion_reliability_experts_unsup
    probes:
    - rf
    - linear
  model:
    hidden_dim: 96
    token_dim: 256
    dropout: 0.1
  training:
    epochs: 8
    batch_size: 1024
    learning_rate: 0.001
    weight_decay: 1.0e-05
  unsupervised:
    variance_weight: 0.1
    covariance_weight: 0.01
    alignment_weight: 0.05
    view_dropout_weight: 0.1
    availability_mode: core_k_x
    information_groups:
      core_views:
      - DNA
      - RNA
      - protein
      knowledge_views:
      - text
      context_views:
      - scRNA
  availability:
    condition_definitions:
      core:
      - DNA
      - RNA
      - protein
      core_plus_K:
      - DNA
      - RNA
      - protein
      - text
      core_plus_X:
      - DNA
      - RNA
      - protein
      - scRNA
      core_plus_K_plus_X:
      - DNA
      - RNA
      - protein
      - text
      - scRNA
  probe:
    logistic_max_iter: 2000
    rf_n_estimators: 300
    rf_max_depth: 16
    rf_min_samples_leaf: 2
    mlp_hidden_dim: 64
    mlp_dropout: 0.1
    mlp_learning_rate: 0.001
    mlp_weight_decay: 1.0e-05
    mlp_batch_size: 256
    mlp_epochs: 30
execution:
  output_root: results/probe_track_20260918
  device: cpu
  max_workers: 8
  skip_completed: true
  fail_fast: false
  save_predictions: true
  save_checkpoints: false
  cleanup_completed_artifacts: false
  cache:
    enabled: true
    directory: results/cache/probe_track_20260918
```
