## Supplementary_Material for "RepGene: Auditing Condition-Specific Fusion Biases for Multi-View Gene Representations": run_all_tracks.sh

```
#!/usr/bin/env bash
# Unified orchestrator for all supplementary tracks, single-threaded jobs with
# 8-way job parallelism on an 8-core container (avoids BLAS thread oversubscription).
# Order: probe sensitivity (B1) -> supervised MLP reference (B2) -> budget scan 32/64 -> shuffle control.
set -uo pipefail
cd ${REPO_ROOT}
export PYTHONPATH=${REPO_ROOT}/src
# one thread per job; parallelism comes from the runner's max_workers
export OMP_NUM_THREADS=1
export MKL_NUM_THREADS=1
export OPENBLAS_NUM_THREADS=1
export NUMEXPR_NUM_THREADS=1
export VECLIB_MAXIMUM_THREADS=1
LOG=experiments_20260918/orchestrator.log
echo "[$(date -u +%H:%M:%S)] orchestrator v2 start (single-thread jobs, 8-way)" >> "$LOG"

run_track () {
  local cfg="$1"; local rid="$2"
  echo "[$(date -u +%H:%M:%S)] starting $rid" >> "$LOG"
  python3 scripts/run_benchmark.py --config "experiments_20260918/configs/${cfg}" --run-id "$rid" >> "$LOG" 2>&1
  echo "[$(date -u +%H:%M:%S)] finished $rid exit=$?" >> "$LOG"
}

run_track probe_track.yaml probe_track_20260918
run_track supervised_ref.yaml supervised_ref_20260918
run_track budget_scan_epochs32.yaml budget_scan_epochs32_20260918
run_track budget_scan_epochs64.yaml budget_scan_epochs64_20260918
run_track shuffle_control.yaml shuffle_control_20260918

echo "[$(date -u +%H:%M:%S)] orchestrator v2 done" >> "$LOG"
```
