## Supplementary_Material for "RepGene: Auditing Condition-Specific Fusion Biases for Multi-View Gene Representations": run_probe_track.sh

```
#!/usr/bin/env bash
# Probe-sensitivity track (B1/B2): 15 tasks x 4 conditions x 3 seeds,
# three frozen probes evaluated on the same exports (rf, logistic, supervised MLP head).
set -euo pipefail
cd ${REPO_ROOT}
export PYTHONPATH=${REPO_ROOT}/src
python3 scripts/run_benchmark.py \
  --config experiments_20260918/configs/probe_track.yaml \
  --run-id probe_track_20260918
echo "PROBE_TRACK_EXIT=$?"
```
