## Supplementary_Material for "RepGene: Auditing Condition-Specific Fusion Biases for Multi-View Gene Representations": run_remaining_tracks.sh

```
#!/usr/bin/env bash
# Orchestrator: run the remaining supplementary tracks sequentially after the
# probe track finishes, so the 8-core container is never oversubscribed.
# Tracks: budget scan (epochs 32, 64) -> view-pair shuffle control.
set -uo pipefail
cd ${REPO_ROOT}
export PYTHONPATH=${REPO_ROOT}/src
LOG=experiments_20260918/orchestrator.log
echo "[$(date -u +%H:%M:%S)] orchestrator start" >> "$LOG"

# 1) wait for the probe-sensitivity track to finish
while pgrep -f "probe_track.yaml" > /dev/null 2>&1; do
  sleep 60
done
echo "[$(date -u +%H:%M:%S)] probe track finished" >> "$LOG"

run_track () {
  local cfg="$1"; local rid="$2"
  echo "[$(date -u +%H:%M:%S)] starting $rid" >> "$LOG"
  python3 scripts/run_benchmark.py --config "experiments_20260918/configs/${cfg}" --run-id "$rid" >> "$LOG" 2>&1
  echo "[$(date -u +%H:%M:%S)] finished $rid exit=$?" >> "$LOG"
}

run_track budget_scan_epochs32.yaml budget_scan_epochs32_20260918
run_track budget_scan_epochs64.yaml budget_scan_epochs64_20260918
run_track shuffle_control.yaml shuffle_control_20260918

echo "[$(date -u +%H:%M:%S)] orchestrator done" >> "$LOG"
```
