## Supplementary_Material for "RepGene: Auditing Condition-Specific Fusion Biases for Multi-View Gene Representations": smoke_runner.sh

```
#!/usr/bin/env bash
# Smoke test: single task, single condition, single seed, three probes.
# Verifies the Pod pipeline reproduces the frozen contract numbers before the
# full supplementary experiment matrix is launched.
set -euo pipefail
cd ${REPO_ROOT}
export PYTHONPATH=${REPO_ROOT}/src
python3 scripts/run_benchmark.py \
  --config experiments_20260918/configs/smoke_probe.yaml \
  --run-id smoke_probe_20260918
echo "SMOKE_EXIT=$?"
```
