## Supplementary_Material for "RepGene: Auditing Condition-Specific Fusion Biases for Multi-View Gene Representations": make_paper_tables.py

```
"""Generate the LaTeX tables for the protocol-extension appendix section.

Inputs (produced by the analysis scripts):
  results/analysis/probe_track_task_equal.csv   probe sensitivity, 15 tasks, 2 probes
  results/analysis/budget_scan_per_task.csv     epochs 8/32/64 on binary9 x Core+K+X
  results/analysis/shuffle_control_per_task.csv view-pair shuffle control
  results/analysis/q6_calibration_summary.csv   calibration / threshold sweep
  ratio of the supervised MLP-head reference    from the supervised_ref run
Outputs: tables/gen/*.tex, included by the new appendix section.
"""
from __future__ import annotations

import glob
from pathlib import Path

import pandas as pd

REPO = Path("${REPO_ROOT}")
OUT = REPO / "docs/paper/ICLR2027/outputs/RepGene-ICLR2027-0918-1000/tables/gen"
OUT.mkdir(parents=True, exist_ok=True)
AN = REPO / "results/analysis"

MNAME = {
    "Linear concat": "Linear concat",
    "PCA add": "PCA add",
    "PCA then concat": "PCA then concat",
    "Interaction MLP": "Interaction MLP",
    "Low-rank higher-order": "Low-rank higher-order",
    "Reliability experts": "Reliability experts",
    "Cross-attention": "Cross-attention",
}
ORDER = ["Linear concat", "PCA add", "PCA then concat", "Interaction MLP",
         "Low-rank higher-order", "Reliability experts"]
CONDS = ["Core", "Core+K", "Core+X", "Core+K+X"]


def f4(x):
    return "--" if pd.isna(x) else f"{x:.4f}"


def table_probe_sensitivity() -> str:
    d = pd.read_csv(AN / "probe_track_task_equal.csv")
    lines = []
    for probe, label in [("rf", "RF probe (frozen primary)"), ("linear", "linear (logistic) probe")]:
        sub = d[d.probe == probe]
        piv = sub.pivot_table(index="method", columns="condition", values="mean")
        lo = sub.pivot_table(index="method", columns="condition", values="lo")
        hi = sub.pivot_table(index="method", columns="condition", values="hi")
        lines.append(f"\\multicolumn{{5}}{{l}}{{\\textit{{{label}}}}} \\\\")
        lines.append("\\midrule")
        best_by_cond = {c: piv[c].idxmax() for c in piv.columns}
        for m in ORDER:
            if m not in piv.index:
                continue
            cells = []
            for c in CONDS:
                if c not in piv.columns:
                    cells.append("--")
                    continue
                if best_by_cond.get(c) == m:
                    cells.append(f"$\\mathbf{{{piv.loc[m, c]:.4f}}}$")
                else:
                    cells.append(f"${piv.loc[m, c]:.4f}$")
            lines.append(f"{m} & " + " & ".join(cells) + " \\\\")
        lines.append("\\midrule")
    body = "\n".join(lines)
    body = body.replace("\\midrule\n\\midrule", "\\midrule")
    # drop a single trailing separator line, keeping row terminators intact
    rows = [ln for ln in body.split("\n")]
    while rows and rows[-1].strip() in {"", "\\midrule"}:
        rows.pop()
    return "\n".join(rows)


def table_budget() -> str:
    d = pd.read_csv(AN / "budget_scan_per_task.csv")
    m = d[d.env == "matched"]
    piv = m.pivot_table(index="method", columns="epochs", values="macro_f1", aggfunc="mean")
    ref = d[d.env == "frozen-ref"].pivot_table(index="method", columns="epochs", values="macro_f1", aggfunc="mean")
    rows = [
        ("Reliability experts", "Reliability experts"),
        ("Interaction MLP", "Interaction MLP"),
        ("Low-rank higher-order", "Low-rank higher-order"),
        ("Cross-attention", "Cross-attention"),
        ("PCA then concat", "PCA then concat"),
        ("Linear concat", "Linear concat"),
    ]
    lines = []
    for disp, key in rows:
        v8m = piv.loc[key, 8] if key in piv.index else float("nan")
        v32 = piv.loc[key, 32] if key in piv.index else float("nan")
        v64 = piv.loc[key, 64] if key in piv.index else float("nan")
        v8f = ref.loc[key, 8] if key in ref.index else float("nan")
        lines.append(f"{disp} & {f4(v8m)} & {f4(v32)} & {f4(v64)} & {f4(v8f)} \\\\")
    return "\n".join(lines)


def table_shuffle() -> str:
    d = pd.read_csv(AN / "shuffle_control_per_task.csv")
    piv = d.pivot_table(index="method", columns="condition", values="macro_f1")
    real = piv.loc["Interaction MLP"]
    shuf = piv.loc["Interaction MLP (pair-shuffled)"]
    l1 = "Real pairing & " + " & ".join(f"{real[c]:.4f}" for c in CONDS) + " \\\\"
    l2 = "Pair-shuffled twin & " + " & ".join(f"{shuf[c]:.4f}" for c in CONDS) + " \\\\"
    dl = "Difference & " + " & ".join(f"$+{real[c]-shuf[c]:.4f}$" for c in CONDS) + " \\\\"
    return "\n".join([l1, l2, "\\midrule", dl])


def table_calibration() -> str:
    d = pd.read_csv(AN / "q6_calibration_summary.csv")
    lines = []
    for probe, label in [("rf", "RF probe"), ("linear", "linear probe")]:
        sub = d[d.probe == probe].set_index("method_key")
        lines.append(f"\\multicolumn{{6}}{{l}}{{\\textit{{{label}}}}} \\\\")
        lines.append("\\midrule")
        for key, disp in [("fusion_reliability_experts_unsup", "Reliability experts"),
                          ("fusion_mlp_interaction_unsup", "Interaction MLP"),
                          ("fusion_lowrank_higher_order_unsup", "Low-rank higher-order"),
                          ("pca_then_concat", "PCA then concat"),
                          ("linear_concat", "Linear concat"),
                          ("pca_add", "PCA add")]:
            r = sub.loc[key]
            lines.append("{} & {} & {} & {} & {} & {} \\\\".format(
                disp, f4(r.macro_f1_default), f4(r.macro_f1_best_threshold), f4(r.brier), f4(r.ece), f4(r.auc)))
        lines.append("\\midrule")
    rows = [ln for ln in "\n".join(lines).split("\n")]
    while rows and rows[-1].strip() in {"", "\\midrule"}:
        rows.pop()
    return "\n".join(rows)


def table_supervised_ref() -> str:
    rows = []
    for f in glob.glob(str(REPO / "results/supervised_ref_20260918/supervised_ref_20260918/task=*/modalities=*/methods=*/seed=*/metrics_long.csv")):
        rows.append(pd.read_csv(f))
    d = pd.concat(rows)
    piv = d.pivot_table(index="representation_method", columns="probe", values="macro_f1", aggfunc="mean")
    lines = []
    for key, disp in [("linear_concat", "Linear concat"), ("pca_then_concat", "PCA then concat"),
                      ("fusion_lowrank_higher_order_unsup", "Low-rank higher-order"),
                      ("fusion_mlp_interaction_unsup", "Interaction MLP"),
                      ("fusion_reliability_experts_unsup", "Reliability experts"),
                      ("pca_add", "PCA add")]:
        lines.append(f"{disp} & {piv.loc[key,'mlp']:.4f} & {piv.loc[key,'rf']:.4f} \\\\")
    return "\n".join(lines)


FLOATS = {
    "tab_probe_sensitivity": ("Fifteen-task task-equal Macro-F1 under two probes evaluated on identical "
        "exported representations (three seeds, three folds, four conditions). Boldface marks the "
        "per-condition leader. Bootstrap intervals of width $\\approx\\pm0.13$ overlap throughout, as in "
        "the main register.", "tab:probe-sensitivity", "lrrrr",
        "Method & Core & Core+K & Core+X & Core+K+X \\\\", "probe_sensitivity_full15"),
    "tab_supervised_ref": ("Supervised MLP-head reference (B2) on the nine binary tasks under Core+K+X, "
        "alongside the frozen random-forest probe on the same exported representations. The trained head "
        "does not restore the learned experts' advantage over the concatenation baselines.",
        "tab:supervised-ref", "lrr", "Method & Supervised MLP head & RF probe \\\\", "supervised_ref_binary9"),
    "tab_budget": ("Nine binary tasks under Core+K+X, task-equal Macro-F1 against the label-free training "
        "budget. Columns 2--3 are the matched re-execution; the last column is the frozen epoch-8 reference "
        "for drift comparison. PCA then concat and Linear concat were not re-run at higher budgets (the "
        "budget question concerns the learned operators) and are shown for the epoch-8 reference only.",
        "tab:budget", "lrrrr", "Method & 8 epochs & 32 epochs & 64 epochs & 8 epochs (frozen ref.) \\\\",
        "budget_scan_binary9"),
    "tab_shuffle": ("View-pair shuffle control on the nine binary tasks (three seeds, four conditions, "
        "random-forest probe). The shuffled twin is capacity-matched; only the gene-wise cross-view pairing "
        "is destroyed.", "tab:shuffle", "lrrrr", "Variant & Core & Core+K & Core+X & Core+K+X \\\\",
        "shuffle_control_binary9"),
    "tab_calibration": ("Calibration and threshold sensitivity on the nine binary tasks, task-equal means. "
        "Macro-F1 is reported at the frozen $0.5$ threshold and at the best swept threshold of the same "
        "probabilities; Brier and ECE quantify calibration; ROC-AUC quantifies rank discrimination.",
        "tab:calibration", "lrrrrr", "Method & F1@$0.5$ & F1@best & Brier & ECE & AUC \\\\",
        "calibration_binary9"),
}


def write_float(name, caption, label, colspec, header, fragment):
    body = (Path(OUT) / f"{fragment}.tex").read_text().rstrip("\n")
    tex = ("\\begin{table}[h]\n\\centering\\scriptsize\n"
           f"\\caption{{{caption}}}\n\\label{{{label}}}\n"
           f"\\begin{{tabular}}{{{colspec}}}\n\\toprule\n{header.replace(chr(92)*4, chr(92)*2)}\n"
           "\\midrule\n" + body + "\n\\bottomrule\n\\end{tabular}\n\\end{table}\n")
    (Path(OUT) / f"{name}.tex").write_text(tex)


if __name__ == "__main__":
    (OUT / "probe_sensitivity_full15.tex").write_text(table_probe_sensitivity() + "\n")
    (OUT / "budget_scan_binary9.tex").write_text(table_budget() + "\n")
    (OUT / "shuffle_control_binary9.tex").write_text(table_shuffle() + "\n")
    (OUT / "calibration_binary9.tex").write_text(table_calibration() + "\n")
    (OUT / "supervised_ref_binary9.tex").write_text(table_supervised_ref() + "\n")
    for name, spec in FLOATS.items():
        write_float(name, *spec)
    for f in sorted(OUT.glob("*.tex")):
        if f.name in {"probe_sensitivity_full15.tex", "budget_scan_binary9.tex", "shuffle_control_binary9.tex",
                      "calibration_binary9.tex", "supervised_ref_binary9.tex"}:
            print("----", f.name)
            print(f.read_text())
```
