## Supplementary_Material for "RepGene: Auditing Condition-Specific Fusion Biases for Multi-View Gene Representations": README.md

# Round-2 protocol extensions (2026-09-18)

All five tracks were executed by re-running the released pipeline (`scripts/run_benchmark.py`)
in the cloud task Pod with the released configuration files as the only input.

| Run id | Coverage | Cells | Probes |
|---|---|---|---|
| `probe_track_20260918` | 15 tasks x 4 conditions x 3 seeds x 6 operators | 180 | RF + logistic |
| `supervised_ref_20260918` | 9 binary x Core+K+X x 3 seeds | 27 | RF + supervised MLP head |
| `budget_scan_epochs32_20260918` | 9 binary x Core+K+X x 3 seeds x 4 learned operators | 27 | RF |
| `budget_scan_epochs64_20260918` | same, 64 epochs | 27 | RF |
| `shuffle_control_20260918` | 9 binary x 4 conditions x 3 seeds x 2 variants | 108 | RF |

Result roots live under `results/` in the ${REPO_ROOT} repository; `Per-gene` probabilities were
dumped for every fold (720 task x condition x method x probe cells in the probe track).

## Fidelity check performed before use
- per-fold train/test `train_index_sha256` / `test_index_sha256` identical to the frozen register;
- sample counts identical (`n_train` 925/926, `n_test` 463 on the smoke cell);
- epoch-8 task-equal means on binary9 x Core+K+X agree with the frozen values to four decimals
  (Reliability experts 0.6995 vs 0.6993; Interaction MLP 0.6965 vs 0.6963; PCA then concat
  0.7037 vs 0.7042).

## Environment difference (documented in the paper)
Rebuild environment: scikit-learn 1.9.0, numpy 2.x, torch 2.13.0+cpu; freeze environment:
numpy 1.26.4, torch 2.1.0+cu121. Comparisons inside a track are matched; the cross-environment
comparison of the epoch-8 reference is reported with this caveat.

## Code change
`mlp_interaction_shuffled` was added as a new structure (${REPO_ROOT} commit `2d0540f`), with
identical parameter count and identical concatenation block; only the gene-wise pairing of the
pair terms is destroyed (row-permuted second view, fixed permutation seed 20260918).
