## Supplementary_Material for "RepGene: Auditing Condition-Specific Fusion Biases for Multi-View Gene Representations": cell_boundary_clustering.pdf

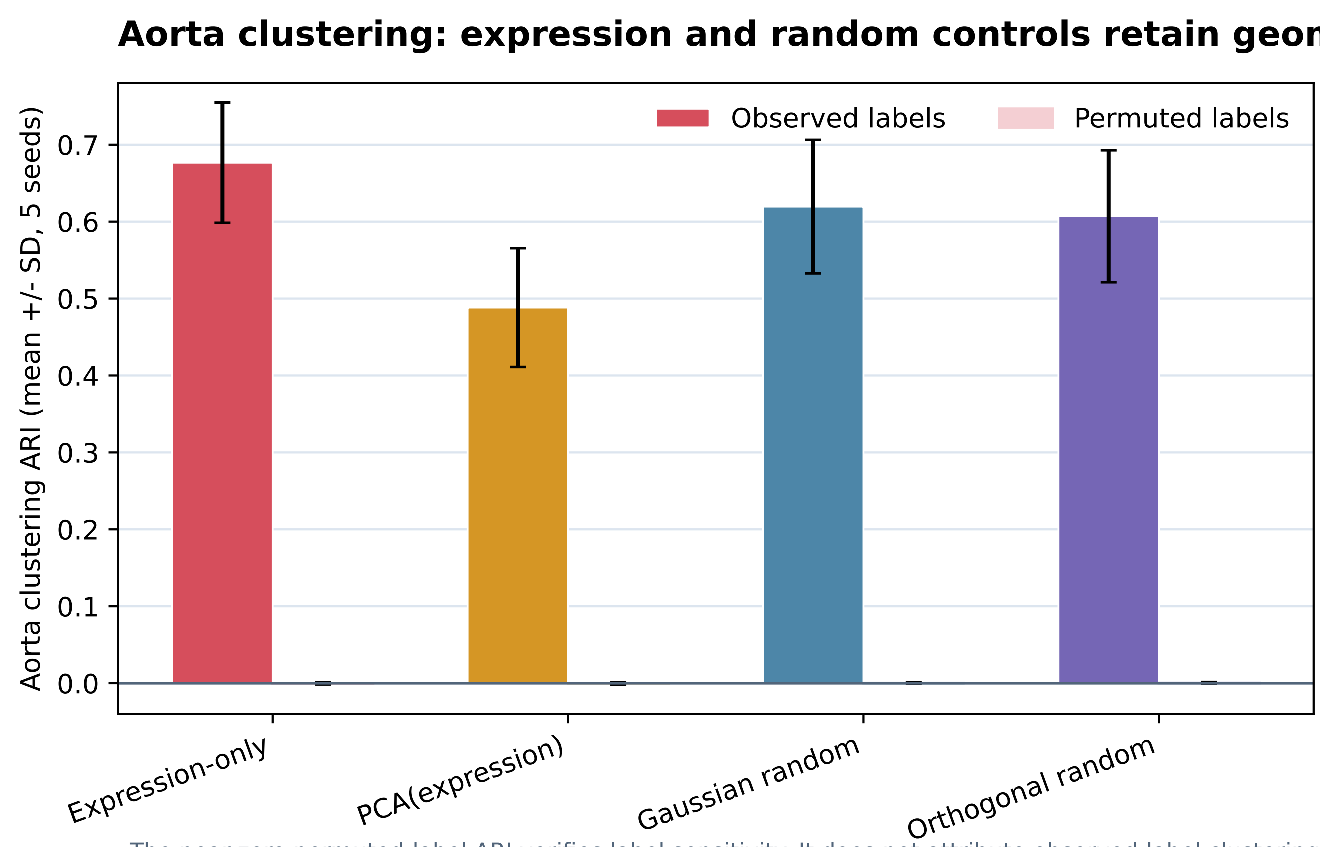

The near-zero permuted-label ARI verifies label sensitivity. It does not attribute observed-label clustering to multimodal semantic information.
