## Supplementary_Material for "RepGene: Auditing Condition-Specific Fusion Biases for Multi-View Gene Representations": cell_boundary_knn.pdf

### Cell annotation confound controls: observed versus independently permuted labels

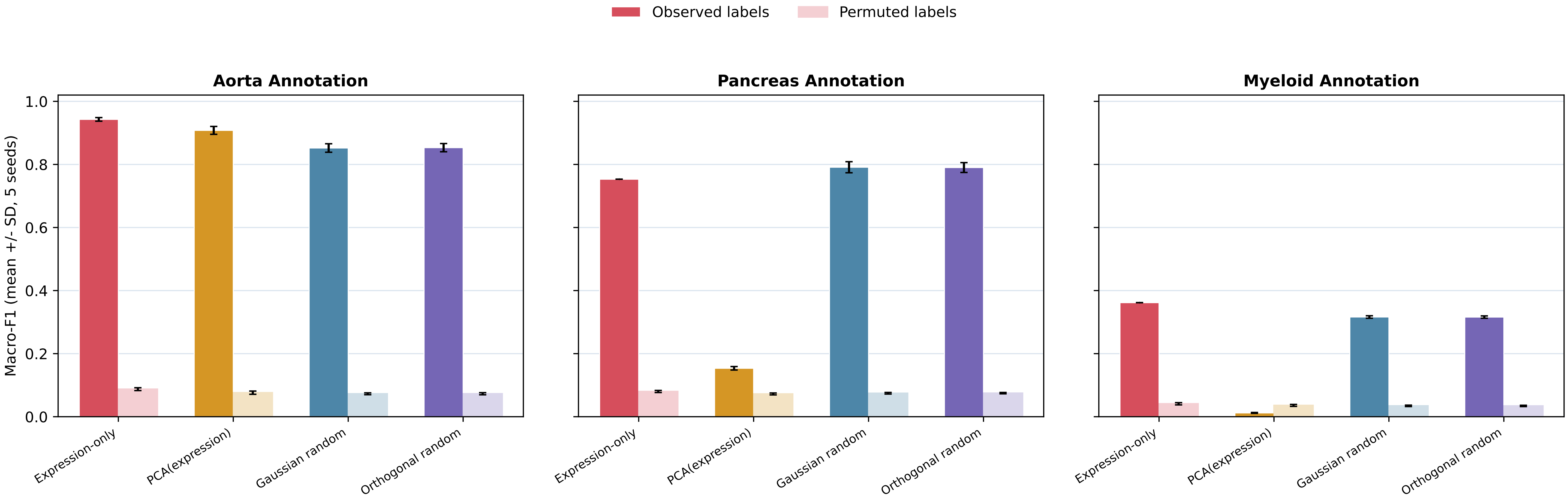


Controlled library-size aggregation, cell-wise L2 normalization, and controlled logistic regression. Strong observed-label scores for expression and random controls are not direct evidence of multimodal gene semantics.
