## Supplementary_Material for "RepGene: Auditing Condition-Specific Fusion Biases for Multi-View Gene Representations": supplementary_material.pdf

### REPGENE: SUPPLEMENTARY MATERIAL
#### ARTIFACT, REPRODUCTION, AND RELEASED-RESULT DOCUMENTATION

**Anonymous authors**
Paper under double-blind review

#### PURPOSE AND SCOPE

This document is the supplementary material for the submitted RepGene paper. It is restricted to three things that the Main Paper does not carry in full: (i) an inventory of the submitted artifact and the role of each file; (ii) a complete, file-level account of every released result table, together with the manuscript location it supports; and (iii) the reproduction order and the code/data access information. The scientific definitions, the complete appendix tables, the statistical procedures, the limitations, and the supplementary figures remain in the Main Paper PDF and its Appendix; they are not duplicated here. No new experiments, datasets, or numerical claims are introduced in this file: every number quoted below is read off a released table or a released manuscript table fragment that is included in the archive.

#### 1 SUBMITTED FILE STRUCTURE

The submission consists of three artifacts with deliberately different roles.

- **Main Paper PDF** (`RepGene_ICLR2027_0921_1740.pdf`): the complete paper, including the nine-page main text, references, the Impact / Reproducibility / Ethics statements, and the numbered Appendix. The Appendix is the primary location of the complete result tables and of the scientific discussion of them.
- **Supplementary PDF** (`supplementary_material.pdf`): this document only. It documents the artifact, the released tables, and the reproduction procedure. It does not contain the Appendix and does not replace it.
- **Anonymous source archive** (`RepGene_ICLR2027_Supplementary_Material.zip`): the LaTeX sources, the figure assets, the released result tables, the analysis scripts and configuration files, and this supplementary source file. The archive is the artifact that makes the reported measurements inspectable and regenerable.

#### 2 ARTIFACT INVENTORY

Table 1 lists the top-level components of the source archive and the role of each one. All paths are relative to the archive root, which is also the compilation directory of the manuscript.

Table 1: Top-level inventory of the anonymous source archive.

| Component | Role |
|---|---|
| main.tex | Manuscript entry point; inputs the section files and the released table fragments. |
| sections/ | Manuscript and appendix sources (abstract, introduction, relatedwork, approach, experiment, conclusion, impactstatement, othes, and the appendix files appendix, appendix_embedding, appendix_extension, appendix_suppfigures). |
| references.bib | Bibliography database. |
| iclr2027_conference.sty, iclr2027_conference.bst, auxiliary style files | Conference style and bibliography style; bundled so that the manuscript compiles without a network fetch. |
| figures/current/ | Figure assets referenced by the manuscript (PDF) and their editable sources (SVG/PNG). |
| figures/large_size_transform/ | Large-format embedding panels (UMAP/t-SNE master panels) used for the embedding-geometry analysis. |
| tables/ | **All released result tables** (CSV), plus the LaTeX table fragments used by the manuscript under tables/gen/. |
| analysis/round2/ | Scripts, configuration files, and result CSVs of the four registered protocol extensions. |
| supplementary_material.tex | Source of this document. |
| REPRODUCE.md | Short build-and-rerun note. |

#### 3 RELEASED RESULT TABLES: COMPLETE INDEX

Table 2 is the complete file-level index of the released result tables. The "register" column states the evaluation set: *full15* is the fifteen-task register (Table 1 of the Main Paper), *binary9* is the nine binary tasks of the sub-register, and *external* is the separately configured random-forest track. "Manuscript location" points to the Main Paper element that the file supports.

Table 2: Complete index of the released result tables in tables/. Every file is a plain UTF-8 CSV with a header row.

| File (tables/) | Register | Content | Manuscript location |
|---|---|---|---|
| F1_plotted.csv | — | Panel/question/element map behind the four-question evaluation figure (4 rows). | Figure 1 |
| F2_plotted.csv | full15 | Task-equal Macro-F1 by condition and operator with descriptive 95% intervals (24 rows: 4 conditions $\times$ 6 operators). | Figure 2 |
| F3_plotted.csv | full15 | Task $\times$ condition $\times$ view-set $\times$ operator long table (216 rows, accuracy and per-view diagnostics) behind the condition heatmap. | Figure 3 |
| F4_plotted.csv | full15 | Per-task operator scores under Core, Core+K, Core+X, Core+K+X (54 rows) behind the paired-condition-effect figure. | Figure 4 |
| F5_plotted.csv | full15 | Mean rank and per-condition win counts by operator (24 rows). | Figure 5 |
| F6_plotted.csv | binary9 | Same schema as F2_plotted.csv for the nine binary tasks. | Figure 6 |
| F7_plotted.csv | full15 | Per-operator task-equal means and intervals for the non-additivity term. | Figure 7 |
| F8_plotted.csv | external | External-track display values with intervals (9 rows). | Figure 8 |

| File (`tables/`) | Register | Content | Manuscript location |
|---|---|---|---|
| `full15_task_equal_summary_rf.csv` | full15 | Task-equal Macro-F1 mean/lo/hi per condition × operator under the frozen RF probe (24 rows). | Section 4, Table 4 |
| `full15_paired_condition_effects_rf.csv` | full15 | Matched condition deltas $\Delta_K$, $\Delta_X$, $\Delta_{KX}$ and the non-additivity term $N_{KX}$ per operator (6 rows). | Section 4, Table 4 |
| `full15_best_method_by_task_condition.csv` | full15 | Best-scoring operator per task and condition with the within-task spread (15 rows). | Appendix per-task tables |
| `full15_task_condition_method_long.csv` | full15 | Complete task × condition × operator records with Macro-F1, balanced accuracy, ROC-AUC and macro ROC-AUC (360 rows). | Appendix per-task tables |
| `binary9_task_equal_summary_rf.csv` | binary9 | Task-equal summaries by condition and operator (24 rows). | Appendix, binary diagnostics |
| `binary9_paired_condition_effects_rf.csv` | binary9 | Matched condition deltas on the sub-register (6 rows). | Appendix, binary diagnostics |
| `binary_task_equal_summary_rf.csv` | binary9 | Task-equal summaries including view-set splits (26 rows). | Appendix, binary diagnostics |
| `binary_paired_condition_effects_rf.csv` | binary9 | Per-task condition contrasts including the two attention operators (72 rows). | Appendix, attention operators |
| `binary_best_method_by_task_condition.csv` | binary9 | Best operator per task and condition (36 rows). | Appendix, attention operators |
| `binary_task_condition_method_summary.csv` | binary9 | Per-task, per-condition operator summary with accuracy dispersion (234 rows). | Appendix, attention operators |
| `family_stratified_task_equal_rf.csv` | full15 | Task-equal Macro-F1 stratified by task family, per condition and operator (96 rows). | Appendix, Table 17 |
| `leakage_control_condition_effects_rf.csv` | full15 | Condition contrasts with and without the five tasks whose labels share semantic overlap with a view (6 rows). | Appendix, leakage control |
| `leakage_control_delta_by_group_rf.csv` | full15 | The same contrasts split into overlap and non-overlap task groups (6 rows). | Appendix, leakage control |
| `paired_task_deltas.csv` | full15 | Paired per-task deltas between operator pairs with the common-gene count (27 rows). | Section 4 / Appendix |
| `table5_task_equal_source.csv` | binary9 | Source table of the external comparison: GenePT text-only baseline, PRISME+scRNA extension, RepGene Linear concat and Reliability experts, with paired deltas and intervals (4 rows). | Section 4 comparison table |
| `task_method_heatmap_source.csv` | full15 | Task-type-labelled heatmap source with condition codes (99 rows). | Figure 3 / Appendix |
| `new_methods_binary_results_long.csv` | binary9 | Fold-level records for the operators added in this revision across protocol variants (162 rows, 26 columns; includes fold, seed, probe, views, task type and protocol identifiers). | Appendix, attention operators |

#### 4 RELEASED RESULT TABLES: DETAIL

This section states, family by family, what each released table contains and which released values it carries. The numbers below are transcriptions of the released files; they are reported here so that the manuscript's tables can be checked against the tables that produced them.

##### 4.1 Fifteen-task register summaries

full15_task_equal_summary_rf.csv (24 rows) reports the task-equal Macro-F1 mean and a descriptive interval for each condition × operator cell, and is the source of the main condition table. Under the frozen random-forest probe the Core+K+X means are 0.5188 (PCA then concat, per-condition leader), 0.5142 (Reliability experts), 0.5122 (Low-rank higher-order), 0.5112 (Interaction MLP), 0.4586 (Linear concat) and 0.3765 (PCA add); the corresponding Core means are 0.4560, 0.4962, 0.4895, 0.4943, 0.4358 and 0.3832. The recorded intervals are approximately ±0.13 wide and overlap in every condition, which is why the manuscript reports the ordering inside the register rather than a winner. full15_paired_condition_effects_rf.csv (6 rows) carries the matched decomposition that the manuscript table uses: for each operator the Core→Core+K delta $\Delta_K$, the Core→Core+X delta $\Delta_X$, the joint delta $\Delta_{KX}$ and the non-additivity term $N_{KX}$. The released values are $+0.0332/+0.0492/+0.0629/-0.0195$ (PCA then concat), $+0.0086/+0.0061/+0.0169/+0.0022$ (Interaction MLP), $+0.0149/+0.0130/+0.0227/-0.0051$ (Low-rank higher-order), $+0.0087/+0.0062/+0.0179/+0.0030$ (Reliability experts), $+0.0088/+0.0189/+0.0228/-0.0049$ (Linear concat) and $-0.0114/+0.0014/-0.0067/+0.0034$ (PCA add). These are descriptive decompositions of matched runs, not causal effects or significance tests. full15_task_condition_method_long.csv (360 rows) is the complete task-level record and additionally reports balanced accuracy, ROC-AUC and macro ROC-AUC; full15_best_method_by_task_condition.csv (15 rows) records the best operator per task and condition with the within-task spread; task_method_heatmap_source.csv (99 rows) is the task-type-labelled condition heatmap source. Secondary metrics are released in tables/gen/secondary_full15.tex: task-equal balanced accuracy stays between 0.050 and 0.527 across operators and conditions, whereas ROC-AUC spans 0.6648 to 0.7894, with PCA then concat highest under Core+K+X (0.7894).

##### 4.2 Binary sub-register summaries and attention operators

binary9_task_equal_summary_rf.csv (24 rows) and binary_task_equal_summary_rf.csv (26 rows) hold the task-equal summaries of the nine binary tasks, the second broken out by view set; binary9_paired_condition_effects_rf.csv (6 rows) and binary_paired_condition_effects_rf.csv (72 rows) hold the matched condition contrasts, the latter including the two additional attention operators; binary_task_condition_method_summary.csv (234 rows) and binary_best_method_by_task_condition.csv (36 rows) give the per-task and per-condition detail. new_methods_binary_results_long.csv (162 rows, 26 columns) is the fold-level record for the operators added in this revision, carrying fold, seed, probe, views, task_type and protocol identifiers so that every reported cell can be traced to its fold.

##### 4.3 Leakage and task-family controls

leakage_control_condition_effects_rf.csv (6 rows) recomputes the condition contrasts with and without the five tasks whose labels share semantic overlap with one of the views. The released values show the ordering is preserved: for PCA then concat $\Delta_{KX}$ is $+0.0629$ on the full register and $+0.0600$ without the overlap group ($N_{KX}$: $-0.0195$ and $-0.0214$); for Linear concat $+0.0228$ and $+0.0168$; the values for the remaining operators are in the file. leakage_control_delta_by_group_rf.csv (6 rows) repeats the contrasts separately for the overlap group (5 tasks) and the non-overlap group (10 tasks). family_stratified_task_equal_rf.csv (96 rows) is the family-stratified source table; the corresponding manuscript fragment (tables/gen/family_table.tex) reports, per family and condition, the top operator and its task-equal Macro-F1, e.g. complex/network (0.5906 → 0.6811 from Core to Core+K+X) and TF/chromatin (0.7777 → 0.7951).

##### 4.4 Paired task deltas and external comparators

paired_task_deltas.csv (27 rows) lists, for each task and condition, the two operators being compared, the number of common genes, both task scores and the delta; it is the per-task backing of the paired statements in the manuscript. table5_task_equal_source.csv (4 rows) is the source of the external comparison table and carries, for the nine binary tasks: GenePT text-only baseline 0.5479, RepGene Linear concat 0.6342, PRISME+scRNA protocol extension 0.7116 and

RepGene Reliability experts 0.7227, together with the paired deltas against GenePT and PRISME and their intervals.

##### 4.5 Registered protocol extensions

The four registered extensions are stored under `analysis/round2/` together with their configuration files and the manuscript fragments in `tables/gen/`. The summary fragment `tables/gen/tab_extensions.tex` states the outcome of each: (a) *probe family* — replacing the forest with a logistic probe on identical exports leaves the structure intact (no operator leads in all four conditions; PCA then concat stays least decomposable with $N_{KX} = -0.0204$ under the forest and $-0.0116$ under the linear probe) but the leader changes and the experts' margin over Linear concat does not survive (0.4929 vs. 0.4366 becomes 0.4964 vs. 0.5057); (b) *supervised head* — a trained MLP head on the same exports gives Linear concat 0.7523, ahead of the learned experts (0.7257–0.7326); (c) *training budget* — every learned operator improves from 8 to 32 and 64 epochs ($\approx +0.022$ and $+0.031$) yet cross-attention never leads; (d) *pairing control* — the capacity-matched pair-shuffled twin loses in all four conditions (paired mean $+0.0259$, median $+0.0212$, 94% of 32 cells positive). The per-track tables are released as `tables/gen/tab_probe_sensitivity.tex` (full fifteen-task grid under both probes), `tables/gen/tab_supervised_ref.tex`, `tables/gen/tab_budget.tex`, `tables/gen/tab_shuffle.tex` and `tables/gen/tab_calibration.tex`. The corresponding per-run records are `probe_track_per_task.csv` (720 cells) and `probe_track_task_equal.csv` (48 rows) for the probe family, `budget_scan_per_task.csv` (176 rows) for the budget scan, `shuffle_control_per_task.csv` (64 rows) and `shuffle_control_paired.csv` (32 rows) for the pairing control, and `q6_calibration_summary.csv` (12 rows) for the calibration analysis. The calibration record shows the inversion discussed in the manuscript: PCA then concat has the best rank discrimination and the worst calibration under the forest probe (AUC 0.9387, ECE 0.1270; Macro-F1 0.6866 at the frozen 0.5 threshold and 0.7708 at the best swept threshold).

##### 4.6 LaTeX table fragments used by the manuscript

`tables/gen/` holds the fragments that the manuscript and appendix input directly, so that every reported table is byte-identical to the released one. They are: `pertask_core_full15.tex`, `pertask_corek_full15.tex`, `pertask_corex_full15.tex`, `pertask_corekx_full15.tex`, `best_by_task_full15.tex`, `family_table.tex`, `leakage_control_table.tex`, `condition_effects_binary9.tex`, `secondary_full15.tex`, `tab_condition_means.tex`, `tab_condition_effects.tex`, `tab_external_track.tex`, `tab_cell_boundary.tex`, `tab_extensions.tex`, `tab_probe_sensitivity.tex`, `tab_supervised_ref.tex`, `tab_budget.tex`, `tab_shuffle.tex` and `tab_calibration.tex`. Three of them carry results that the manuscript quotes directly: the mask-weighted condition table (`tab_condition_means`: Core+K+X leaders 0.5188 for PCA then concat and 0.5142 for Reliability experts), the matched condition deltas (`tab_condition_effects`) and the external track (`tab_external_track`: Low-rank higher-order 0.7251 with interval $[0.63, 0.81]$, GenePT text-only 0.5479, random 256-D control 0.4240).

#### 5 Figure inventory

`figures/current/` contains the figure assets and their editable sources. The files referenced directly by the manuscript are: `Figure_1_four_questions.pdf` (four-question evaluation map), `Figure_2_task_equal_condition_curves.pdf` (binary sub-register diagnostic of the condition curves), `Figure_3_task_method_condition_heatmap.pdf`, `Figure_4_paired_condition_effects.pdf`, `Figure_5_method_rank_and_wins.pdf`, `Figure_6_binary_suite_profiles.pdf`, `Figure_7_KX_nonadditivity.pdf`, `Figure_8_external_benchmark.pdf`, and the embedding-geometry set `emb_cka_matrix_all_methods.pdf`, `emb_cross_view_cka_heatmap.pdf`, `emb_embedding_stats_all.pdf`, `emb_knn_overlap_k10_heatmap.pdf`, `emb_pca_variance_explained.pdf`, `cell_boundary_clustering.pdf` and `cell_boundary_knn.pdf`.

The vector (SVG) and raster (PNG) sources of the main-text figures are shipped alongside the typeset PDFs so that the figures can be edited without regenerating the underlying measurements. `figures/large_size_transform/` holds the large-format embedding panels (`emb_umap_master_panel_01.pdf`, `emb_tsne_master_panel_01.pdf`, `emb_native_umap_panel_01.pdf`, `emb_fusion_umap_comparison_01.pdf`, `fusion_umap_comparison_01.pdf`) used by the embedding-geometry appendix. The manuscript's Reproducibility Statement refers to the t-SNE master panel; the corresponding file in this archive is `emb_tsne_master_panel_01.pdf`.

#### 6 Reproduction instructions

The manuscript is compiled from the archive root, which is the directory that contains `main.tex`:

```
latexmk -cd -pdf -interaction=nonstopmode -halt-on-error main.tex
# or, without a system TeX Live installation:
tectonic -X compile main.tex
```

The same command compiles this document from `supplementary_material.tex`. The analysis scripts under `analysis/` and `analysis/round2/` are written against the repository variable `${REPO_ROOT}`; set it to the archive root before re-running them. Nothing in the figure or table pipeline requires re-running a learned model: every figure and every manuscript table can be regenerated from the released CSV tables in `tables/` and the fragments in `tables/gen/`.

#### 7 Code and data access

Code and artifacts are available from the anonymized repository at `https://anonymous.4open.science/r/RepGene-E0A8/`. The released embedding data are available at `https://drive.google.com/drive/folders/1a86OycK98xvCq7ZNaDWZjtpyu2T-dghh?usp=drive_link`. Access to data and to upstream models remains subject to their original licences; the identifier-mapping and source-version checksums are recorded in the released snapshot shipped with the artifact. Authors should verify access permissions and remove any identity-revealing metadata before public release.

#### 8 Evidence boundaries and caveats

Three caveats apply to the tables released here and are repeated here so that the released files are not read beyond their scope.

- **Descriptive intervals.** All intervals are descriptive task-bootstrap intervals over the fixed register; they are not significance tests, and they overlap in every condition of the primary comparison.
- **Cross-environment re-execution.** The protocol extensions were re-run in a rebuilt environment (scikit-learn 1.9.0, numpy 2.x, torch 2.13.0+cpu) against the freeze environment (numpy 1.26.4, torch 2.1.0+cu121). Comparisons within a track are matched; the cross-environment comparison of the epoch-8 reference values is reported with this caveat, and the released epoch-8 reference column in `tables/gen/tab_budget.tex` is the drift check for it.
- **Magnitude-only records.** The historical multiplicative-interaction screening and the external random-forest track are retained as magnitude-only supplementary evidence, because fold-level records for those runs are not available; they are not used to promote any primary claim.

#### 9 Non-duplication statement

This supplementary file does not replace the Appendix of the Main Paper. The definitions, the full appendix tables, the statistical procedures, the figure-to-claim correspondence, the design-decision log, and the limitations are stated in the Main Paper and its Appendix; the release documented here is the artifact view of the same submission, and it introduces no result that is not already reported there.
