## Supplementary_Material for "RepGene: Auditing Condition-Specific Fusion Biases for Multi-View Gene Representations": best_by_task_full15.tex

\begin{tabular}{lccccc}
\toprule
Task & Core & Core+K & Core+X & Core+K+X & Spread \\
\midrule
\midrule
Blood expression cluster & INT & REL & PC & PC & 0.024 \\
Brain expression cluster & REL & REL & PC & PC & 0.064 \\
CCD Protein & REL & PC & PC & PC & 0.025 \\
CCD Transcript & REL & REL & PC & PC & 0.146 \\
Cell line expression cluster & REL & INT & PC & PC & 0.033 \\
N1 network & PC & PC & PC & PC & 0.075 \\
N1 targets & HO & REL & HO & PC & 0.048 \\
Pathology prognostics - Breast cancer & HO & REL & REL & REL & 0.005 \\
Pathology prognostics - Cervical cancer & REL & REL & HO & REL & 0.001 \\
Pathology prognostics - Colorectal cancer & REL & REL & INT & REL & 0.011 \\
TF vs non-TF & PC & PC & PC & PC & 0.037 \\
bivalent vs lys4-only & INT & REL & HO & HO & 0.023 \\
bivalent vs non-methylated & INT & INT & HO & REL & 0.005 \\
dosage sensitive vs insensitive TF & HO & HO & INT & INT & 0.009 \\
long vs short range TF & REL & HO & INT & INT & 0.025 \\
\bottomrule
\end{tabular}
