## Supplementary_Material for "RepGene: Auditing Condition-Specific Fusion Biases for Multi-View Gene Representations": condition_effects_binary9.tex

\begin{tabular}{lrrrr}
\toprule
Method & $\Delta_K$ & $\Delta_X$ & $\Delta_{KX}$ & $N_{KX}$ \\
\midrule
PCA then concat & $+0.0539$ & $+0.0672$ & $+0.0908$ & $-0.0303$ \\
Linear concat & $+0.0135$ & $+0.0296$ & $+0.0365$ & $-0.0066$ \\
PCA add & $-0.0186$ & $+0.0024$ & $-0.0107$ & $+0.0055$ \\
Interaction MLP & $+0.0120$ & $+0.0060$ & $+0.0236$ & $+0.0055$ \\
Low-rank higher-order & $+0.0243$ & $+0.0181$ & $+0.0335$ & $-0.0089$ \\
Reliability experts & $+0.0127$ & $+0.0082$ & $+0.0250$ & $+0.0041$ \\
\bottomrule
\end{tabular}
