## Supplementary_Material for "RepGene: Auditing Condition-Specific Fusion Biases for Multi-View Gene Representations": family_table.tex

% \begin{tabular}{lcccccccc}
% \toprule
% Family & \multicolumn{2}{c}{Core} & \multicolumn{2}{c}{Core+K} & \multicolumn{2}{c}{Core+X} & \multicolumn{2}{c}{Core+K+X} \\
%  & top operator & Macro-F1 & top operator & Macro-F1 & top operator & Macro-F1 & top operator & Macro-F1 \\
% \midrule
% complex/network (4 binary) & REL & 0.5906 & PC & 0.6145 & PC & 0.6561 & PC & 0.6811 \\
% TF/chromatin (5 binary) & REL & 0.7777 & HO & 0.7920 & INT & 0.7836 & REL & 0.7951 \\
% expression cluster (3 multiclass) & REL & 0.0574 & REL & 0.0619 & PC & 0.0982 & PC & 0.0958 \\
% pathology (3 multiclass) & REL & 0.3403 & REL & 0.3412 & INT & 0.3433 & REL & 0.3455 \\
% \bottomrule
% \end{tabular}

% -------------------------------------------
\setlength{\tabcolsep}{3.2pt}
\begin{tabular}{lcccccccc}
\toprule
Family 
& \multicolumn{2}{c}{Core} 
& \multicolumn{2}{c}{Core+K} 
& \multicolumn{2}{c}{Core+X} 
& \multicolumn{2}{c}{Core+K+X} \\
& op & F1 & op & F1 & op & F1 & op & F1 \\
\midrule
complex/network (4 binary) & REL & 0.5906 & PC & 0.6145 & PC & 0.6561 & PC & 0.6811 \\
TF/chromatin (5 binary) & REL & 0.7777 & HO & 0.7920 & INT & 0.7836 & REL & 0.7951 \\
expression cluster (3 multiclass) & REL & 0.0574 & REL & 0.0619 & PC & 0.0982 & PC & 0.0958 \\
pathology (3 multiclass) & REL & 0.3403 & REL & 0.3412 & INT & 0.3433 & REL & 0.3455 \\
\bottomrule
\end{tabular}


% -------------------------------------------
