## Supplementary_Material for "RepGene: Auditing Condition-Specific Fusion Biases for Multi-View Gene Representations": leakage_control_table.tex

\begin{tabular}{lrrrr}
\toprule
Method & $\Delta_{KX}$ (full) & $\Delta_{KX}$ (no overlap) & $N_{KX}$ (full) & $N_{KX}$ (no overlap) \\
\midrule
PCA then concat & $+0.0629$ & $+0.0600$ & $-0.0195$ & $-0.0214$ \\
Linear concat & $+0.0228$ & $+0.0168$ & $-0.0049$ & $-0.0047$ \\
PCA add & $-0.0067$ & $+0.0041$ & $+0.0034$ & $+0.0019$ \\
Interaction MLP & $+0.0169$ & $+0.0108$ & $+0.0022$ & $+0.0016$ \\
Low-rank higher-order & $+0.0227$ & $+0.0182$ & $-0.0051$ & $-0.0030$ \\
Reliability experts & $+0.0179$ & $+0.0169$ & $+0.0030$ & $-0.0012$ \\
\bottomrule
\end{tabular}
