## Supplementary_Material for "RepGene: Auditing Condition-Specific Fusion Biases for Multi-View Gene Representations": pertask_core_full15.tex

\begin{tabular}{lrrrrrr}
\toprule
Task & \rotatebox{90}{Linear concat} & \rotatebox{90}{PCA add} & \rotatebox{90}{PCA then concat} & \rotatebox{90}{Interaction MLP} & \rotatebox{90}{Low-rank higher-order} & \rotatebox{90}{Reliability experts} \\
\midrule
CCD Protein & 0.5204 & 0.5045 & 0.5267 & 0.5319 & 0.5223 & \textbf{0.5364} \\
CCD Transcript & 0.4145 & 0.4076 & 0.4421 & 0.5352 & 0.5335 & \textbf{0.5428} \\
N1 network & 0.6439 & 0.5939 & \textbf{0.6916} & 0.6119 & 0.5994 & 0.6178 \\
N1 targets & 0.4727 & 0.4430 & 0.5919 & 0.6697 & \textbf{0.6727} & 0.6655 \\
TF vs non-TF & 0.7608 & 0.6417 & \textbf{0.8307} & 0.7753 & 0.7732 & 0.7934 \\
bivalent vs non-methylated & 0.5126 & 0.4224 & 0.4417 & \textbf{0.8185} & 0.7974 & 0.8079 \\
bivalent vs lys4-only & 0.8084 & 0.6556 & 0.8482 & \textbf{0.8694} & 0.8552 & 0.8574 \\
dosage sensitive vs insensitive TF & 0.8545 & 0.5431 & 0.9034 & 0.9285 & \textbf{0.9295} & 0.9226 \\
long vs short range TF & 0.4281 & 0.4355 & 0.4207 & 0.4913 & 0.4737 & \textbf{0.5071} \\
\midrule
Blood expression cluster & 0.0332 & 0.0276 & 0.0395 & \textbf{0.0404} & 0.0390 & 0.0401 \\
Brain expression cluster & 0.0507 & 0.0446 & 0.0565 & 0.0615 & 0.0598 & \textbf{0.0634} \\
Cell line expression cluster & 0.0556 & 0.0474 & 0.0659 & 0.0655 & 0.0666 & \textbf{0.0686} \\
Pathology prognostics - Breast cancer & 0.3276 & 0.3276 & 0.3276 & 0.3281 & \textbf{0.3316} & 0.3310 \\
Pathology prognostics - Cervical cancer & 0.3260 & 0.3260 & 0.3260 & 0.3260 & 0.3260 & \textbf{0.3264} \\
Pathology prognostics - Colorectal cancer & 0.3274 & 0.3274 & 0.3274 & 0.3619 & 0.3618 & \textbf{0.3633} \\
\bottomrule
\end{tabular}
