## Supplementary_Material for "RepGene: Auditing Condition-Specific Fusion Biases for Multi-View Gene Representations": pertask_corek_full15.tex

\begin{tabular}{lrrrrrr}
\toprule
Task & \rotatebox{90}{Linear concat} & \rotatebox{90}{PCA add} & \rotatebox{90}{PCA then concat} & \rotatebox{90}{Interaction MLP} & \rotatebox{90}{Low-rank higher-order} & \rotatebox{90}{Reliability experts} \\
\midrule
CCD Protein & 0.5475 & 0.5022 & \textbf{0.5540} & 0.5478 & 0.5348 & 0.5483 \\
CCD Transcript & 0.4467 & 0.4078 & 0.4929 & 0.5505 & 0.5389 & \textbf{0.5510} \\
N1 network & 0.6703 & 0.5947 & \textbf{0.7136} & 0.6368 & 0.6493 & 0.6518 \\
N1 targets & 0.5242 & 0.4608 & 0.6974 & 0.6915 & 0.6926 & \textbf{0.6995} \\
TF vs non-TF & 0.7951 & 0.6307 & \textbf{0.8649} & 0.8212 & 0.8197 & 0.8232 \\
bivalent vs non-methylated & 0.5108 & 0.4240 & 0.6445 & \textbf{0.8155} & 0.8129 & 0.8111 \\
bivalent vs lys4-only & 0.7879 & 0.5773 & 0.8733 & 0.8695 & 0.8797 & \textbf{0.8798} \\
dosage sensitive vs insensitive TF & 0.8338 & 0.4401 & 0.9136 & 0.9305 & \textbf{0.9325} & 0.9286 \\
long vs short range TF & 0.4207 & 0.4426 & 0.4281 & 0.4768 & \textbf{0.5152} & 0.4716 \\
\midrule
Blood expression cluster & 0.0352 & 0.0267 & 0.0407 & 0.0447 & 0.0439 & \textbf{0.0455} \\
Brain expression cluster & 0.0550 & 0.0417 & 0.0624 & 0.0665 & 0.0659 & \textbf{0.0671} \\
Cell line expression cluster & 0.0602 & 0.0469 & 0.0714 & \textbf{0.0735} & 0.0716 & 0.0731 \\
Pathology prognostics - Breast cancer & 0.3276 & 0.3276 & 0.3276 & 0.3287 & 0.3287 & \textbf{0.3297} \\
Pathology prognostics - Cervical cancer & 0.3260 & 0.3260 & 0.3260 & 0.3260 & 0.3260 & \textbf{0.3268} \\
Pathology prognostics - Colorectal cancer & 0.3274 & 0.3274 & 0.3274 & 0.3645 & 0.3528 & \textbf{0.3669} \\
\bottomrule
\end{tabular}
