## Supplementary_Material for "RepGene: Auditing Condition-Specific Fusion Biases for Multi-View Gene Representations": pertask_corekx_full15.tex

\begin{tabular}{lrrrrrr}
\toprule
Task & \rotatebox{90}{Linear concat} & \rotatebox{90}{PCA add} & \rotatebox{90}{PCA then concat} & \rotatebox{90}{Interaction MLP} & \rotatebox{90}{Low-rank higher-order} & \rotatebox{90}{Reliability experts} \\
\midrule
CCD Protein & 0.5564 & 0.4941 & \textbf{0.5612} & 0.5427 & 0.5423 & 0.5521 \\
CCD Transcript & 0.5307 & 0.4105 & \textbf{0.6886} & 0.5750 & 0.5813 & 0.5749 \\
N1 network & 0.6928 & 0.5743 & \textbf{0.7663} & 0.6722 & 0.6732 & 0.6891 \\
N1 targets & 0.5361 & 0.4669 & \textbf{0.7084} & 0.6790 & 0.6868 & 0.6842 \\
TF vs non-TF & 0.7875 & 0.6286 & \textbf{0.8675} & 0.8189 & 0.8223 & 0.8325 \\
bivalent vs non-methylated & 0.5607 & 0.4667 & 0.6923 & 0.8118 & 0.8151 & \textbf{0.8187} \\
bivalent vs lys4-only & 0.8009 & 0.6513 & 0.8810 & 0.8736 & \textbf{0.8919} & 0.8817 \\
dosage sensitive vs insensitive TF & 0.8433 & 0.4380 & 0.9209 & \textbf{0.9389} & 0.9302 & 0.9337 \\
long vs short range TF & 0.4355 & 0.4207 & 0.4281 & \textbf{0.5319} & 0.5150 & 0.5088 \\
\midrule
Blood expression cluster & 0.0348 & 0.0285 & \textbf{0.0604} & 0.0484 & 0.0499 & 0.0486 \\
Brain expression cluster & 0.0573 & 0.0428 & \textbf{0.1249} & 0.0755 & 0.0755 & 0.0745 \\
Cell line expression cluster & 0.0619 & 0.0449 & \textbf{0.1020} & 0.0785 & 0.0775 & 0.0771 \\
Pathology prognostics - Breast cancer & 0.3276 & 0.3276 & 0.3276 & 0.3325 & 0.3292 & \textbf{0.3352} \\
Pathology prognostics - Cervical cancer & 0.3260 & 0.3260 & 0.3260 & 0.3264 & 0.3260 & \textbf{0.3273} \\
Pathology prognostics - Colorectal cancer & 0.3274 & 0.3274 & 0.3274 & 0.3633 & 0.3666 & \textbf{0.3742} \\
\bottomrule
\end{tabular}
