## Supplementary_Material for "RepGene: Auditing Condition-Specific Fusion Biases for Multi-View Gene Representations": pertask_corex_full15.tex

\begin{tabular}{lrrrrrr}
\toprule
Task & \rotatebox{90}{Linear concat} & \rotatebox{90}{PCA add} & \rotatebox{90}{PCA then concat} & \rotatebox{90}{Interaction MLP} & \rotatebox{90}{Low-rank higher-order} & \rotatebox{90}{Reliability experts} \\
\midrule
CCD Protein & 0.5396 & 0.5070 & \textbf{0.5470} & 0.5272 & 0.5395 & 0.5361 \\
CCD Transcript & 0.5113 & 0.4150 & \textbf{0.6824} & 0.5634 & 0.5690 & 0.5572 \\
N1 network & 0.6834 & 0.5934 & \textbf{0.7670} & 0.6299 & 0.6413 & 0.6625 \\
N1 targets & 0.5086 & 0.4328 & 0.6281 & 0.6474 & \textbf{0.6599} & 0.6587 \\
TF vs non-TF & 0.7714 & 0.6309 & \textbf{0.8314} & 0.7749 & 0.7844 & 0.7959 \\
bivalent vs non-methylated & 0.5544 & 0.4915 & 0.6290 & 0.8143 & \textbf{0.8203} & 0.8134 \\
bivalent vs lys4-only & 0.8208 & 0.6822 & 0.8630 & 0.8675 & \textbf{0.8692} & 0.8654 \\
dosage sensitive vs insensitive TF & 0.8576 & 0.4963 & 0.9195 & \textbf{0.9366} & 0.9318 & 0.9294 \\
long vs short range TF & 0.4347 & 0.4200 & 0.4347 & \textbf{0.5246} & 0.5044 & 0.5063 \\
\midrule
Blood expression cluster & 0.0381 & 0.0265 & \textbf{0.0648} & 0.0456 & 0.0451 & 0.0443 \\
Brain expression cluster & 0.0604 & 0.0452 & \textbf{0.1277} & 0.0723 & 0.0726 & 0.0702 \\
Cell line expression cluster & 0.0588 & 0.0475 & \textbf{0.1020} & 0.0733 & 0.0721 & 0.0724 \\
Pathology prognostics - Breast cancer & 0.3276 & 0.3276 & 0.3276 & 0.3310 & 0.3287 & \textbf{0.3331} \\
Pathology prognostics - Cervical cancer & 0.3260 & 0.3260 & 0.3260 & 0.3263 & \textbf{0.3269} & 0.3260 \\
Pathology prognostics - Colorectal cancer & 0.3274 & 0.3274 & 0.3274 & \textbf{0.3726} & 0.3718 & 0.3665 \\
\bottomrule
\end{tabular}
