## Supplementary_Material for "RepGene: Auditing Condition-Specific Fusion Biases for Multi-View Gene Representations": secondary_full15.tex

\begin{tabular}{l@{\hspace{6pt}}rrrrrrrr}
\toprule
Method & \multicolumn{2}{c}{Core} & \multicolumn{2}{c}{Core+K} & \multicolumn{2}{c}{Core+X} & \multicolumn{2}{c}{Core+K+X} \\
 & BAcc & AUC & BAcc & AUC & BAcc & AUC & BAcc & AUC \\
\midrule
Reliability experts & 0.5046 & 0.7170 & 0.5138 & 0.7310 & 0.5116 & 0.7311 & 0.5224 & 0.7441 \\
Interaction MLP & 0.5029 & 0.7157 & 0.5115 & 0.7328 & 0.5080 & 0.7274 & 0.5184 & 0.7462 \\
Low-rank higher-order & 0.4988 & 0.7116 & 0.5127 & 0.7350 & 0.5104 & 0.7325 & 0.5196 & 0.7467 \\
PCA then concat & 0.4766 & 0.7299 & 0.5013 & 0.7492 & 0.5150 & 0.7823 & 0.5269 & 0.7894 \\
Linear concat & 0.4563 & 0.7035 & 0.4619 & 0.7186 & 0.4691 & 0.7234 & 0.4707 & 0.7365 \\
PCA add & 0.4133 & 0.6671 & 0.4061 & 0.6631 & 0.4141 & 0.6666 & 0.4092 & 0.6648 \\
\bottomrule
\end{tabular}
