## Supplementary_Material for "RepGene: Auditing Condition-Specific Fusion Biases for Multi-View Gene Representations": tab_budget.tex

\begin{table}[h]
\centering\footnotesize
\caption{Nine binary tasks under Core+K+X, task-equal Macro-F1 against the label-free training budget. Columns 2--3 are the matched re-execution; the last column is the frozen epoch-8 reference for drift comparison. PCA then concat and Linear concat were not re-run at higher budgets (the budget question concerns the learned operators) and are shown for the epoch-8 reference only.}
\label{tab:budget}
\begin{tabular}{lrrrr}
\toprule
Method & 8 epochs & 32 epochs & 64 epochs & 8 epochs (frozen ref.) \\
\midrule
Reliability experts & 0.6995 & 0.7217 & 0.7354 & 0.6993 \\
Interaction MLP & 0.6965 & 0.7208 & 0.7308 & 0.6963 \\
Low-rank higher-order & 0.6957 & 0.7182 & 0.7279 & 0.6958 \\
Cross-attention & -- & 0.7198 & 0.7318 & 0.6936 \\
PCA then concat & -- & -- & -- & -- \\
Linear concat & -- & -- & -- & -- \\
\bottomrule
\end{tabular}
\end{table}
