## Supplementary_Material for "RepGene: Auditing Condition-Specific Fusion Biases for Multi-View Gene Representations": tab_calibration.tex

\begin{table}[h]
\centering\footnotesize
\caption{Calibration and threshold sensitivity on the nine binary tasks, task-equal means. Macro-F1 is reported at the frozen $0.5$ threshold and at the best swept threshold of the same probabilities; Brier and ECE quantify calibration; ROC-AUC quantifies rank discrimination.}
\label{tab:calibration}
\begin{tabular}{lrrrrr}
\toprule
Method & F1@$0.5$ & F1@best & Brier & ECE & AUC \\
\midrule
\multicolumn{6}{l}{\textit{RF probe}} \\
\midrule
Reliability experts & 0.7045 & 0.7372 & 0.3196 & 0.0749 & 0.9378 \\
Interaction MLP & 0.7040 & 0.7347 & 0.3212 & 0.0730 & 0.9338 \\
Low-rank higher-order & 0.7037 & 0.7367 & 0.3215 & 0.0761 & 0.9320 \\
PCA then concat & 0.6866 & 0.7708 & 0.3291 & 0.1270 & 0.9387 \\
Linear concat & 0.6220 & 0.7342 & 0.3662 & 0.1182 & 0.9026 \\
PCA add & 0.5100 & 0.6712 & 0.4063 & 0.0988 & 0.7988 \\
\midrule
\multicolumn{6}{l}{\textit{linear probe}} \\
\midrule
Reliability experts & 0.7146 & 0.7394 & 0.3515 & 0.1198 & 0.9383 \\
Interaction MLP & 0.7104 & 0.7363 & 0.3552 & 0.1212 & 0.9296 \\
Low-rank higher-order & 0.7084 & 0.7331 & 0.3665 & 0.1350 & 0.9318 \\
PCA then concat & 0.7291 & 0.7563 & 0.3605 & 0.1518 & 0.9303 \\
Linear concat & 0.7318 & 0.7603 & 0.3389 & 0.1322 & 0.9211 \\
PCA add & 0.6399 & 0.6693 & 0.4663 & 0.1859 & 0.7873 \\
\bottomrule
\end{tabular}
\end{table}
