## Supplementary_Material for "RepGene: Auditing Condition-Specific Fusion Biases for Multi-View Gene Representations": tab_cell_boundary.tex

\begin{table}[t]
\centering\footnotesize
\setlength{\tabcolsep}{6pt}
\renewcommand{\arraystretch}{0.88}
\caption{Gene-to-cell boundary diagnostic on the aorta annotation task: Macro-F1 over five seeds ($\pm$ seed SD). The expression-only reference leads and a dimension-matched random 256-D control sits within $0.004$ of the best fused representation; one fixed design, not a validated method.}
\label{tab:cell-boundary}
\begin{tabular}{lr}
\toprule
Representation & Macro-F1 \\
\midrule
Expression only & $\mathbf{0.9295}$ $\pm$ 0.0047 \\
PCA on expression & 0.9123 $\pm$ 0.0101 \\
Linear concat & 0.9115 $\pm$ 0.0136 \\
PCA then concat & 0.9091 $\pm$ 0.0104 \\
PCA add & 0.9078 $\pm$ 0.0138 \\
Low-rank higher-order & 0.9024 $\pm$ 0.0129 \\
Random 256-D control & 0.8987 $\pm$ 0.0100 \\
Interaction MLP & 0.8985 $\pm$ 0.0241 \\
Reliability experts & 0.8889 $\pm$ 0.0246 \\
\bottomrule
\end{tabular}
\end{table}
