## Supplementary_Material for "RepGene: Auditing Condition-Specific Fusion Biases for Multi-View Gene Representations": tab_condition_effects.tex

\begin{table}[t]
\centering\footnotesize
\setlength{\tabcolsep}{6pt}
\renewcommand{\arraystretch}{0.88}
\caption{Matched condition deltas relative to Core and the non-additivity term $N_{KX}$ over the fifteen-task register; boldface marks the largest gain per contrast. Descriptive decompositions, not causal effects or significance tests.}
\label{tab:condition-effects}
\begin{tabular}{lrrrr}
\toprule
Method & $\Delta_K$ & $\Delta_X$ & $\Delta_{KX}$ & $N_{KX}$ \\
\midrule
Linear concat & $+0.0088$ & $+0.0189$ & $+0.0228$ & $-0.0049$ \\
PCA add & $-0.0114$ & $+0.0014$ & $-0.0067$ & $+0.0034$ \\
PCA then concat & $\mathbf{+0.0332}$ & $\mathbf{+0.0492}$ & $\mathbf{+0.0629}$ & $\mathbf{-0.0195}$ \\
Interaction MLP & $+0.0086$ & $+0.0061$ & $+0.0169$ & $+0.0022$ \\
Low-rank higher-order & $+0.0149$ & $+0.0130$ & $+0.0227$ & $-0.0051$ \\
Reliability experts & $+0.0087$ & $+0.0062$ & $+0.0179$ & $+0.0030$ \\
\bottomrule
\end{tabular}
\end{table}
