## Supplementary_Material for "RepGene: Auditing Condition-Specific Fusion Biases for Multi-View Gene Representations": tab_condition_means.tex

\begin{table}[t]
\centering\footnotesize
\setlength{\tabcolsep}{5pt}
\renewcommand{\arraystretch}{0.88}
\caption{Task-equal Macro-F1 by protocol condition over the fifteen-task register (six operators, three seeds, three folds); boldface marks the per-condition leader. All intervals overlap (widths $\approx\pm0.13$).}
\label{tab:condition-means}
\begin{tabular}{lrrrr}
\toprule
Method & Core & Core+K & Core+X & Core+K+X \\
\midrule
Linear concat & $0.4358$ & $0.4446$ & $0.4547$ & $0.4586$ \\
PCA add & $0.3832$ & $0.3718$ & $0.3846$ & $0.3765$ \\
PCA then concat & $0.4560$ & $0.4892$ & $\mathbf{0.5052}$ & $\mathbf{0.5188}$ \\
Interaction MLP & $0.4943$ & $0.5029$ & $0.5005$ & $0.5112$ \\
Low-rank higher-order & $0.4895$ & $0.5043$ & $0.5025$ & $0.5122$ \\
Reliability experts & $\mathbf{0.4962}$ & $\mathbf{0.5049}$ & $0.5025$ & $0.5142$ \\
\bottomrule
\end{tabular}
\end{table}
