## Supplementary_Material for "RepGene: Auditing Condition-Specific Fusion Biases for Multi-View Gene Representations": tab_extensions.tex

\begin{table}[t]
\centering\footnotesize
\setlength{\tabcolsep}{4pt}
\caption{Summary of the four registered protocol extensions, each varying one factor of the frozen contract while the rest stays fixed. The following subsections give the full tables, the provenance of the re-execution, and the per-track caveats.}
\label{tab:extensions}
\begin{tabular}{@{}p{0.15\linewidth}p{0.24\linewidth}p{0.53\linewidth}@{}}
\toprule
Extension & Changed factor & Outcome \\
\midrule
Probe family & RF $\rightarrow$ logistic probe, identical exports (15 tasks $\times$ 4 conditions) & Structure survives: no operator leads in all four conditions and PCA then concat stays least decomposable ($N_{KX}=-0.0204$ RF, $-0.0116$ linear), but the leader changes with the probe and the experts' edge over Linear concat does not survive ($0.4929$ vs.\ $0.4366$ $\rightarrow$ $0.4964$ vs.\ $0.5057$) \\
Supervised head & RF $\rightarrow$ supervised MLP head (binary9 $\times$ Core+K+X) & Linear concat leads ($0.7523$); the learned experts follow ($0.7257$--$0.7326$) \\
Training budget & $8 \rightarrow 32 \rightarrow 64$ epochs, four learned operators (binary9 $\times$ Core+K+X) & Every operator improves ($\approx+0.022$/$+0.031$), yet cross-attention never leads (residual gap $+0.0057$/$+0.0020$/$+0.0036$; at most two of nine per-task wins) \\
Pairing control & real pairing vs.\ capacity-matched shuffled twin & Real pairing wins in all four conditions: paired mean $+0.0259$, median $+0.0212$, $94\%$ of $32$ cells positive \\
Calibration & Brier/ECE and threshold sweeps from stored probabilities & PCA then concat ranks best (AUC $0.9387$) and is calibrated worst (ECE $0.1270$): it loses at the frozen $0.5$ threshold ($0.6866$ vs.\ $0.7045$) and wins once thresholds are tuned ($0.7708$ vs.\ $0.7372$) \\
\bottomrule
\end{tabular}
\end{table}
