## Supplementary_Material for "RepGene: Auditing Condition-Specific Fusion Biases for Multi-View Gene Representations": tab_external_track.tex

\begin{table}[t]
\centering\footnotesize
\setlength{\tabcolsep}{6pt}
\renewcommand{\arraystretch}{0.88}
\caption{External RF track over the nine binary tasks under Core+K+X: task-equal Macro-F1 with descriptive 95\% intervals (two decimals); the probe was configured before the internal results were inspected.}
\label{tab:external-binary}
\begin{tabular}{lr}
\toprule
Method & Macro-F1 [95\%] \\
\midrule
Random 256-D & 0.4240 [0.41, 0.45] \\
PCA add & 0.5103 [0.46, 0.57] \\
GenePT (text only) & 0.5479 [0.48, 0.64] \\
Linear concat & 0.6342 [0.54, 0.73] \\
PRISME+scRNA ext. & 0.7116 [0.62, 0.80] \\
Interaction MLP & 0.7192 [0.63, 0.81] \\
PCA then concat & 0.7221 [0.62, 0.81] \\
Reliability experts & 0.7227 [0.63, 0.82] \\
Low-rank higher-order & $\mathbf{0.7251}$ [0.63, 0.81] \\
\bottomrule
\end{tabular}
\end{table}
