## Supplementary_Material for "RepGene: Auditing Condition-Specific Fusion Biases for Multi-View Gene Representations": tab_probe_sensitivity.tex

\begin{table}[h]
\centering\footnotesize
\caption{Fifteen-task task-equal Macro-F1 under two probes evaluated on identical exported representations (three seeds, three folds, four conditions). Boldface marks the per-condition leader. Bootstrap intervals of width $\approx\pm0.13$ overlap throughout, as in the main register.}
\label{tab:probe-sensitivity}
\begin{tabular}{lrrrr}
\toprule
Method & Core & Core+K & Core+X & Core+K+X \\
\midrule
\multicolumn{5}{l}{\textit{RF probe (frozen primary)}} \\
\midrule
Linear concat & $0.4366$ & $0.4448$ & $0.4544$ & $0.4588$ \\
PCA add & $0.3838$ & $0.3711$ & $0.3853$ & $0.3765$ \\
PCA then concat & $0.4565$ & $0.4898$ & $\mathbf{0.5058}$ & $\mathbf{0.5188}$ \\
Interaction MLP & $0.4929$ & $0.5053$ & $0.5021$ & $0.5118$ \\
Low-rank higher-order & $0.4920$ & $\mathbf{0.5056}$ & $0.5020$ & $0.5109$ \\
Reliability experts & $\mathbf{0.4951}$ & $0.5034$ & $0.5019$ & $0.5135$ \\
\midrule
\multicolumn{5}{l}{\textit{linear (logistic) probe}} \\
\midrule
Linear concat & $\mathbf{0.5057}$ & $0.5190$ & $0.5326$ & $0.5355$ \\
PCA add & $0.4668$ & $0.4598$ & $0.4602$ & $0.4605$ \\
PCA then concat & $0.5019$ & $\mathbf{0.5201}$ & $\mathbf{0.5357}$ & $\mathbf{0.5423}$ \\
Interaction MLP & $0.4964$ & $0.5056$ & $0.5060$ & $0.5179$ \\
Low-rank higher-order & $0.4902$ & $0.5076$ & $0.5057$ & $0.5185$ \\
Reliability experts & $0.4978$ & $0.5060$ & $0.5094$ & $0.5199$ \\
\bottomrule
\end{tabular}
\end{table}
