## Supplementary_Material for "RepGene: Auditing Condition-Specific Fusion Biases for Multi-View Gene Representations": tab_shuffle.tex

\begin{table}[h]
\centering\footnotesize
\caption{View-pair shuffle control on the nine binary tasks (three seeds, four conditions, random-forest probe). The shuffled twin is capacity-matched; only the gene-wise cross-view pairing is destroyed.}
\label{tab:shuffle}
\begin{tabular}{lrrrr}
\toprule
Variant & Core & Core+K & Core+X & Core+K+X \\
\midrule
Real pairing & 0.6702 & 0.6888 & 0.6768 & 0.6974 \\
Pair-shuffled twin & 0.6529 & 0.6619 & 0.6538 & 0.6611 \\
\midrule
Difference & $+0.0173$ & $+0.0269$ & $+0.0230$ & $+0.0363$ \\
\bottomrule
\end{tabular}
\end{table}
