## Supplementary_Material for "RepGene: Auditing Condition-Specific Fusion Biases for Multi-View Gene Representations": tab_supervised_ref.tex

\begin{table}[h]
\centering\footnotesize
\caption{Supervised MLP-head reference (B2) on the nine binary tasks under Core+K+X, alongside the frozen random-forest probe on the same exported representations. The trained head does not restore the learned experts' advantage over the concatenation baselines.}
\label{tab:supervised-ref}
\begin{tabular}{lrr}
\toprule
Method & Supervised MLP head & RF probe \\
\midrule
Linear concat & 0.7523 & 0.6385 \\
PCA then concat & 0.7483 & 0.7236 \\
Low-rank higher-order & 0.7326 & 0.7206 \\
Interaction MLP & 0.7261 & 0.7184 \\
Reliability experts & 0.7257 & 0.7169 \\
PCA add & 0.6707 & 0.5059 \\
\bottomrule
\end{tabular}
\end{table}
