## Supplementary figures and images for "RepGene: Auditing Condition-Specific Fusion Biases for Multi-View Gene Representations"

### cka_matrix_all_methods.pdf

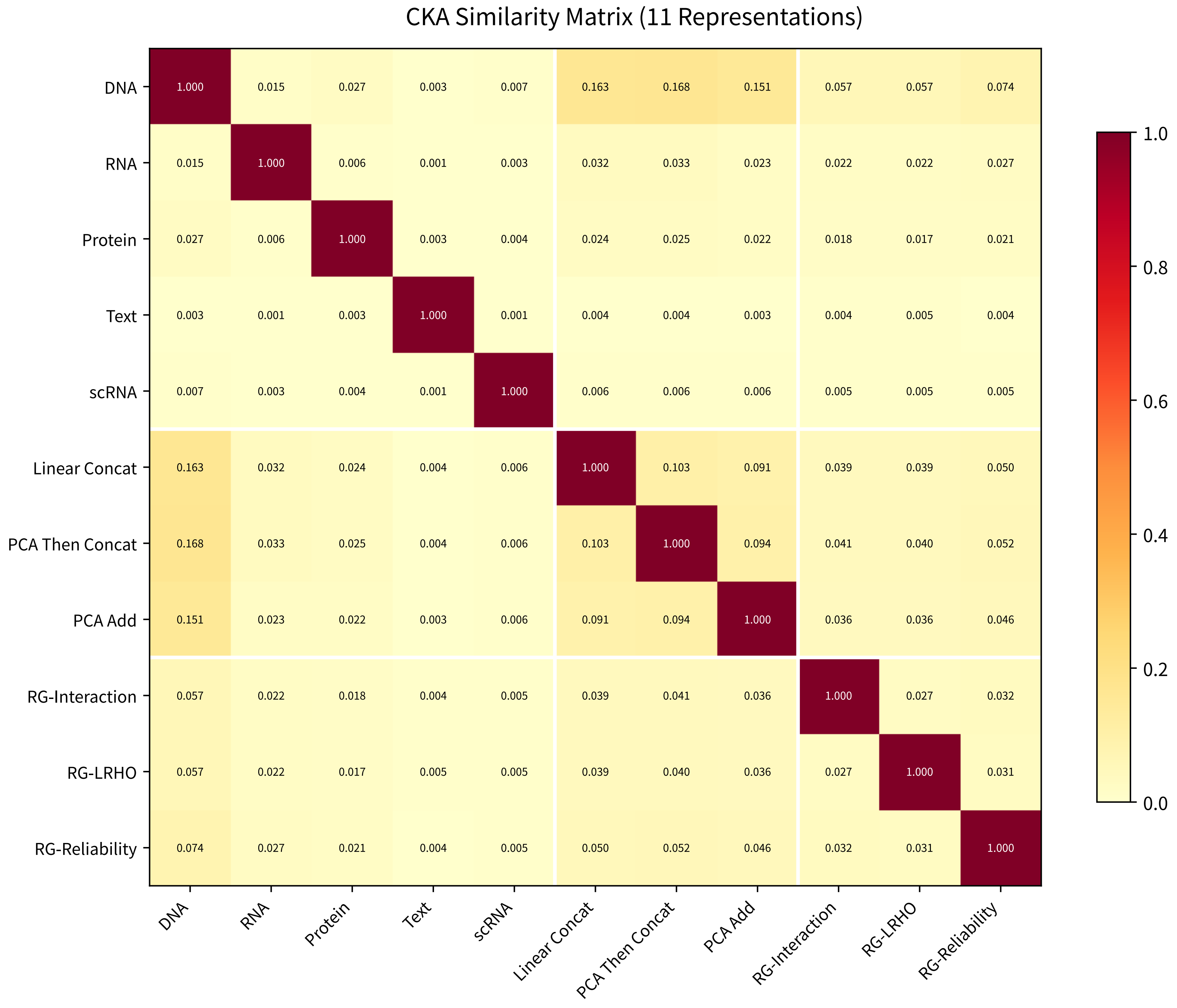

### cross_view_knn_overlap.pdf

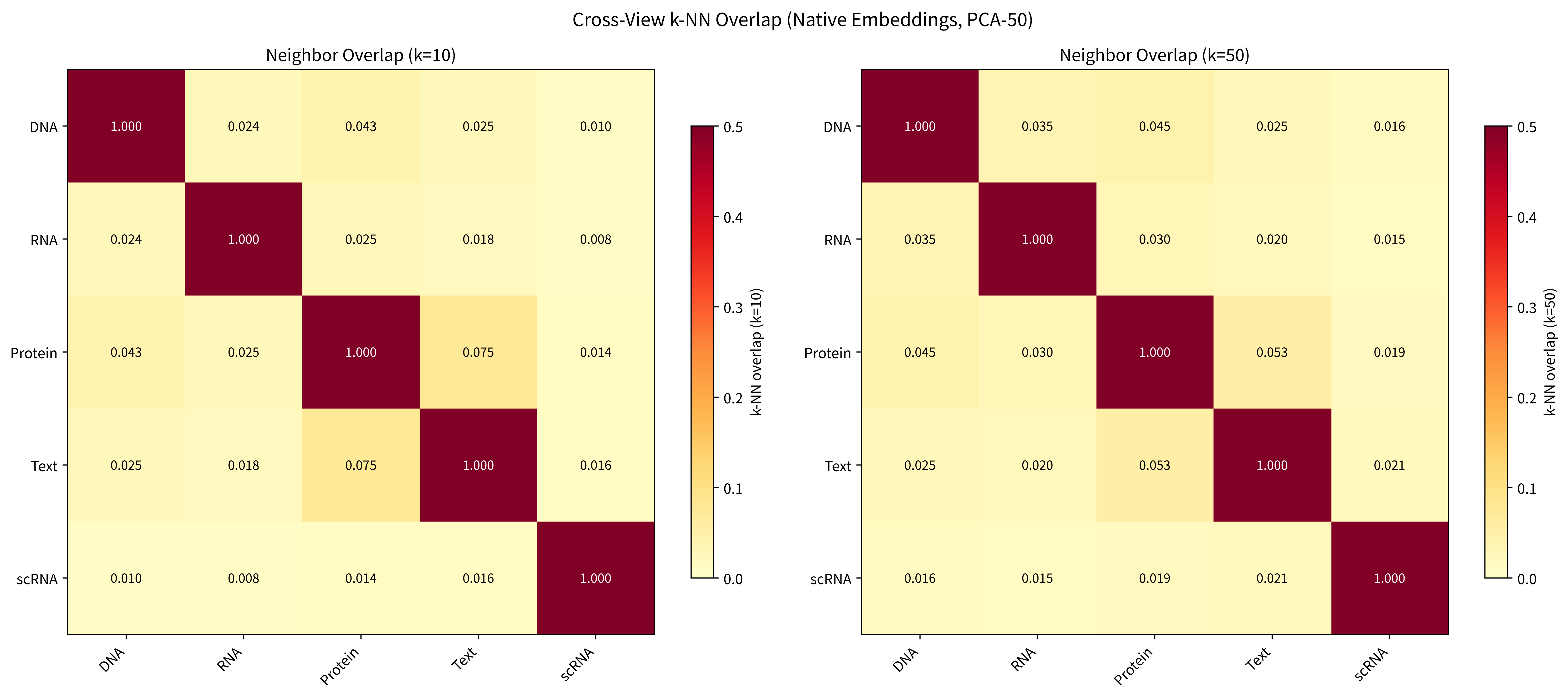

### emb_cross_view_cka_heatmap.pdf

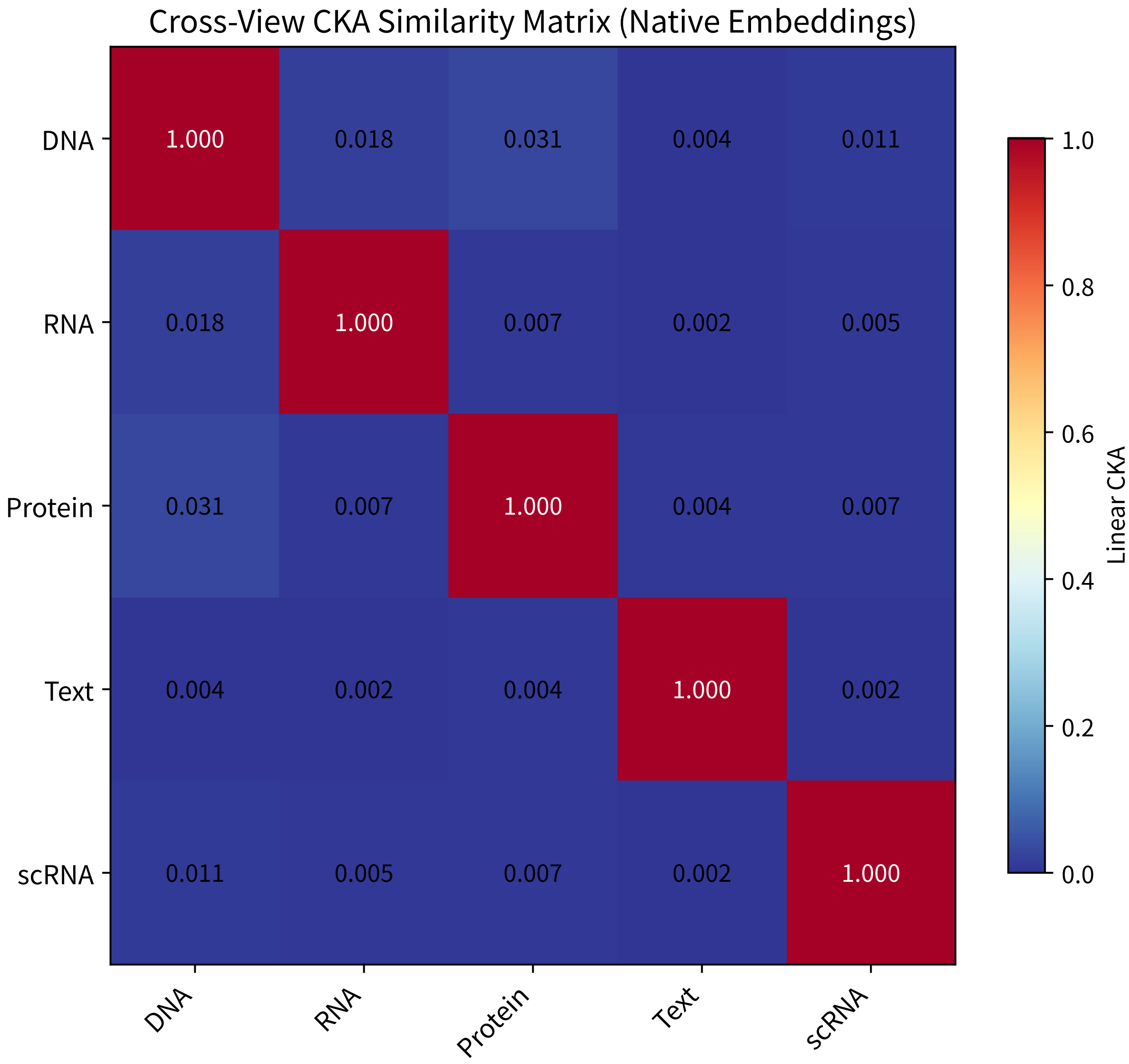

### emb_embedding_stats_all.pdf

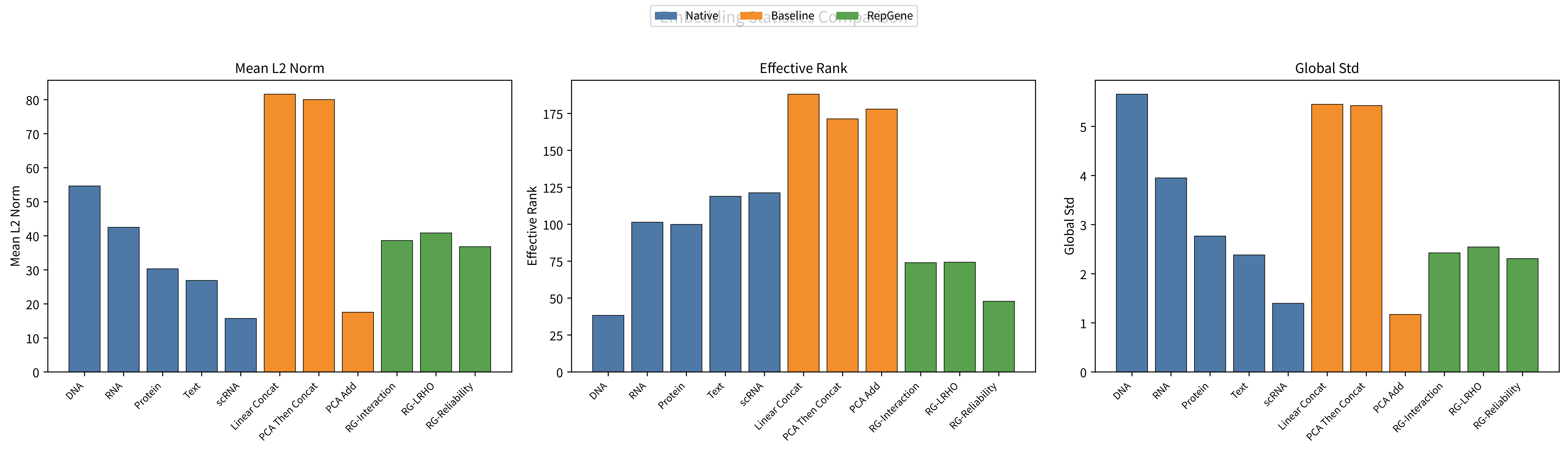

### emb_fusion_umap_comparison_01.pdf

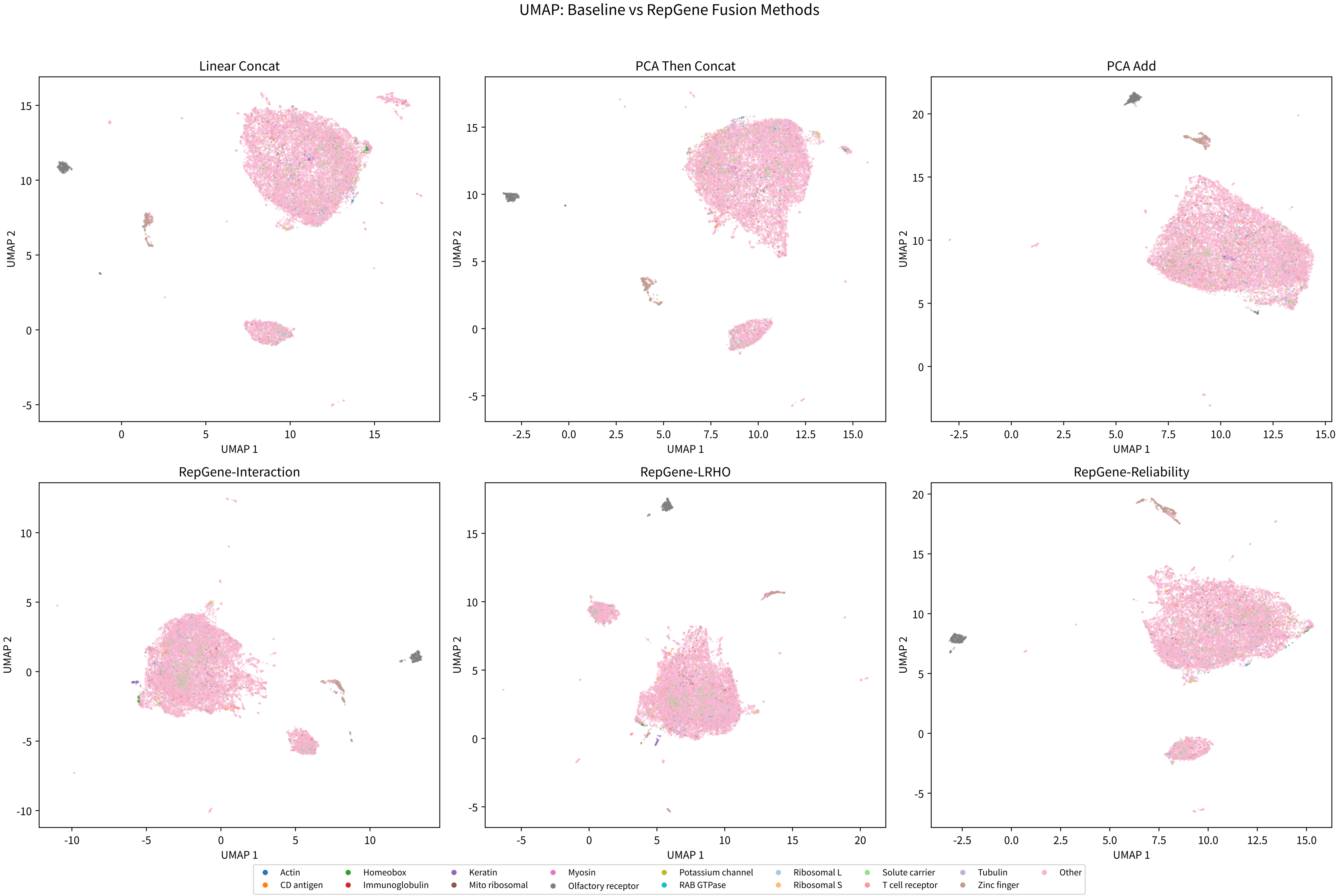

### emb_knn_overlap_k10_heatmap.pdf

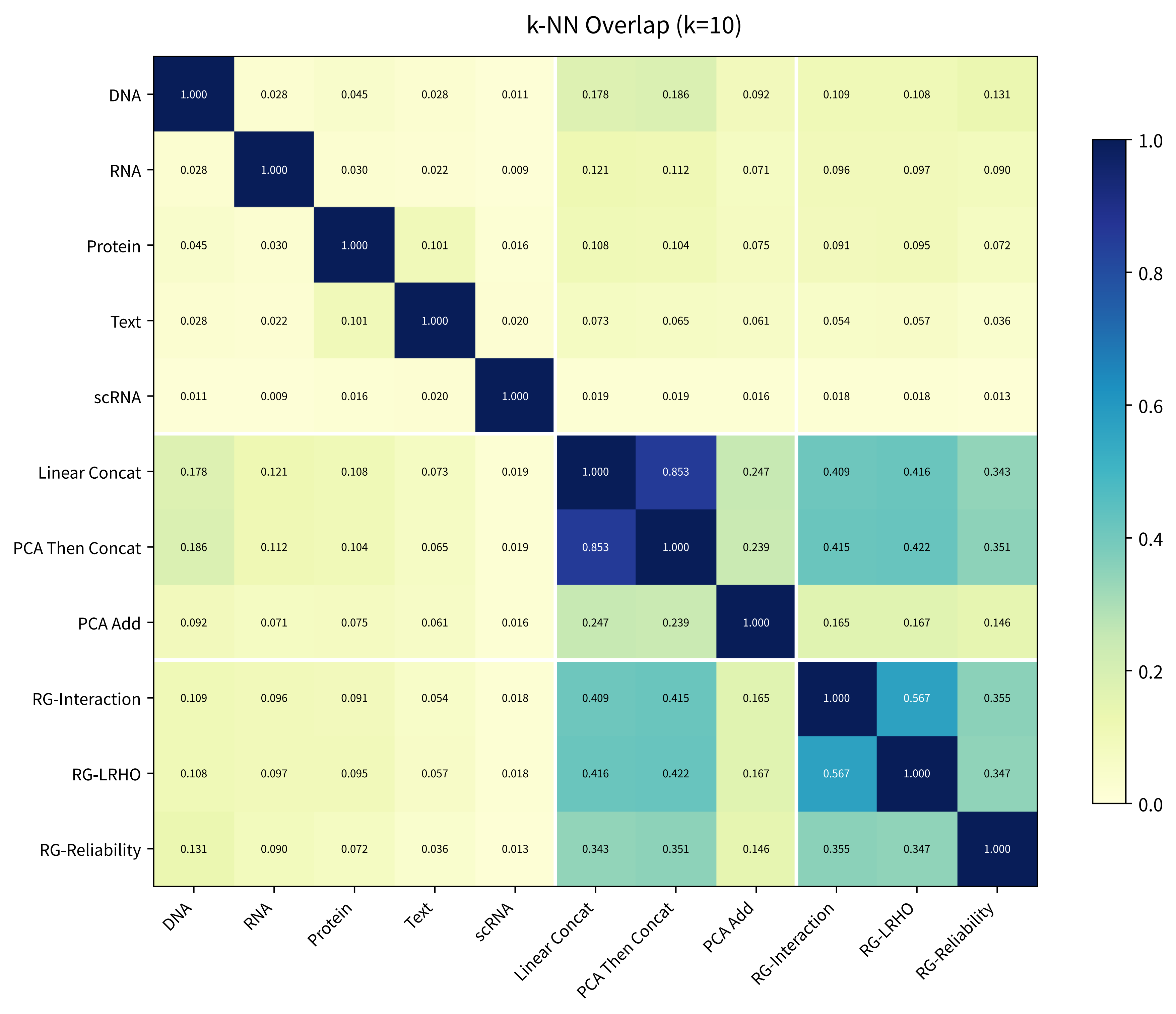

### emb_native_umap_panel_01.pdf

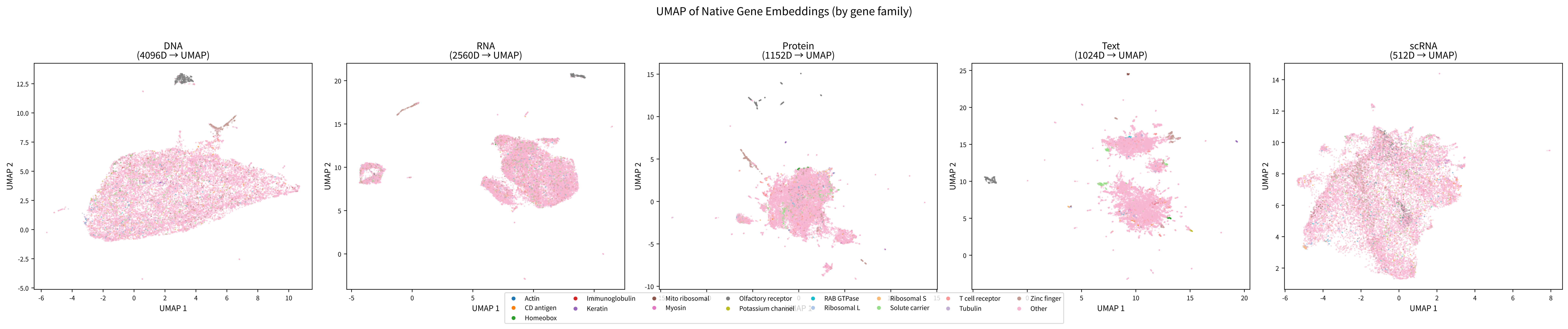

### emb_pca_variance_explained.pdf

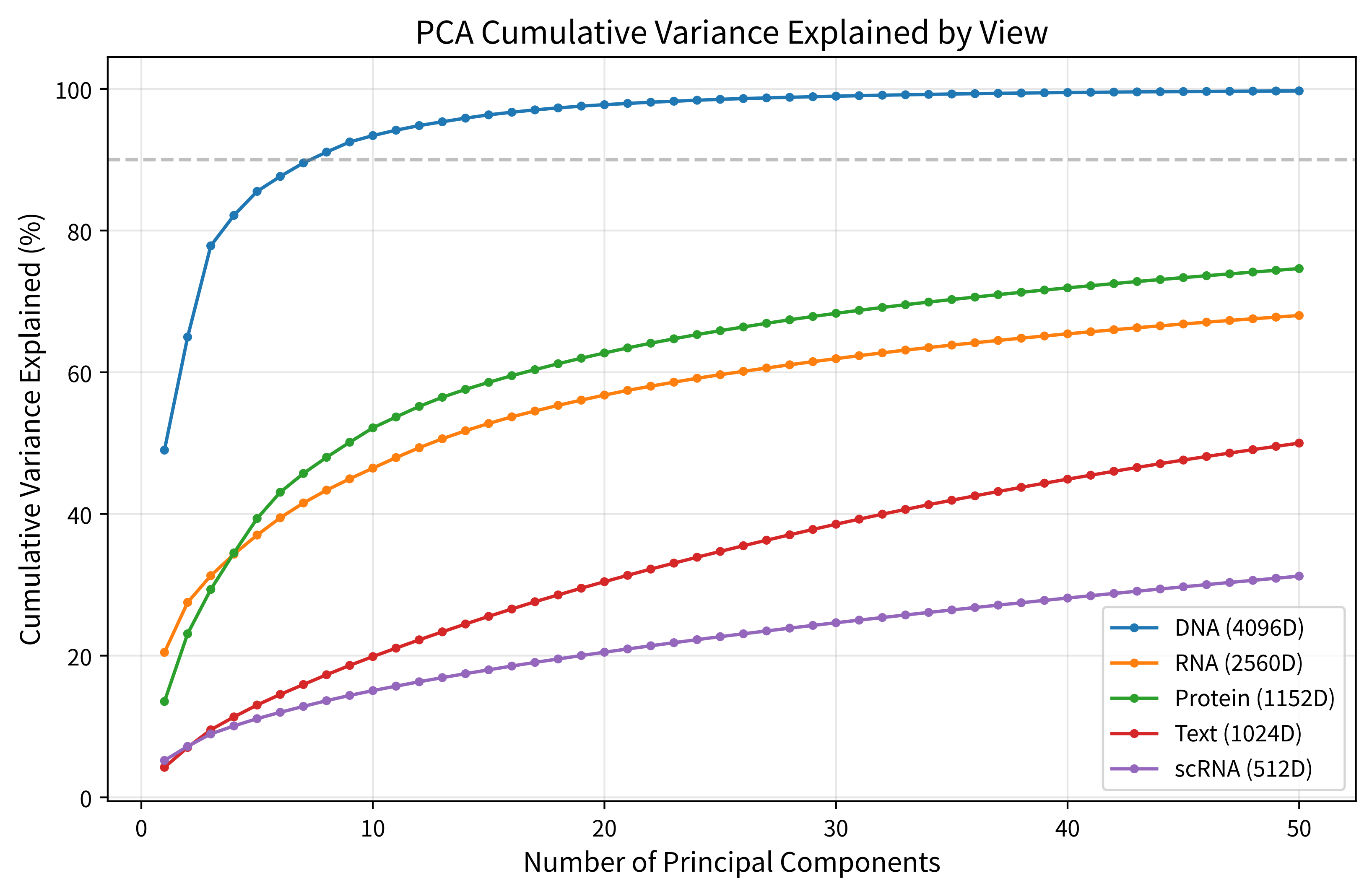

### emb_silhouette_comparison_all.pdf

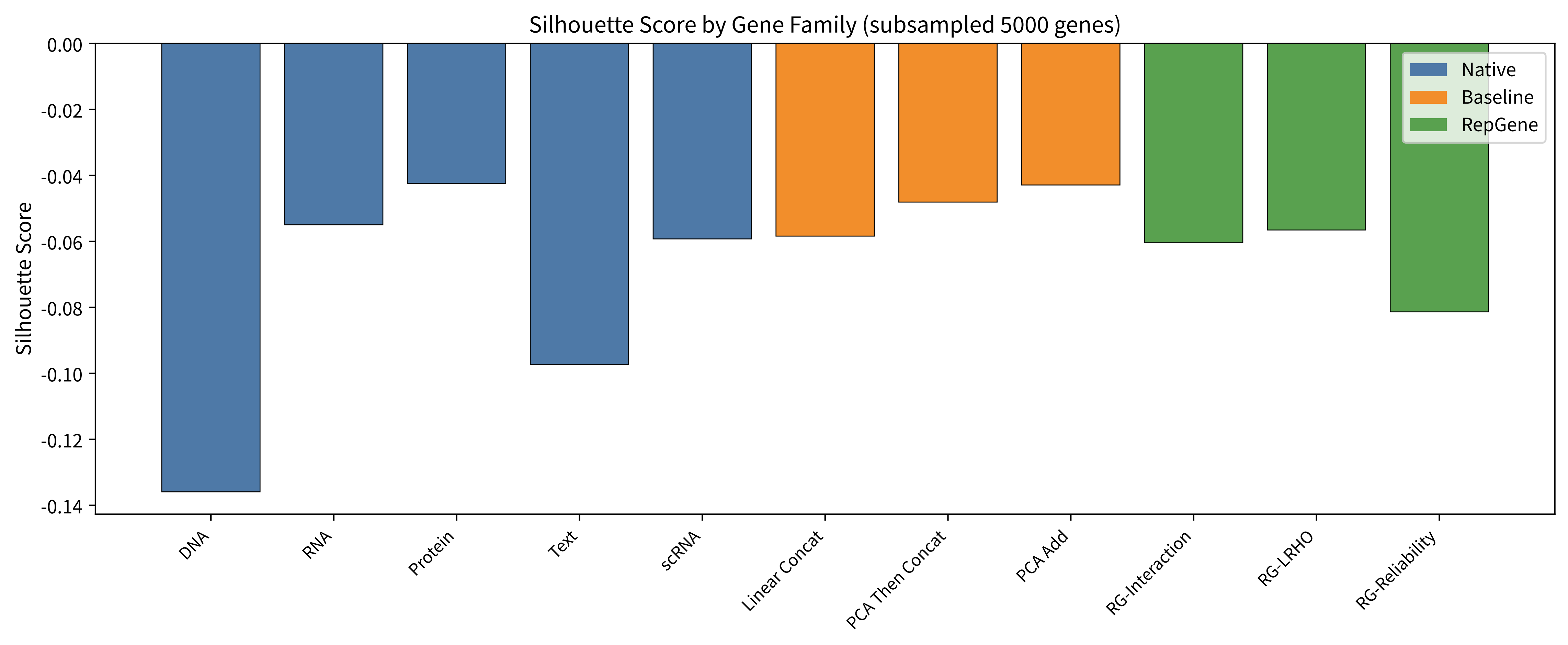

### emb_tsne_master_panel_01.pdf

t-SNE Comparison: Native Views vs Baseline vs RepGene Fusion (by gene family)

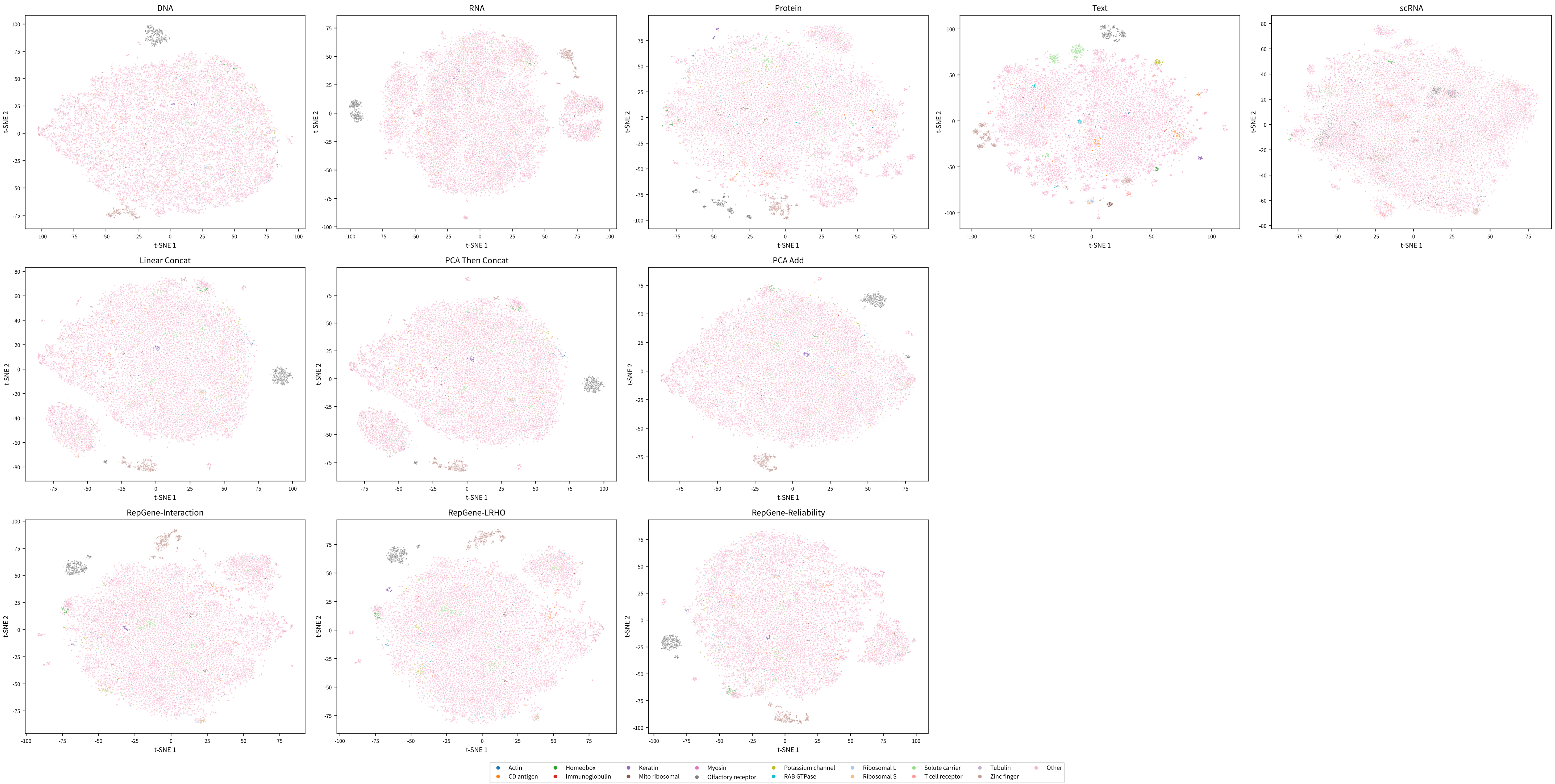

### emb_umap_master_panel_01.pdf

UMAP Comparison: Native Views vs Baseline vs RepGene Fusion (by gene family)

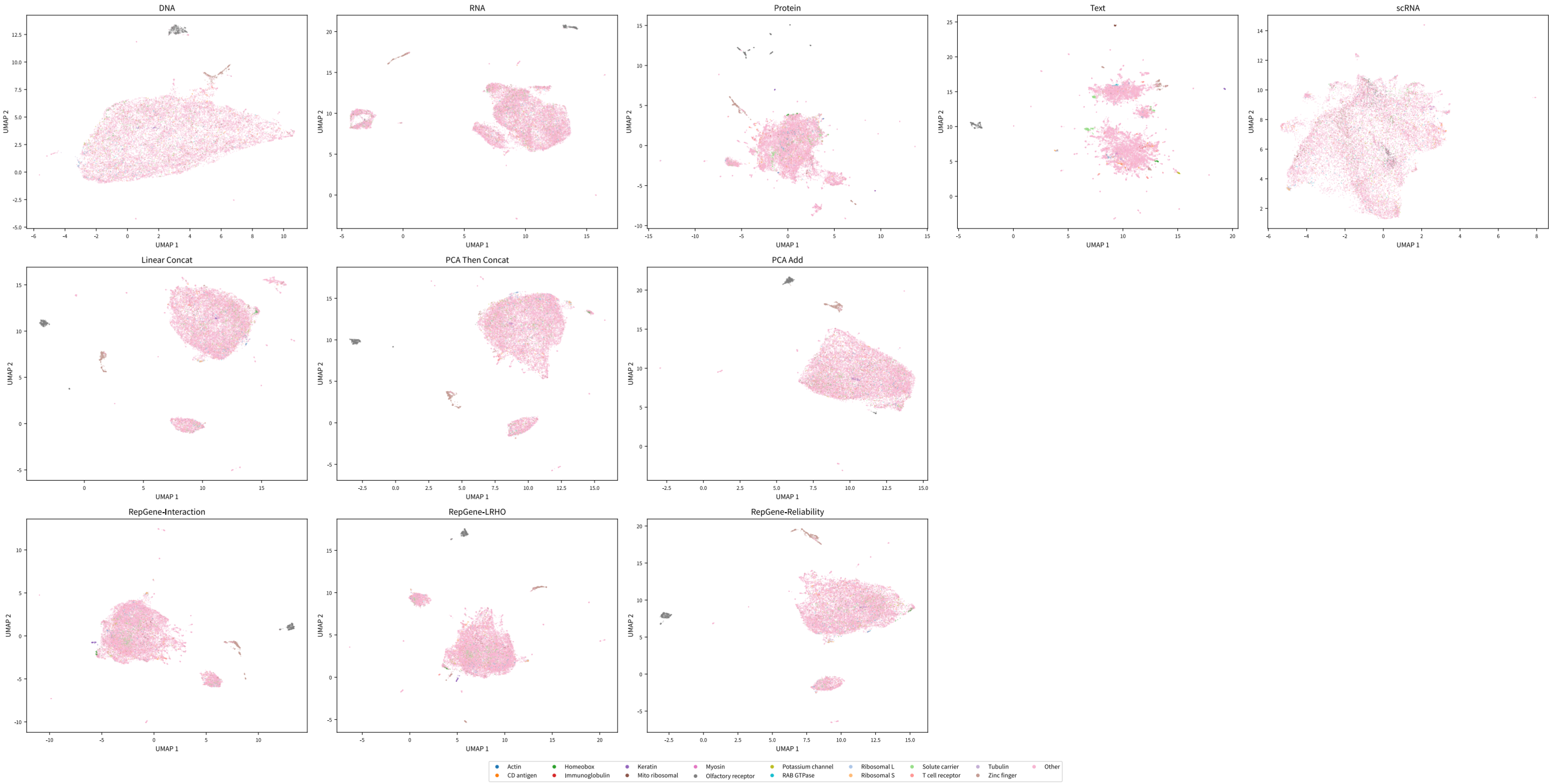

### Figure_1_four_questions.pdf

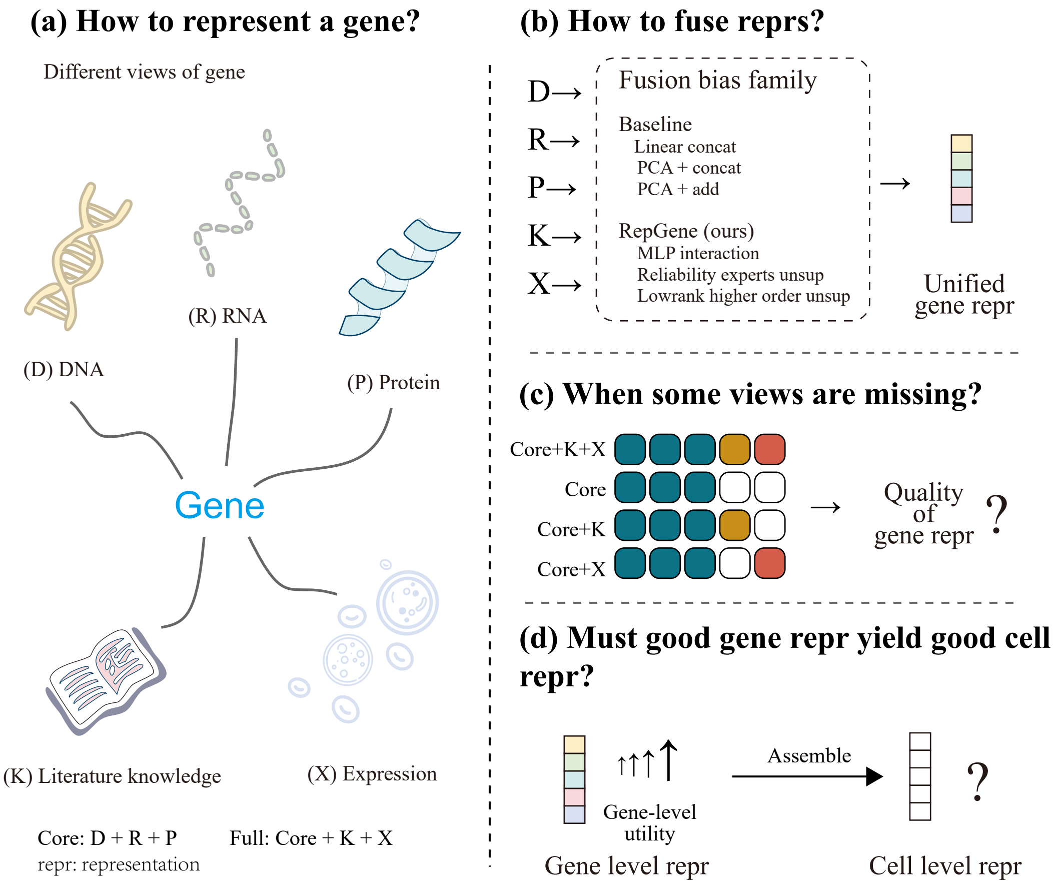

### Figure_1_task_method_condition_heatmap.pdf

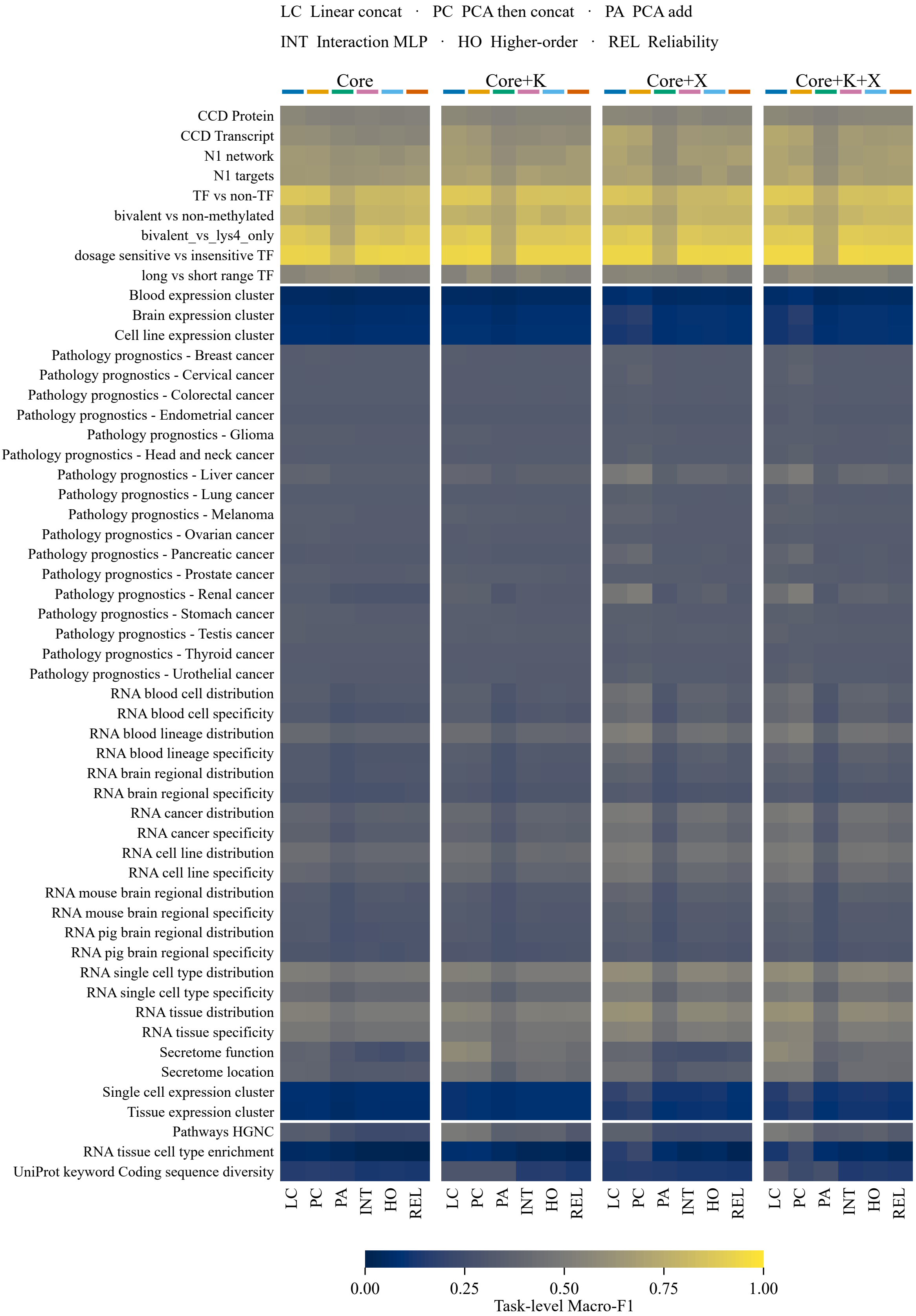

### Figure_2_task_equal_condition_curves.pdf

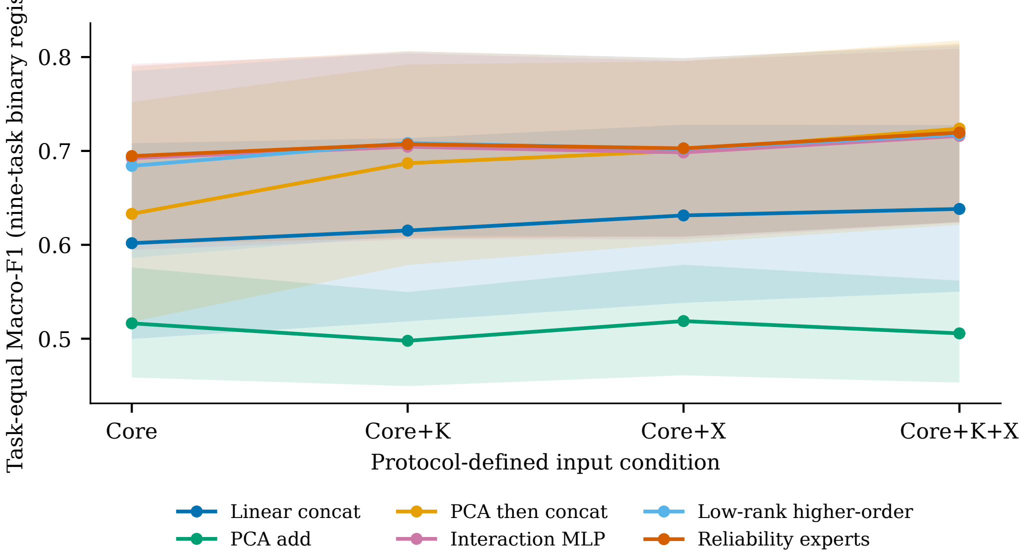

### Figure_2_task_equal_condition_curves.png

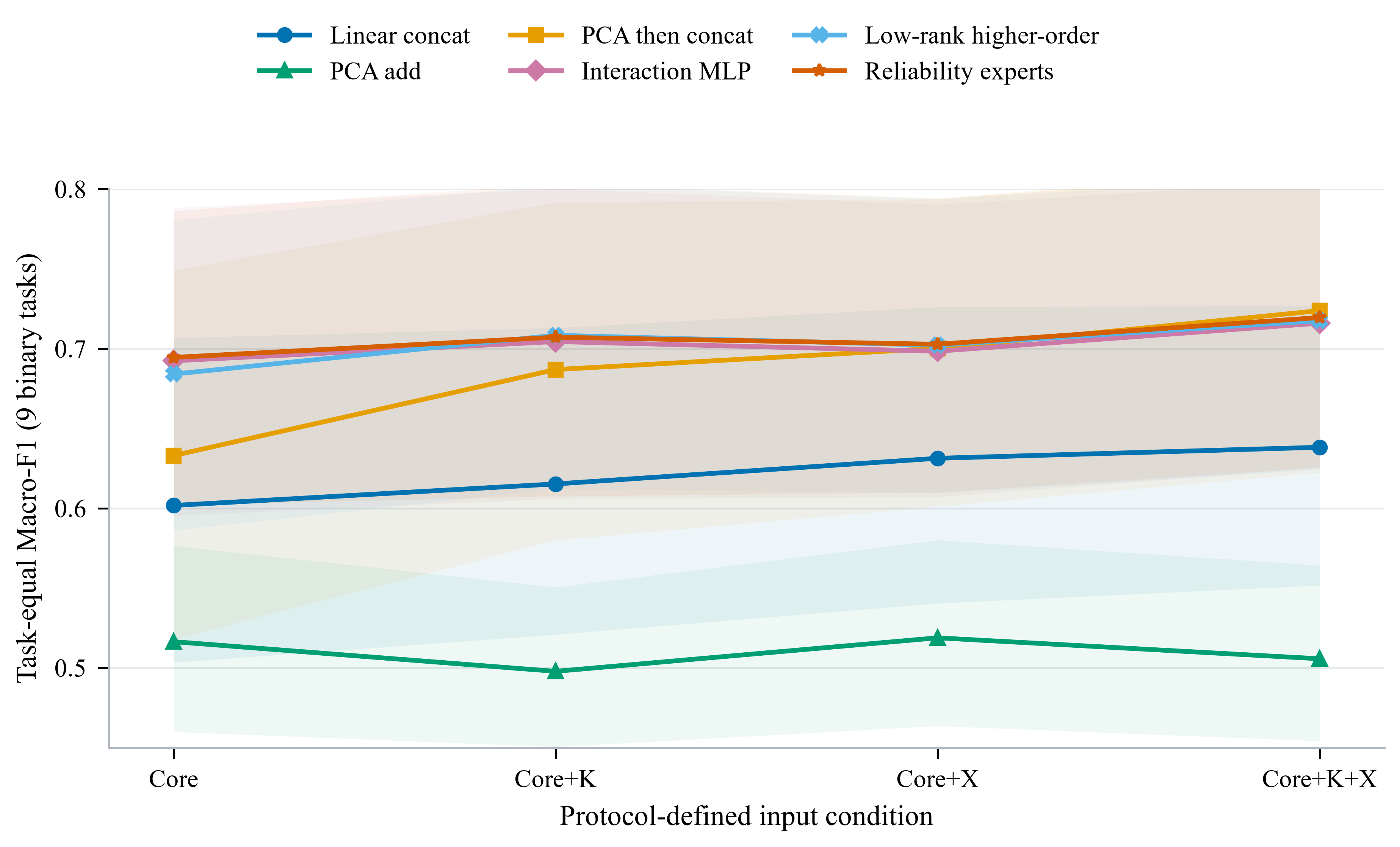

### Figure_3_paired_condition_effects.pdf

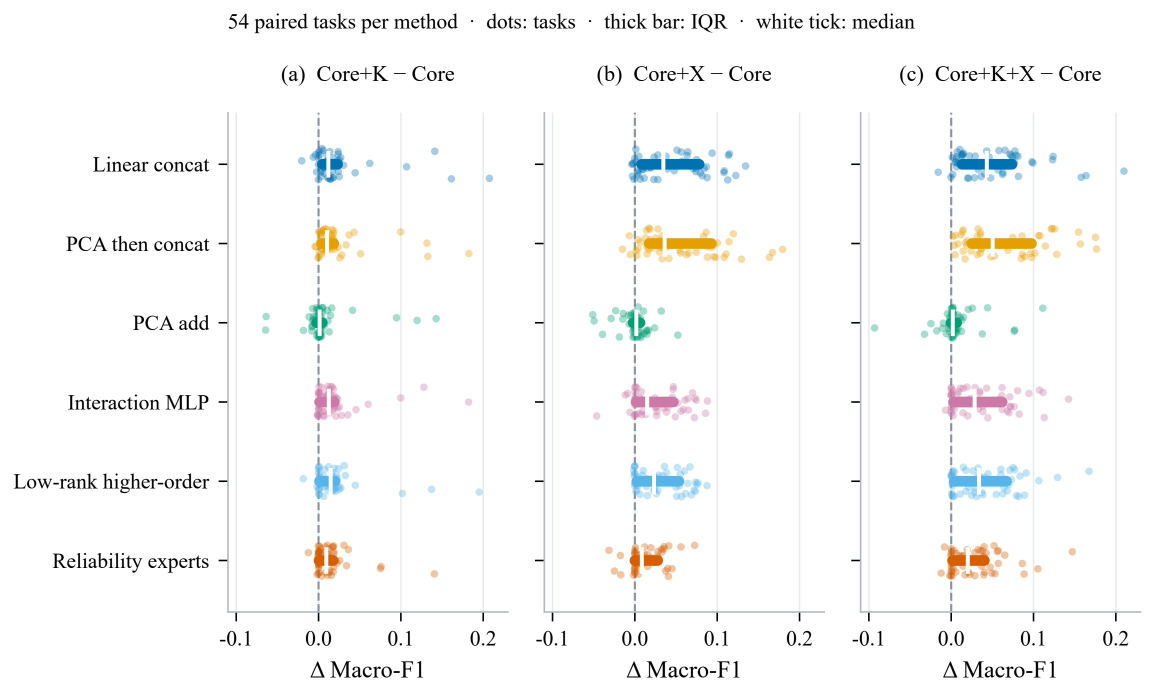

### Figure_3_paired_condition_effects.png

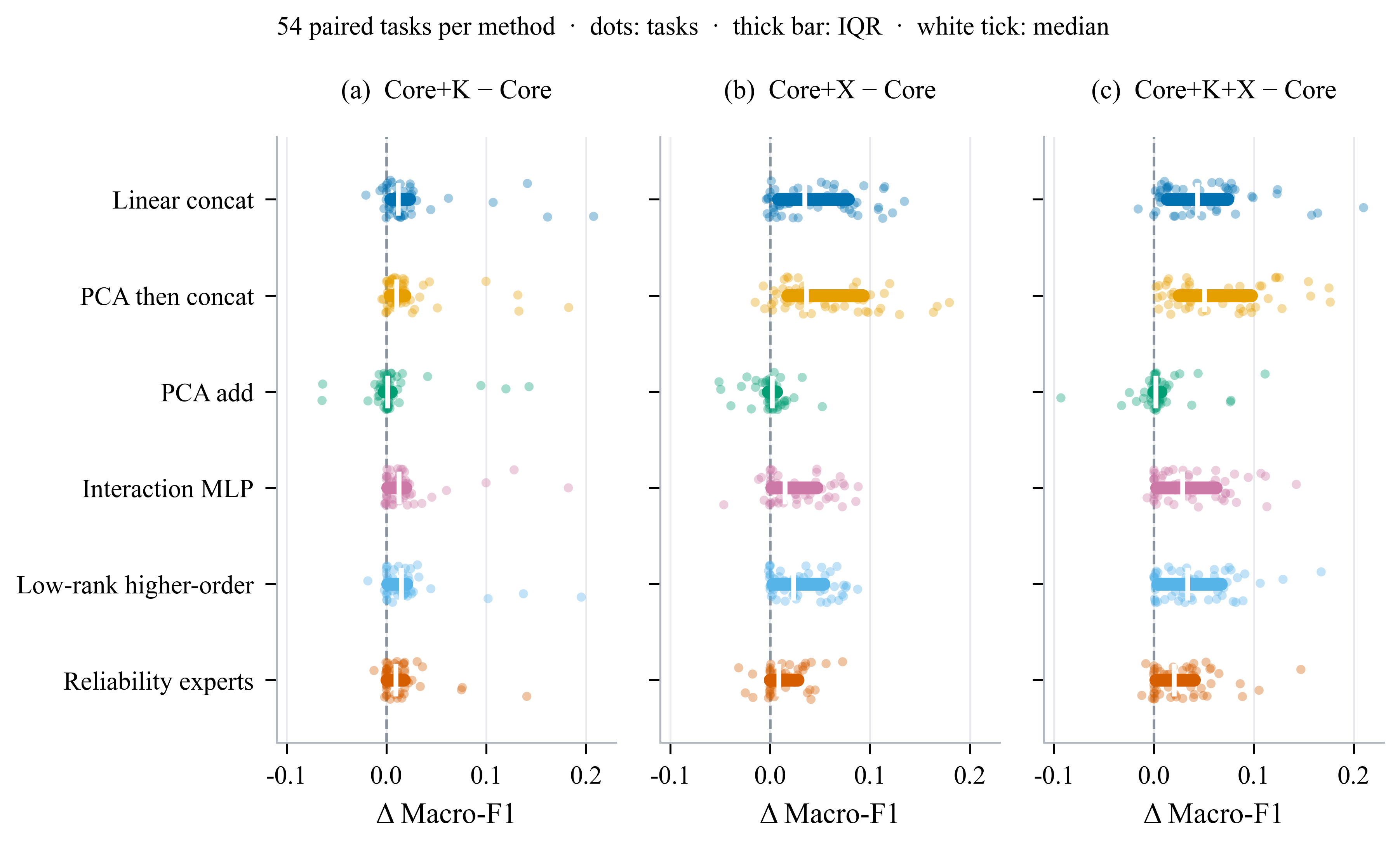

### Figure_3_task_method_condition_heatmap.pdf

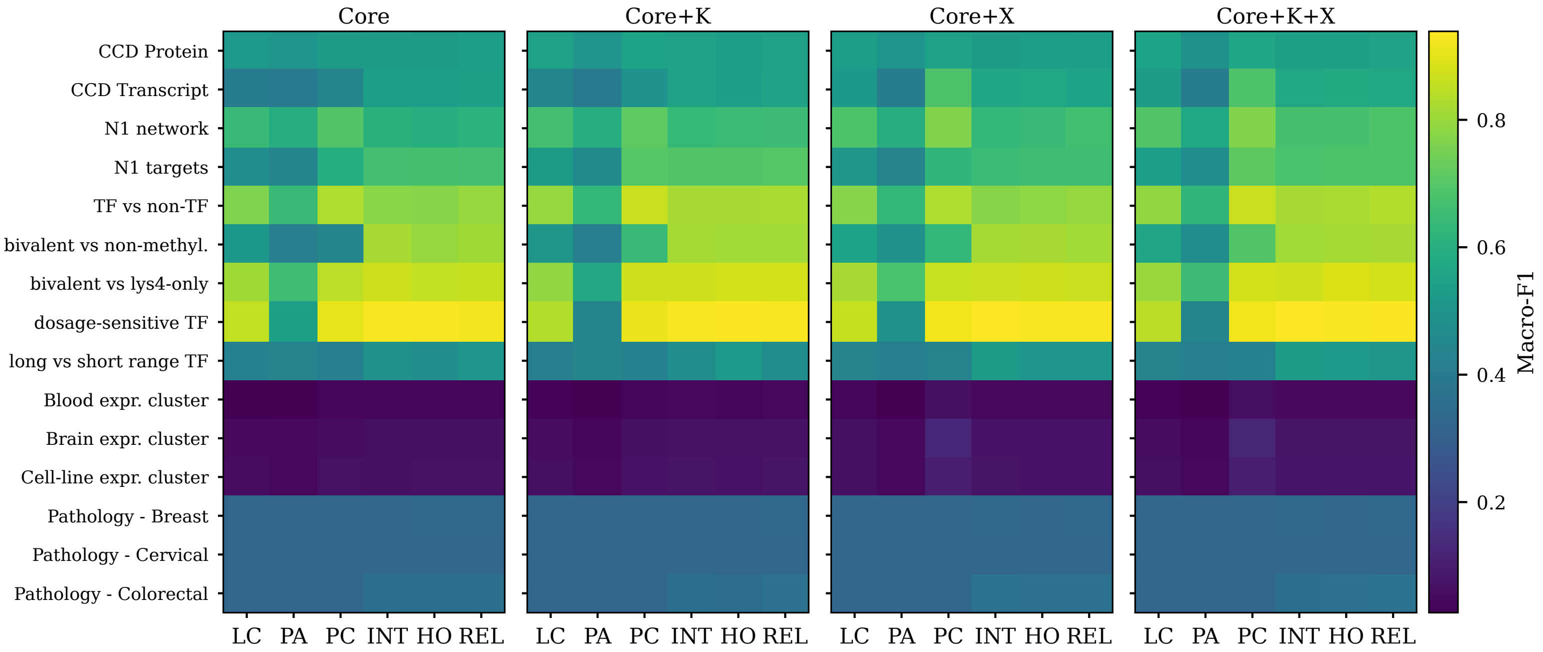

### Figure_3_task_method_condition_heatmap.png

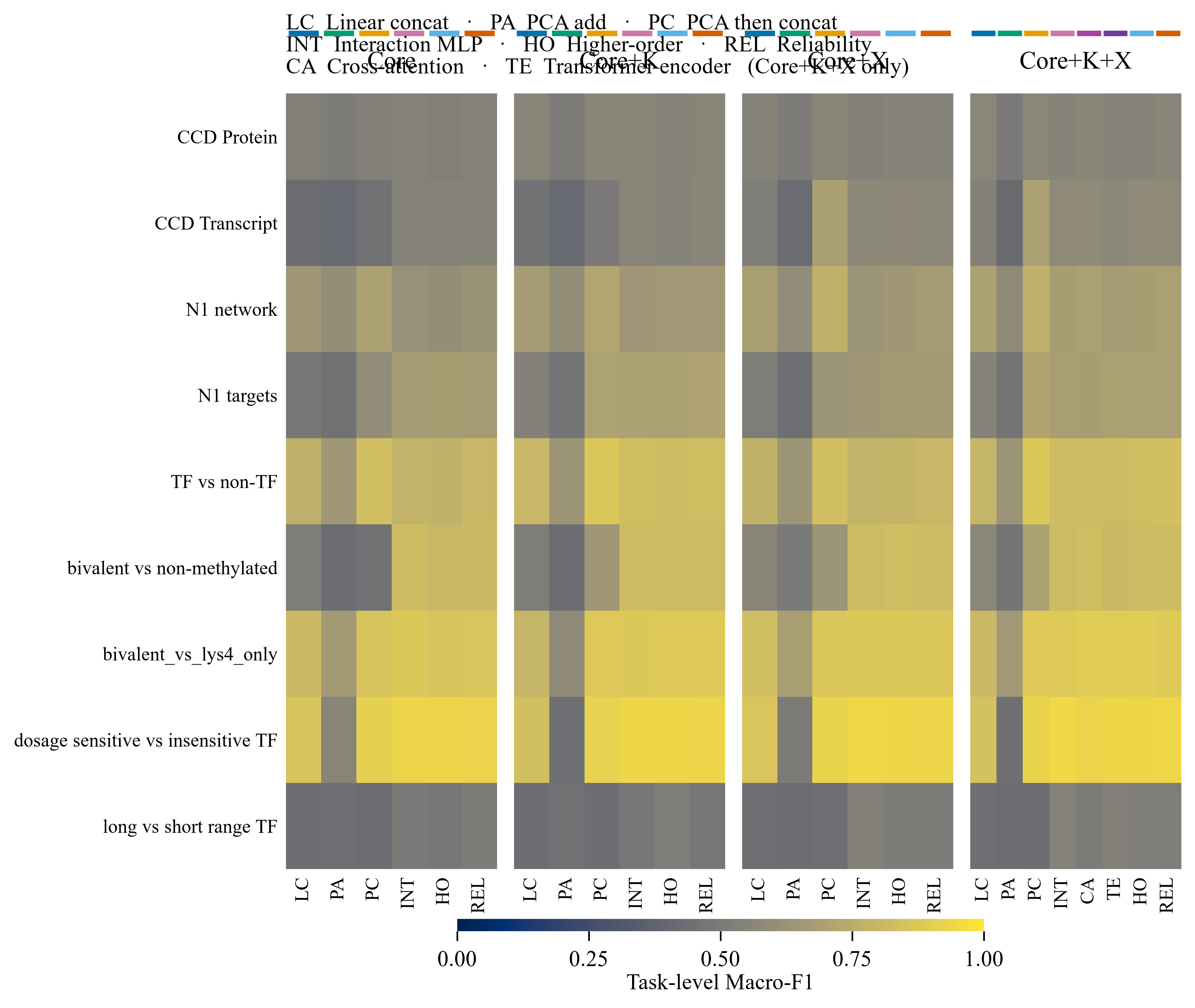

### Figure_4_method_rank_and_wins.pdf

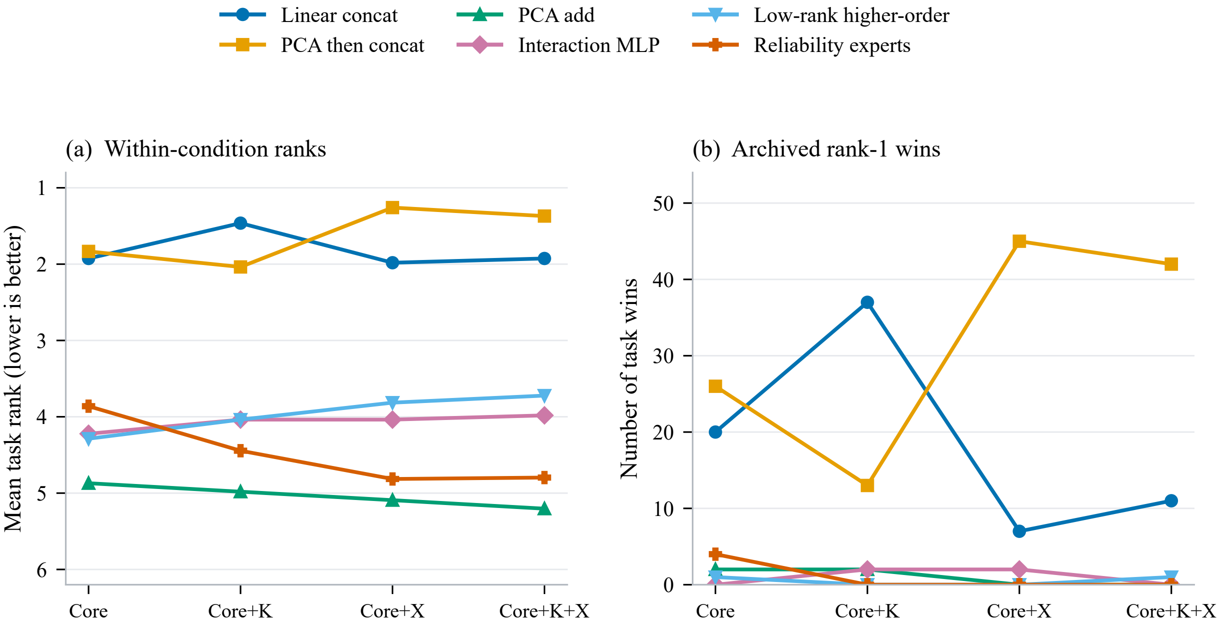

### Figure_4_method_rank_and_wins.png

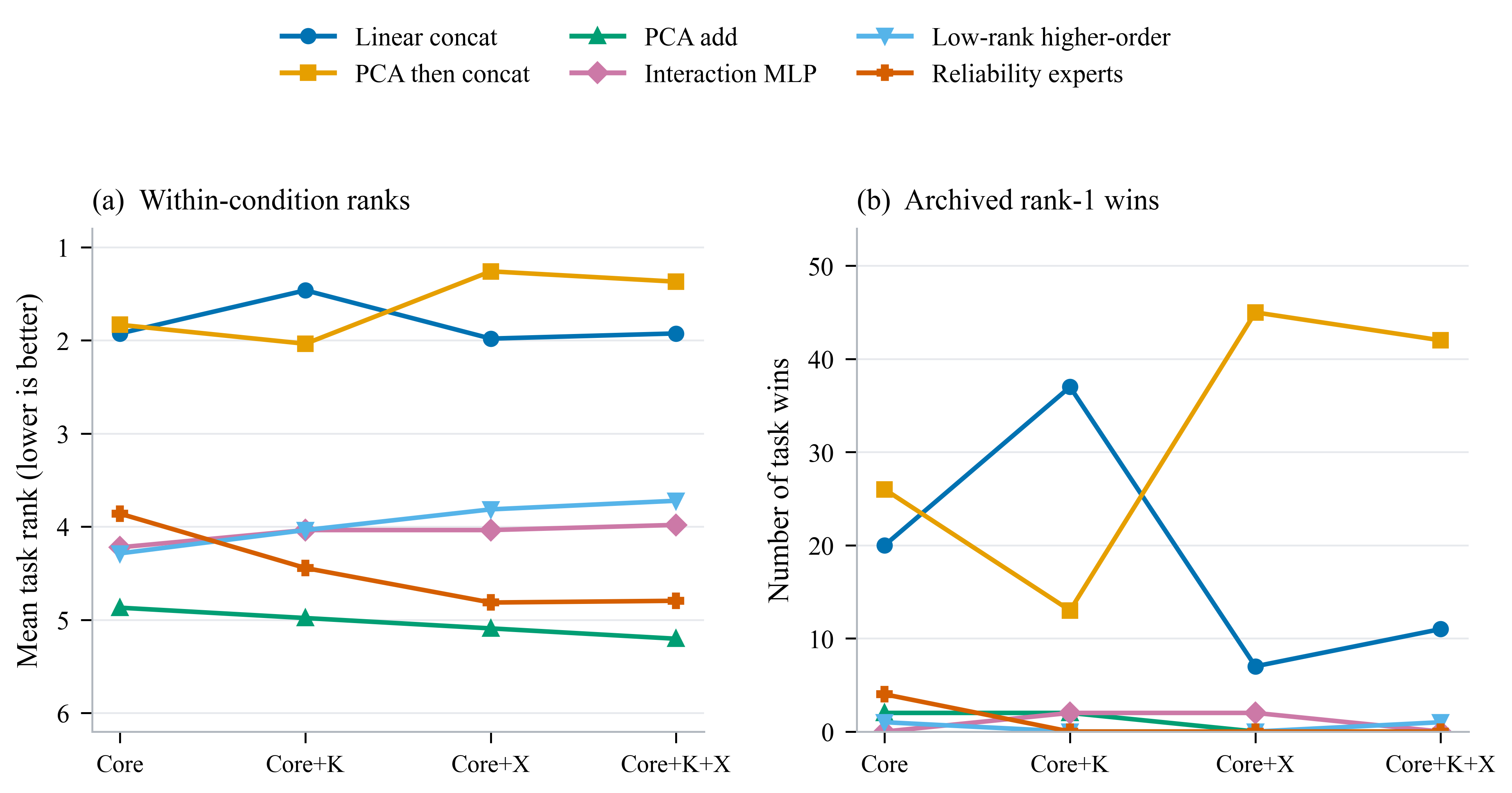

### Figure_4_paired_condition_effects.pdf

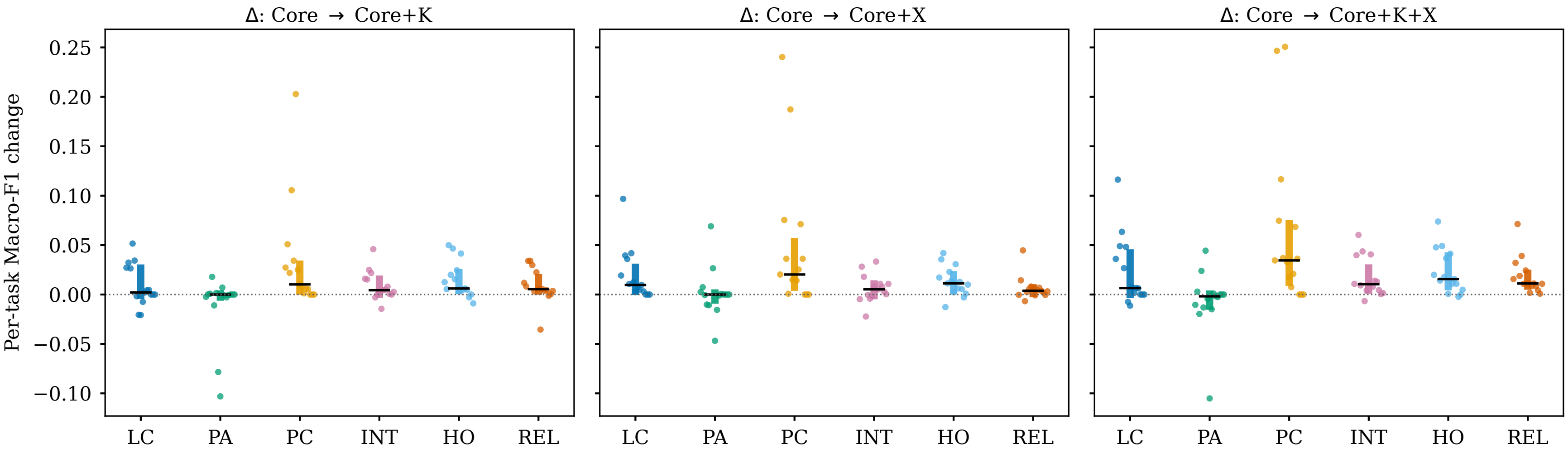

### Figure_4_paired_condition_effects.png

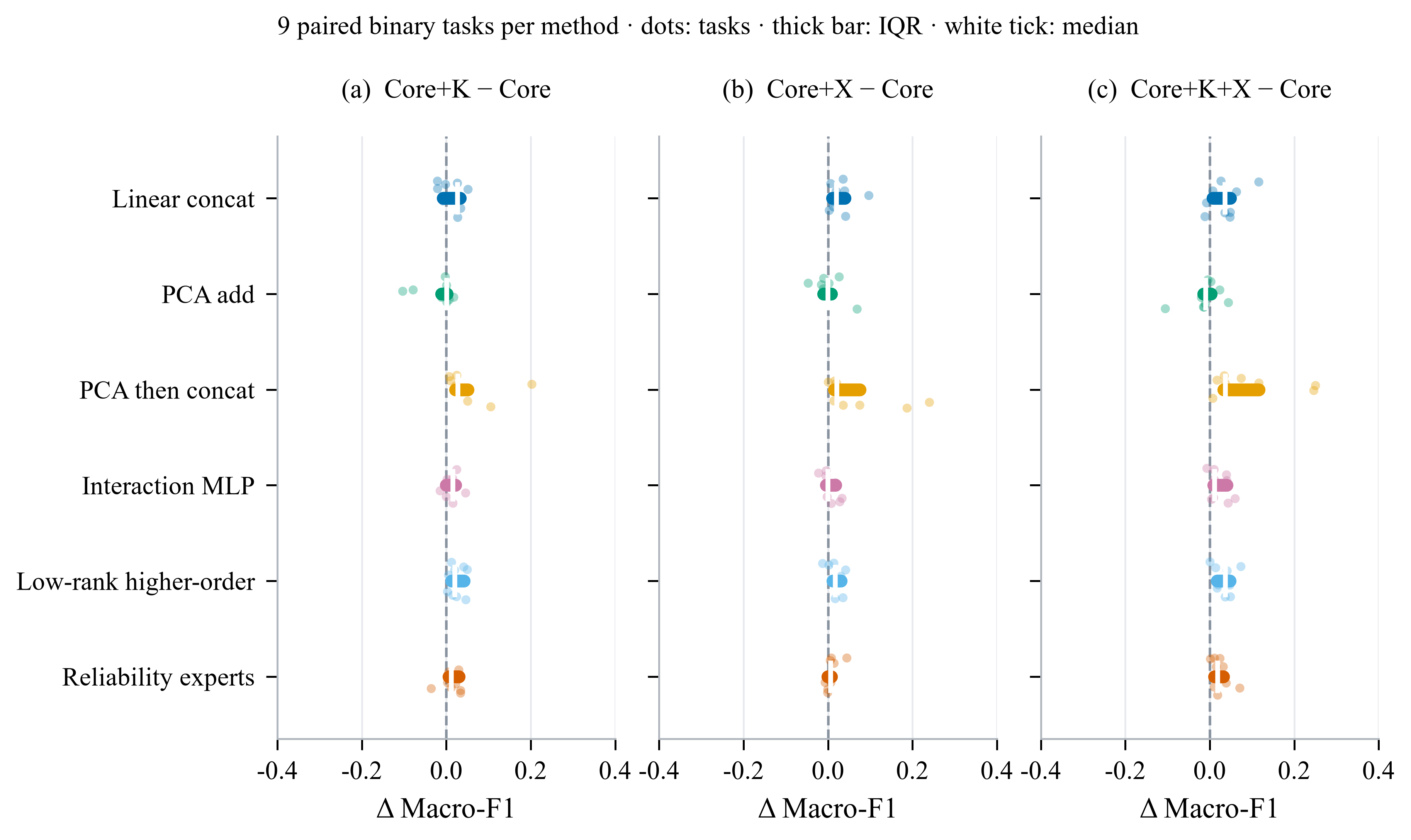

### Figure_5_method_rank_and_wins.pdf

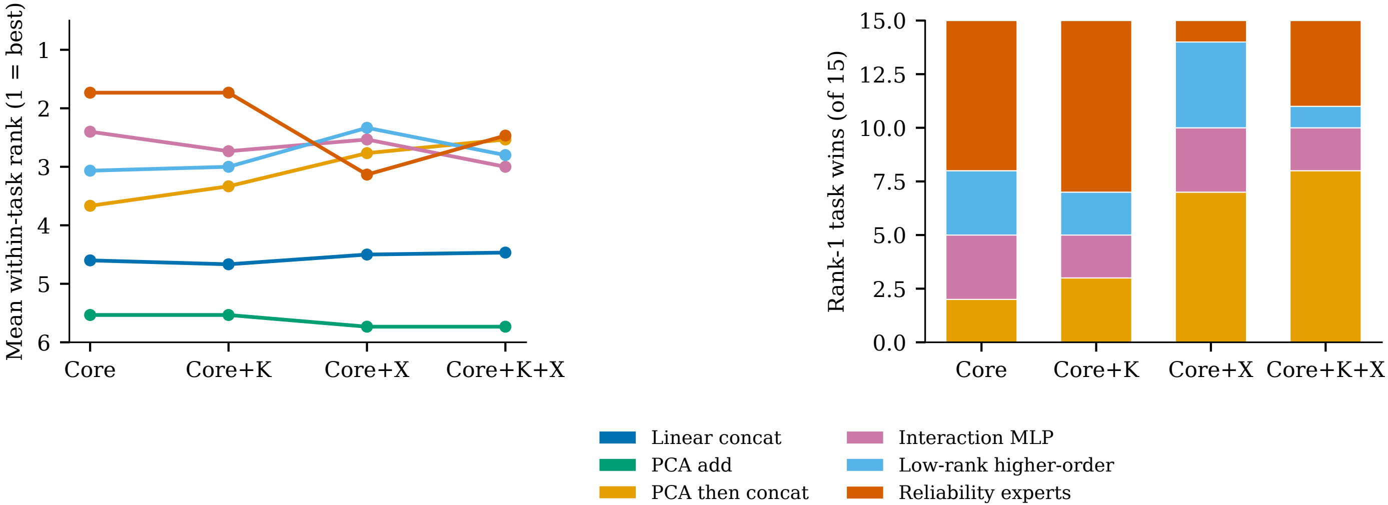

### Figure_5_method_rank_and_wins.png

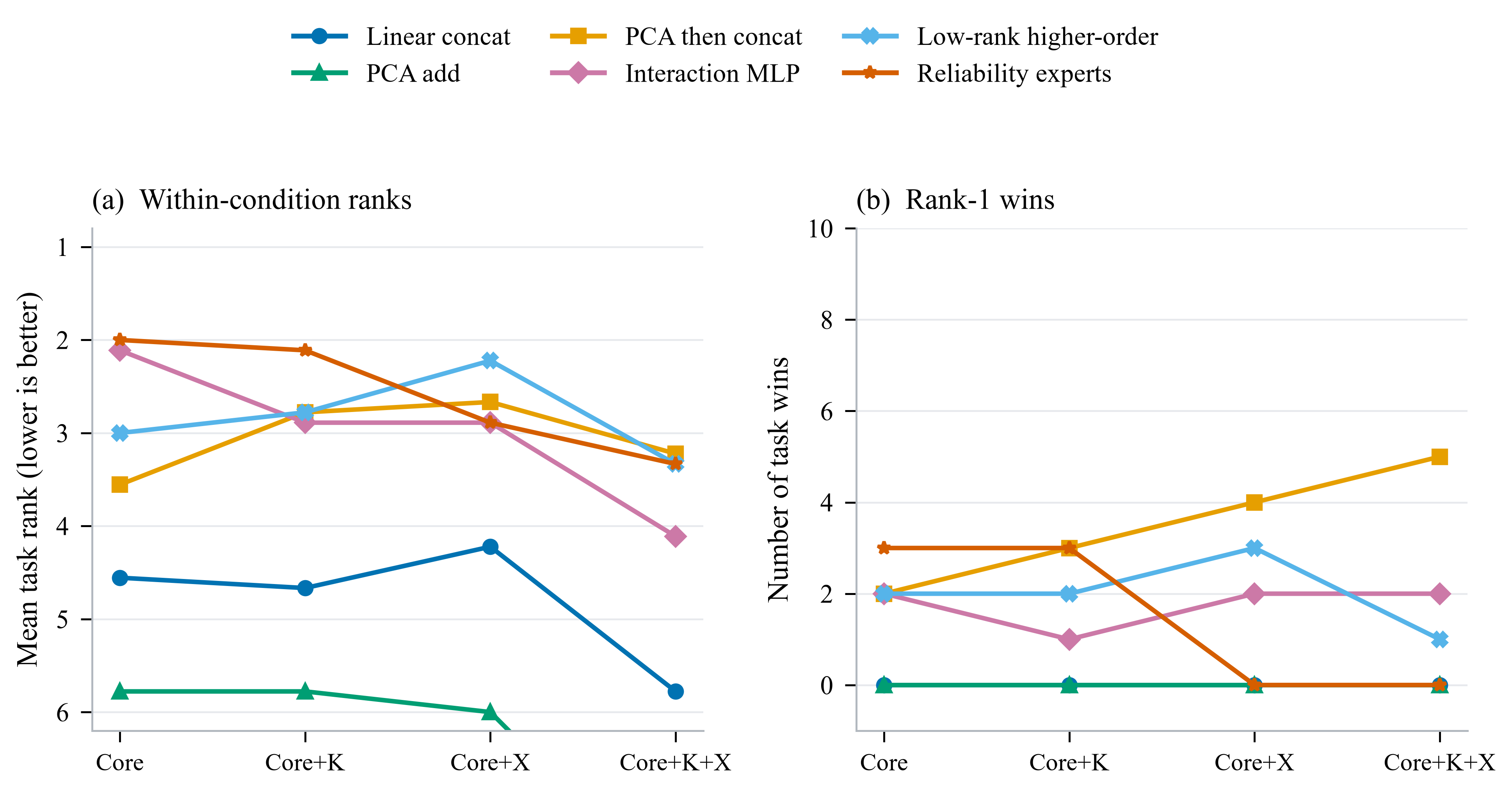

### Figure_5_task_type_stratified_profiles.pdf

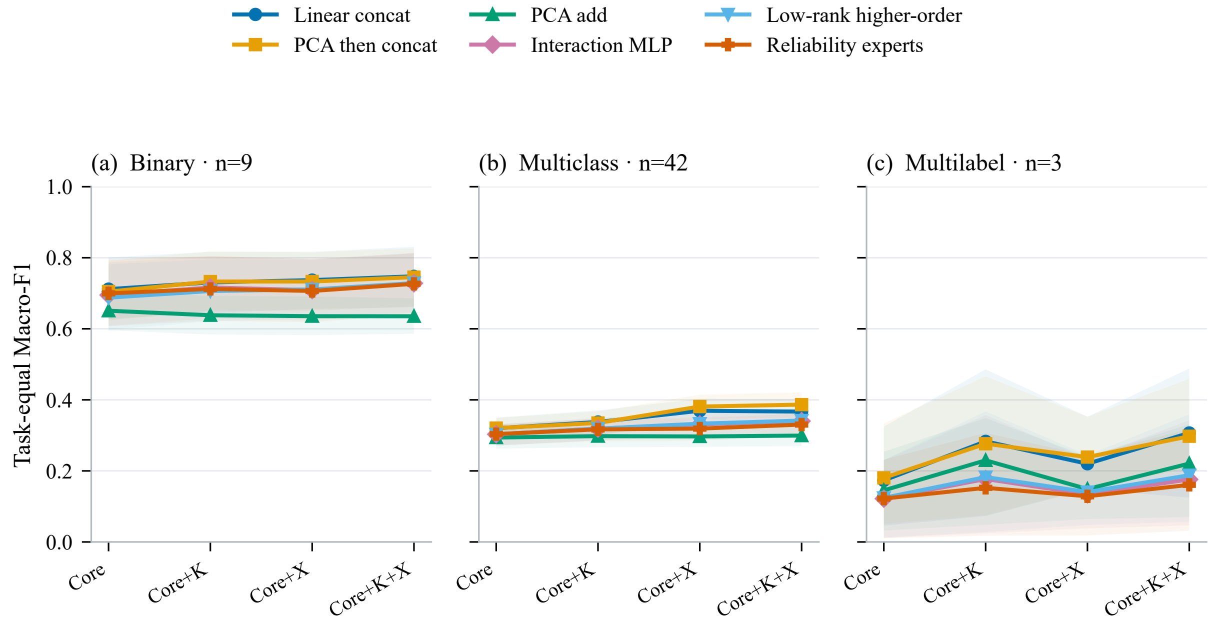

### Figure_5_task_type_stratified_profiles.png

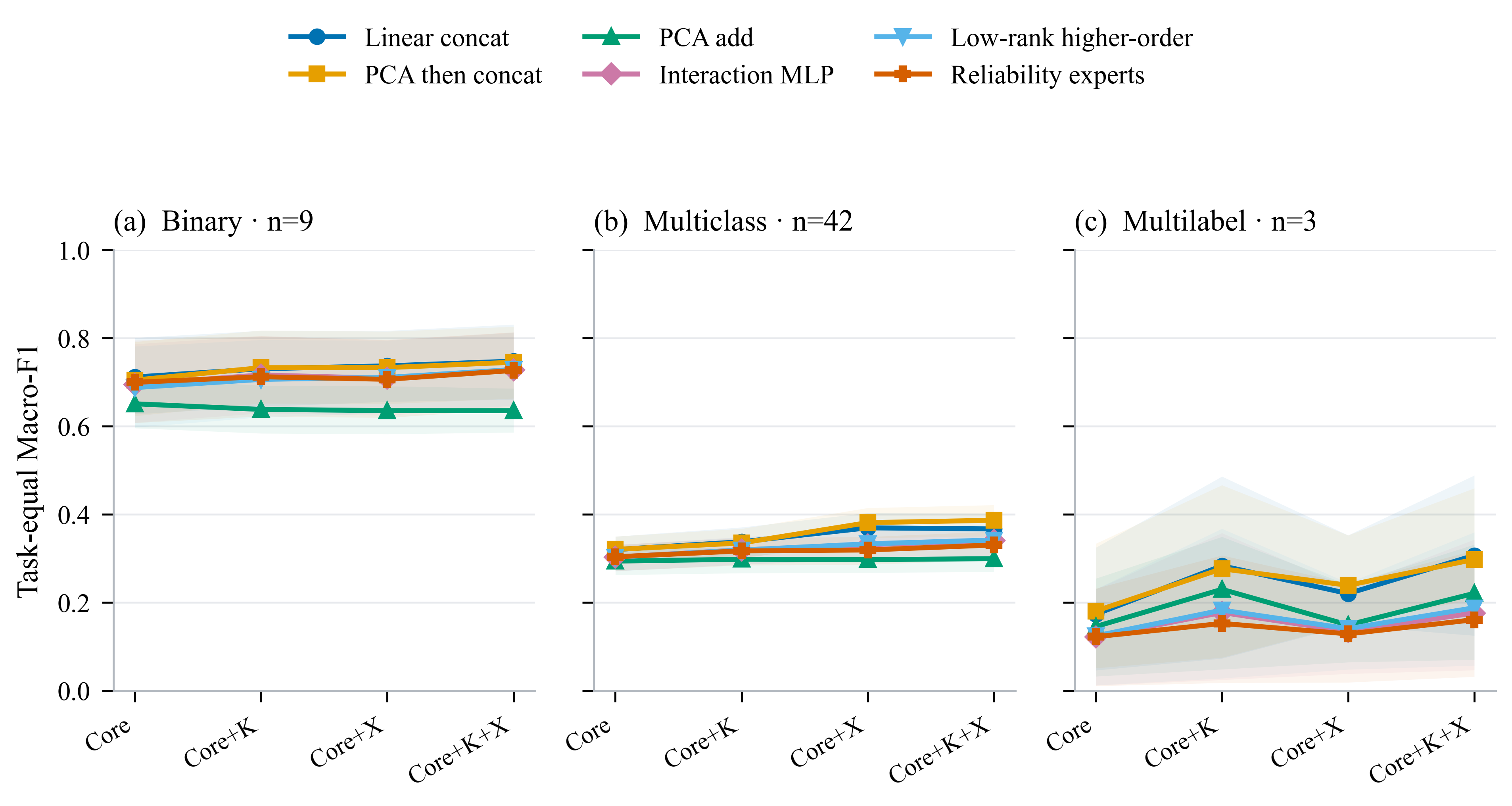

### Figure_6_binary_suite_profiles.pdf

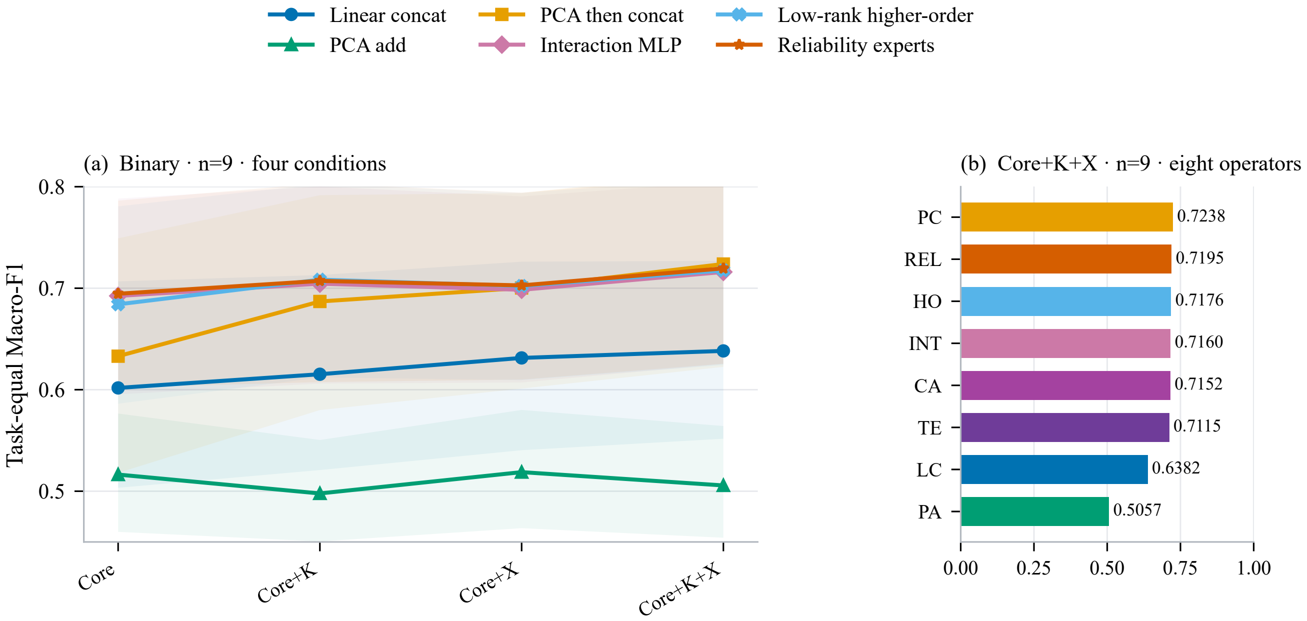

### Figure_6_binary_suite_profiles.png

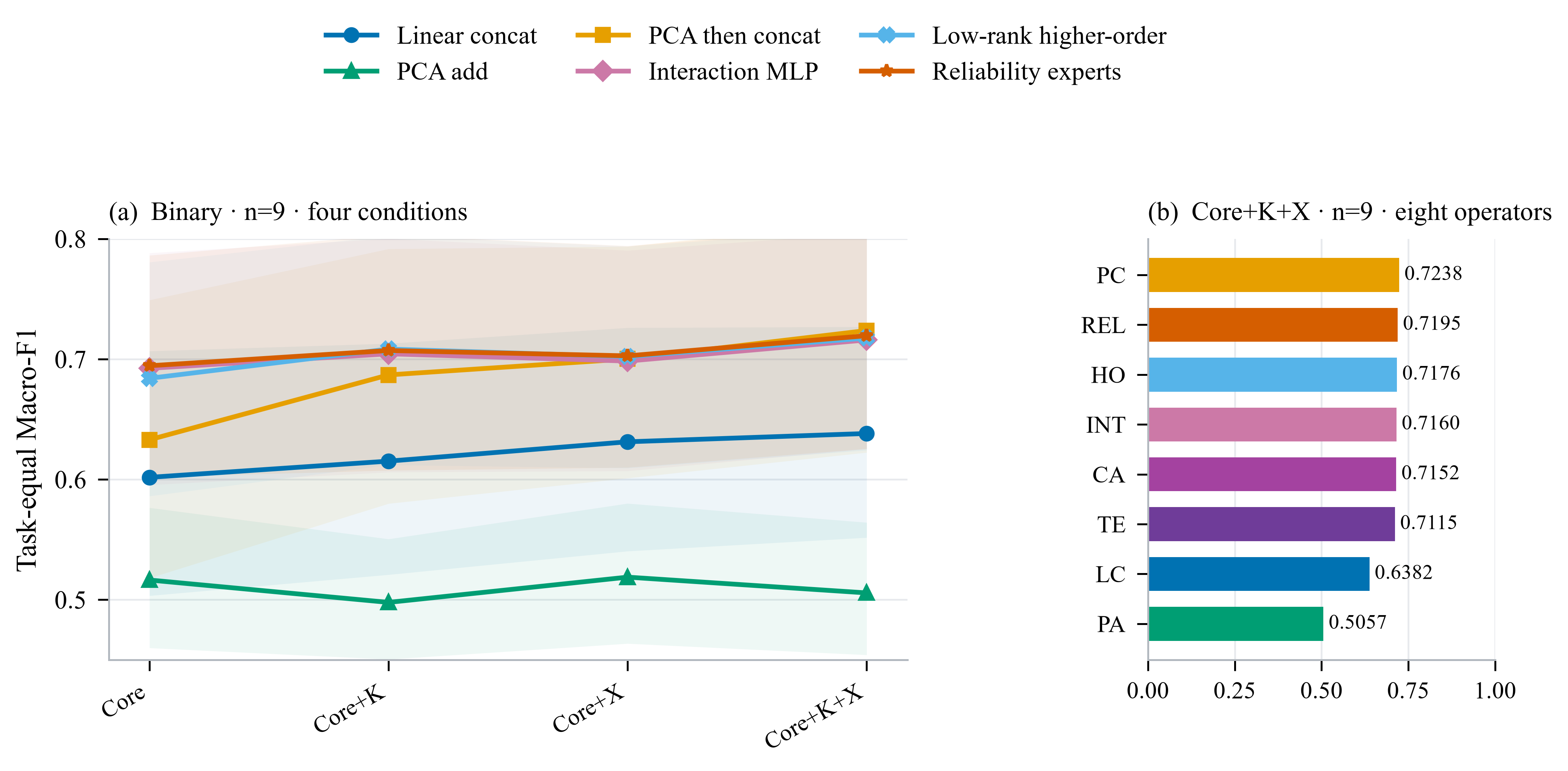

### Figure_6_KX_nonadditivity.pdf

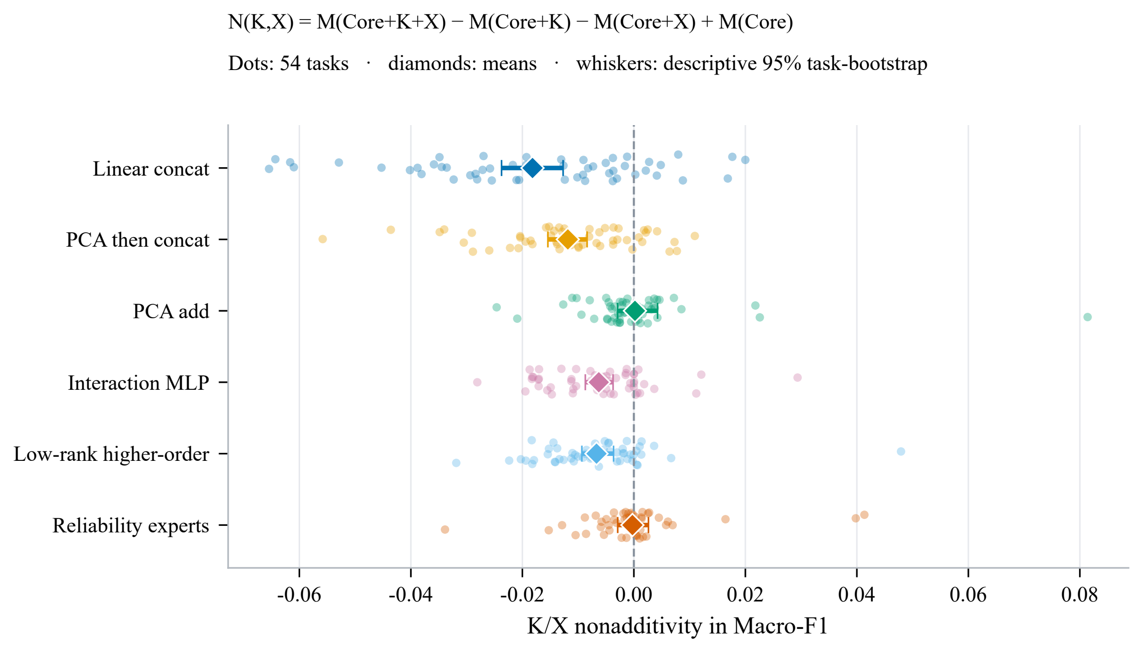

### Figure_6_KX_nonadditivity.png

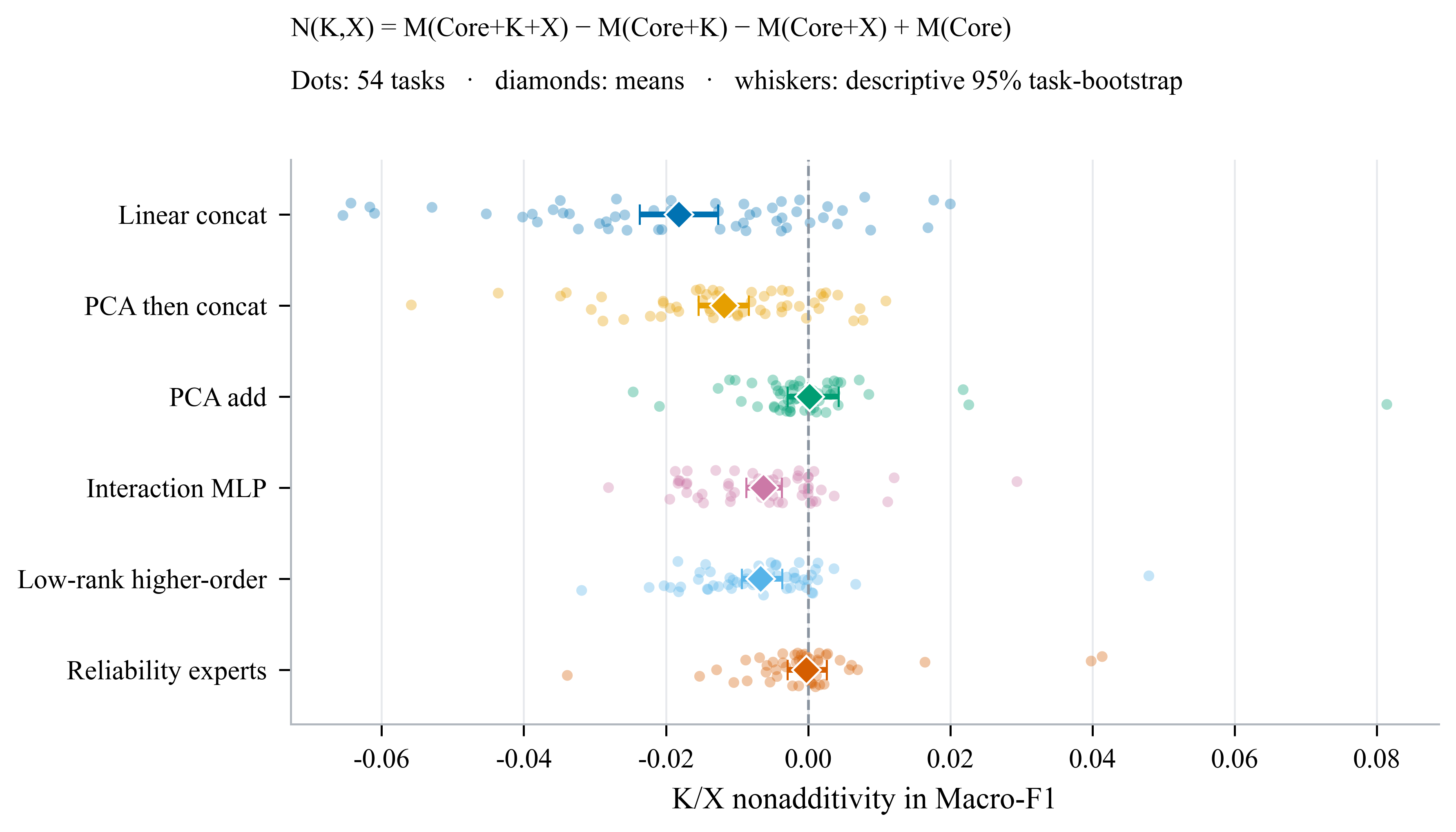

### Figure_7_KX_nonadditivity.pdf

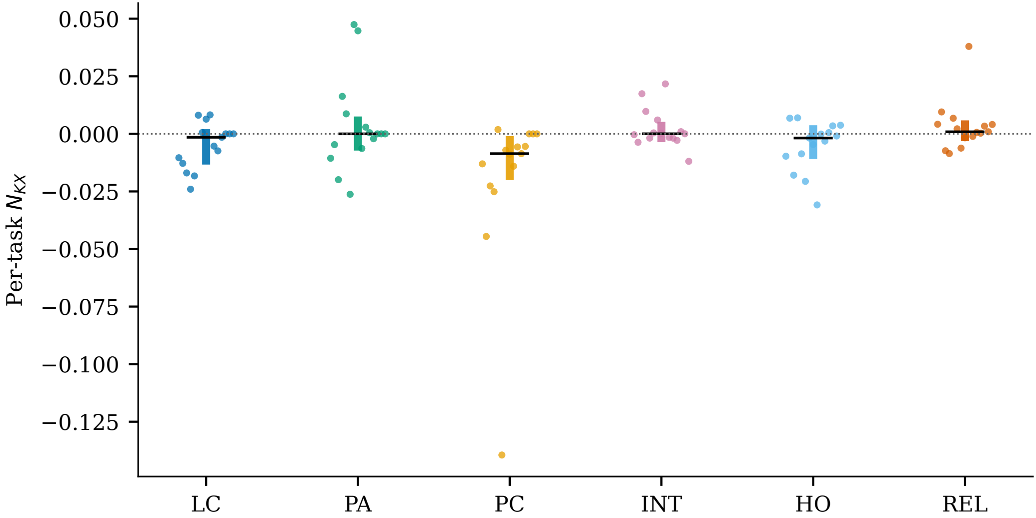

### Figure_7_KX_nonadditivity.png

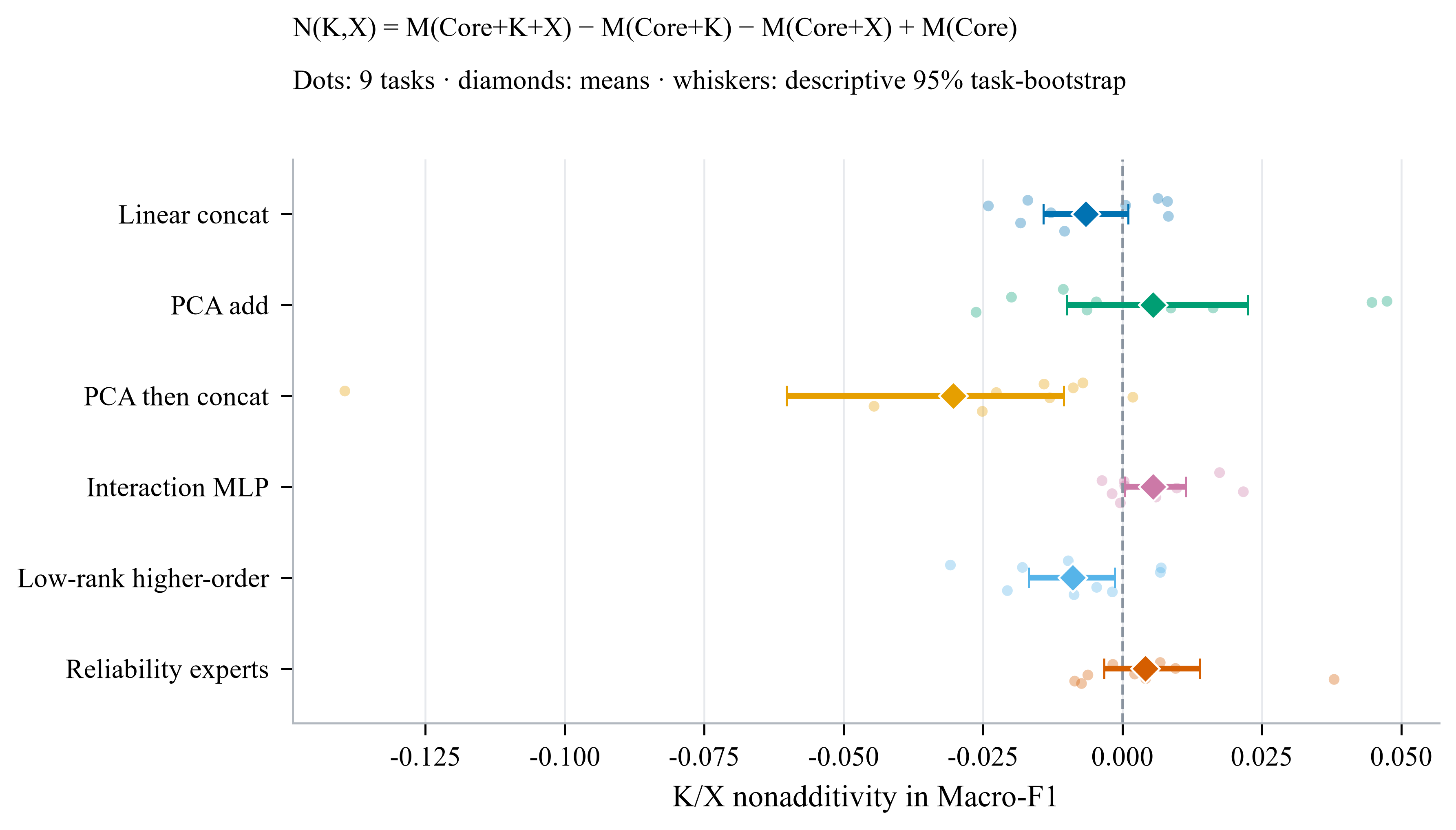

### Figure_8_external_benchmark.pdf

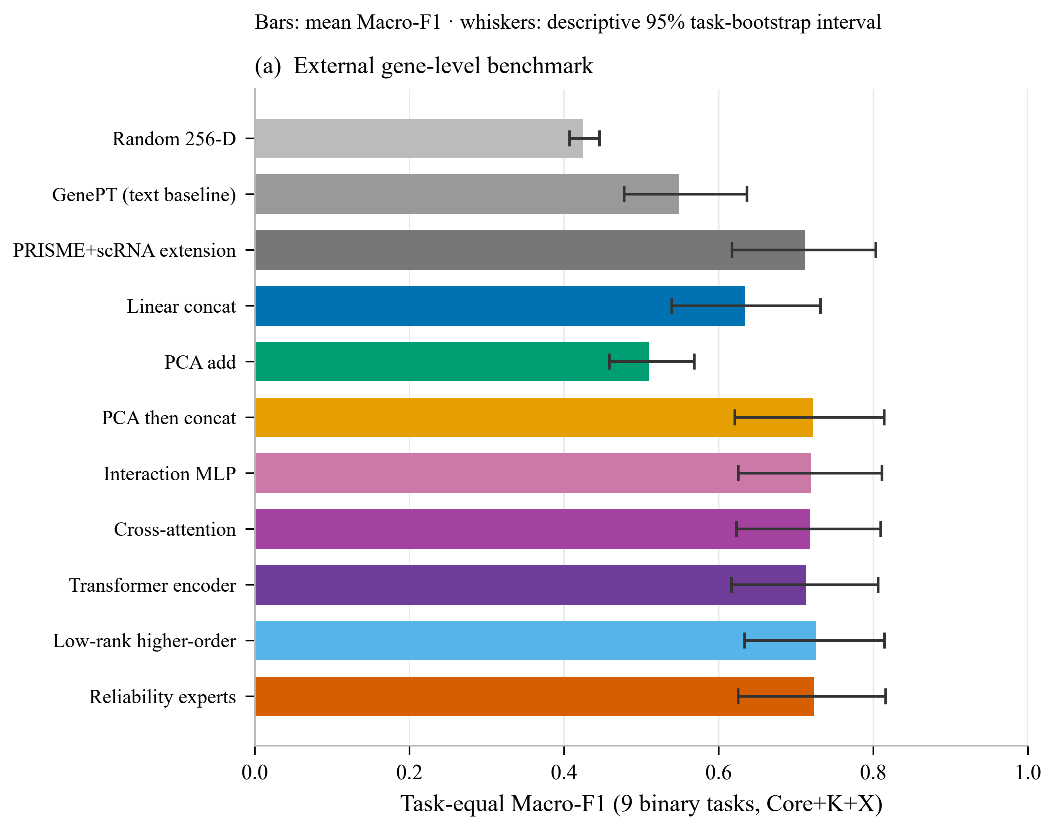

### Figure_8_external_benchmark.png

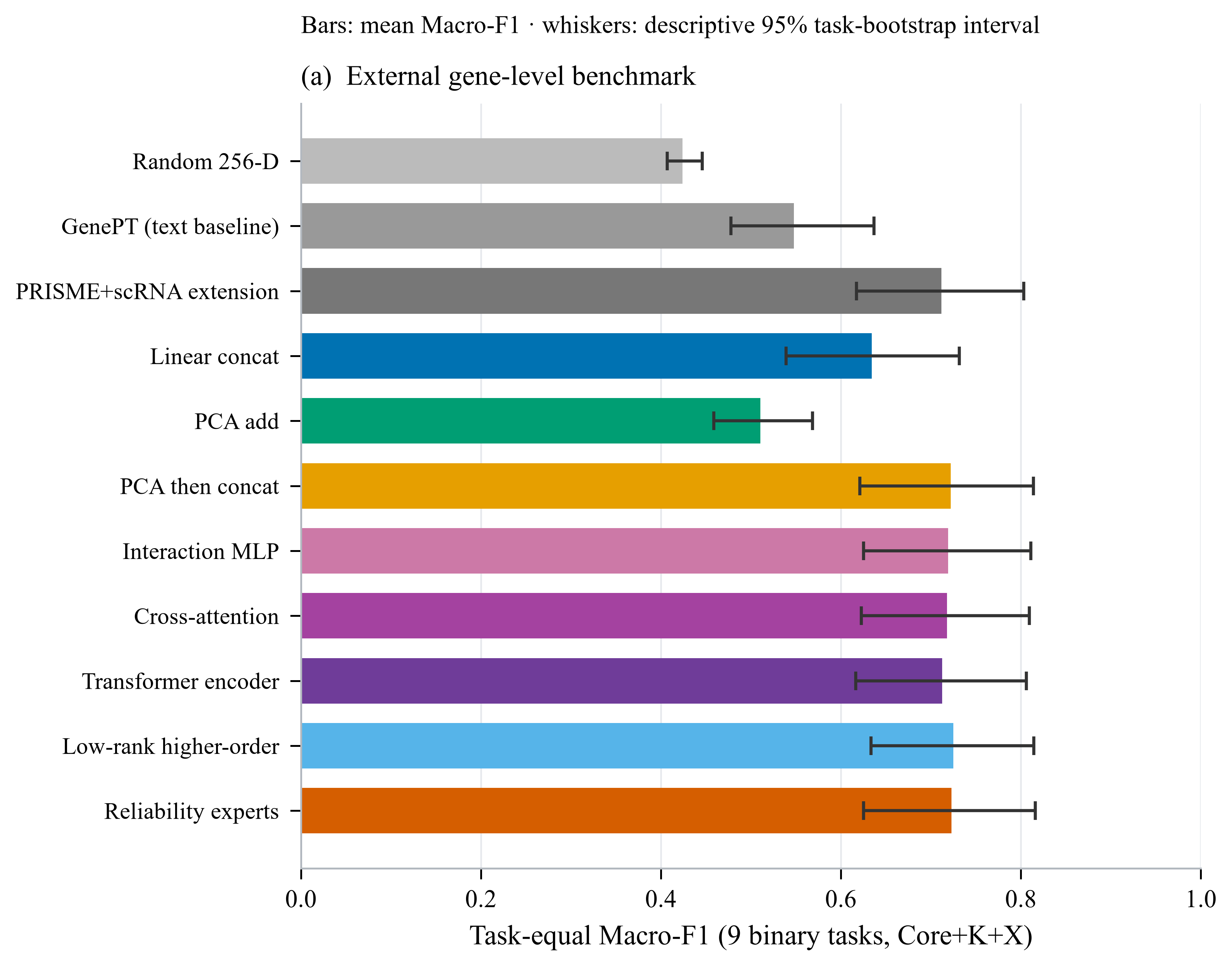

### Figure_framework_overview.pdf

#### RepGene-PRF: Matched Multi-View Fusion-Bias Audit Framework

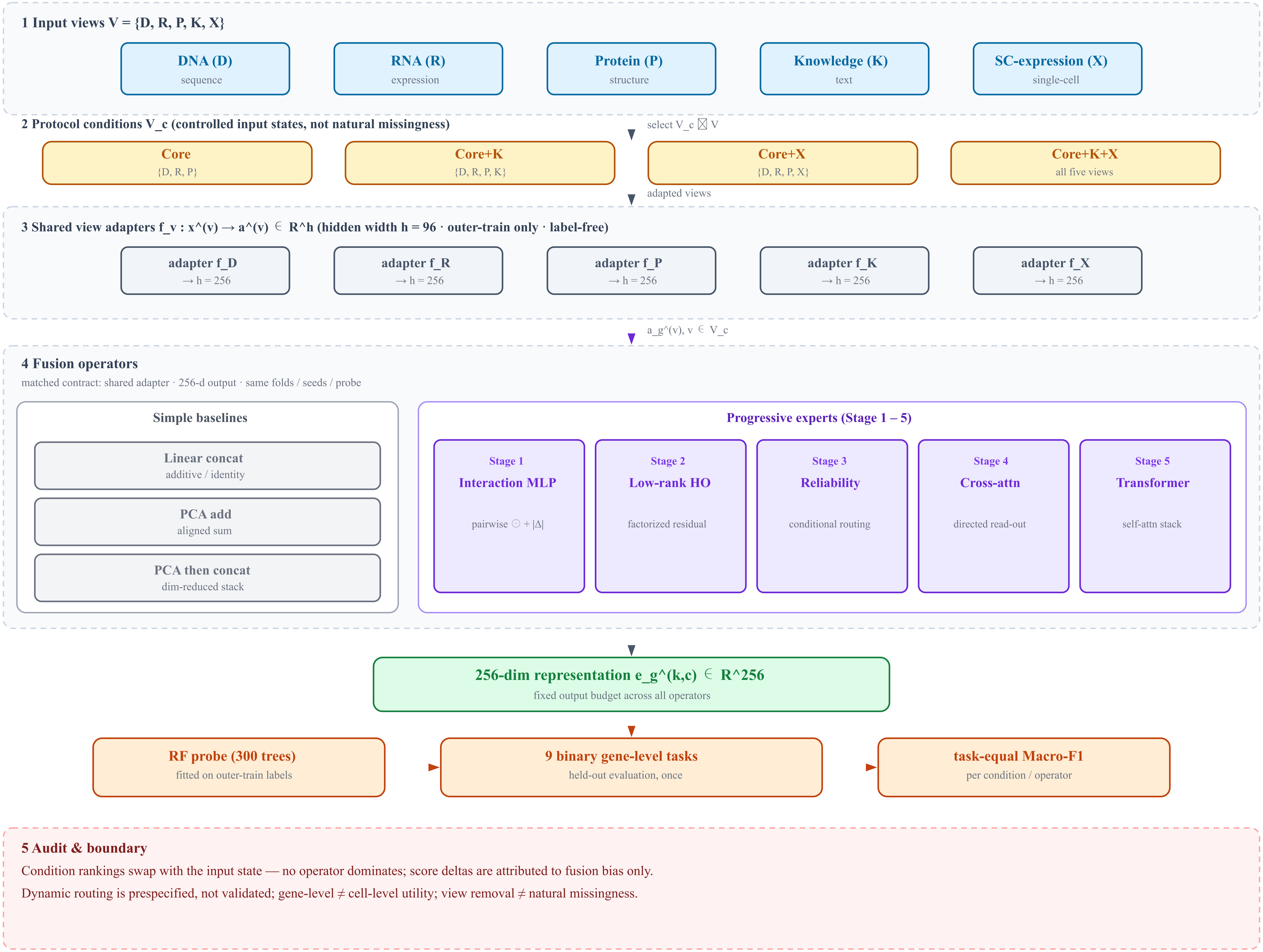

### fusion_umap_comparison_01.pdf

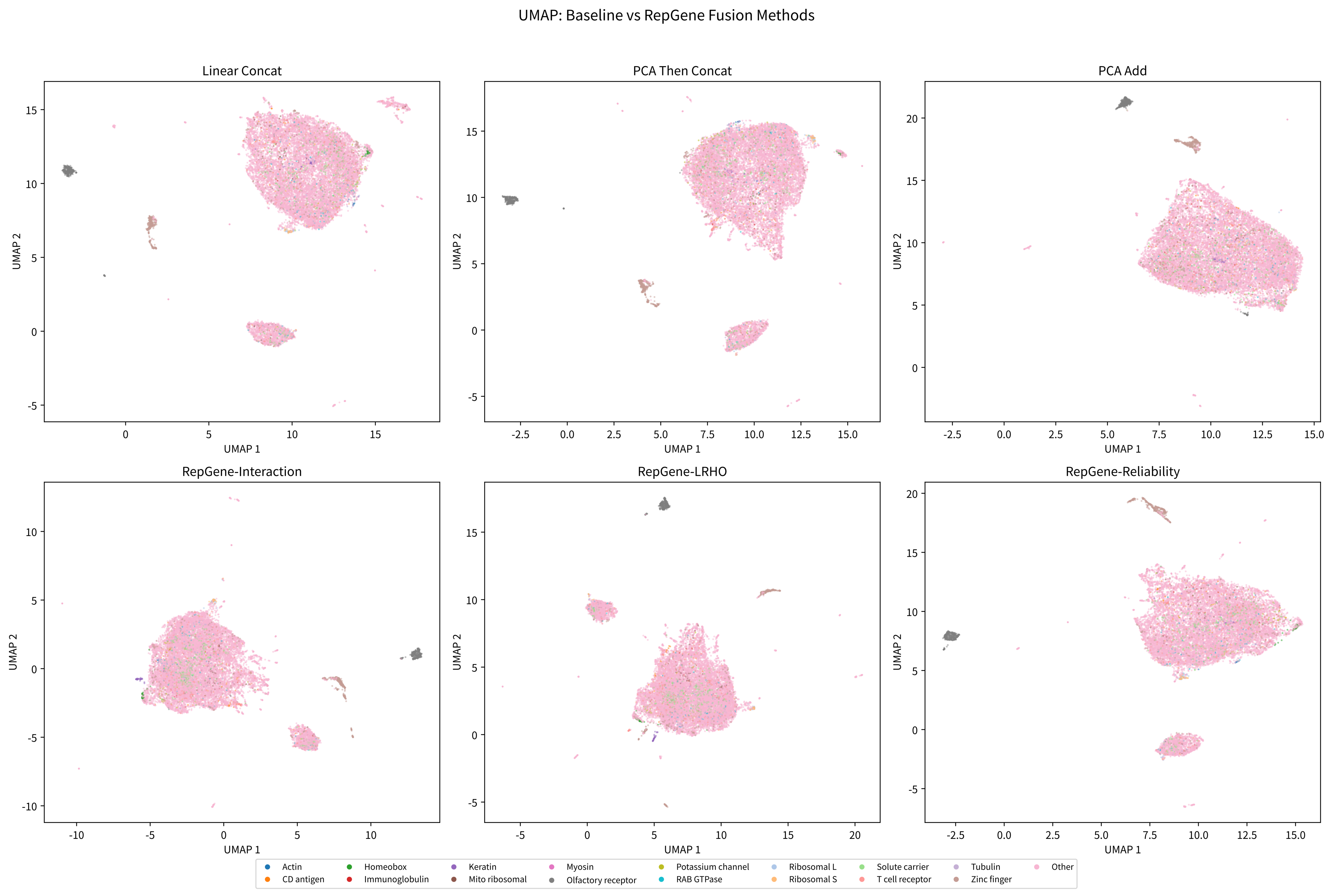

### knn_overlap_all_methods.pdf

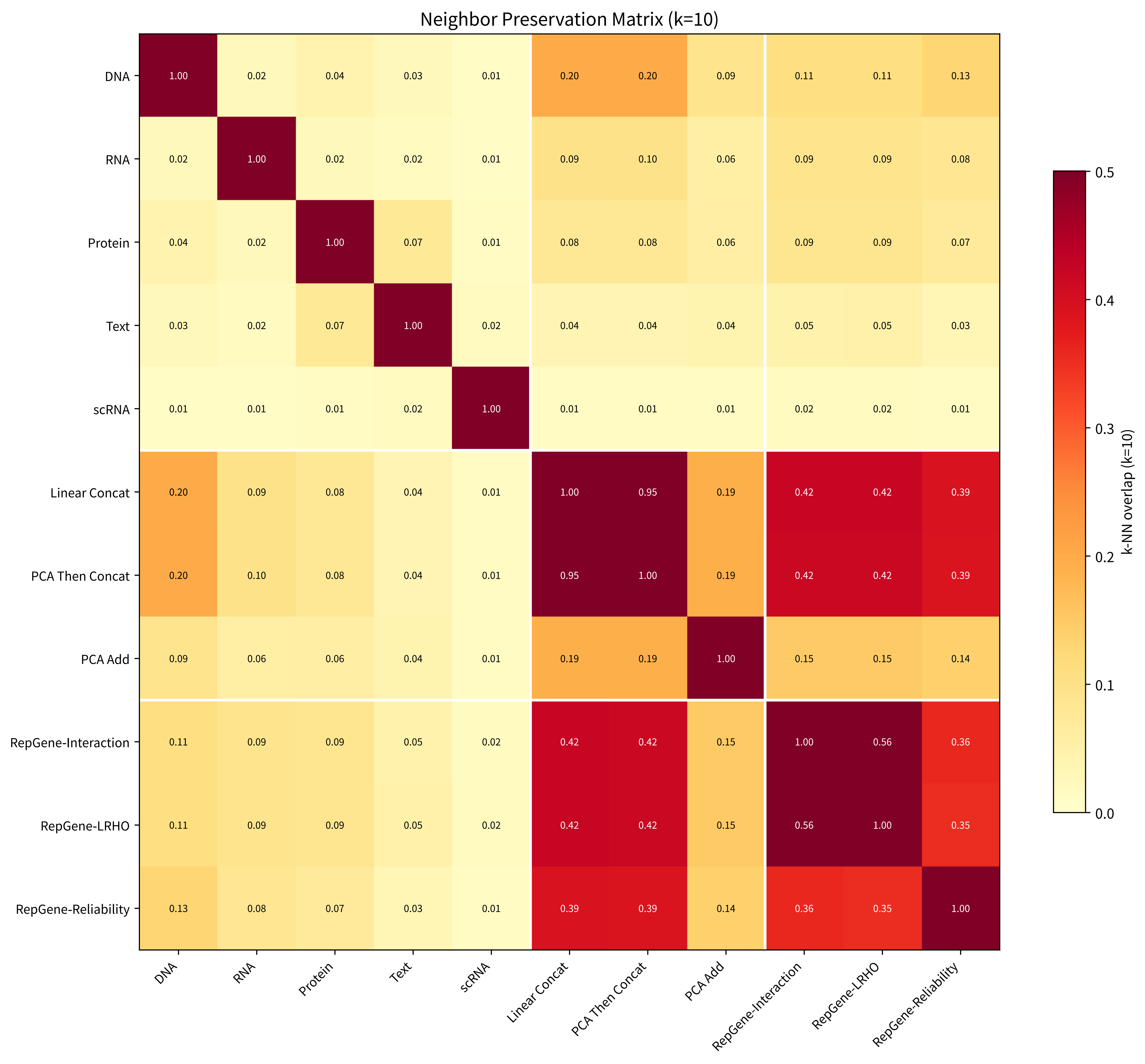

### knn_overlap_k50_heatmap.pdf

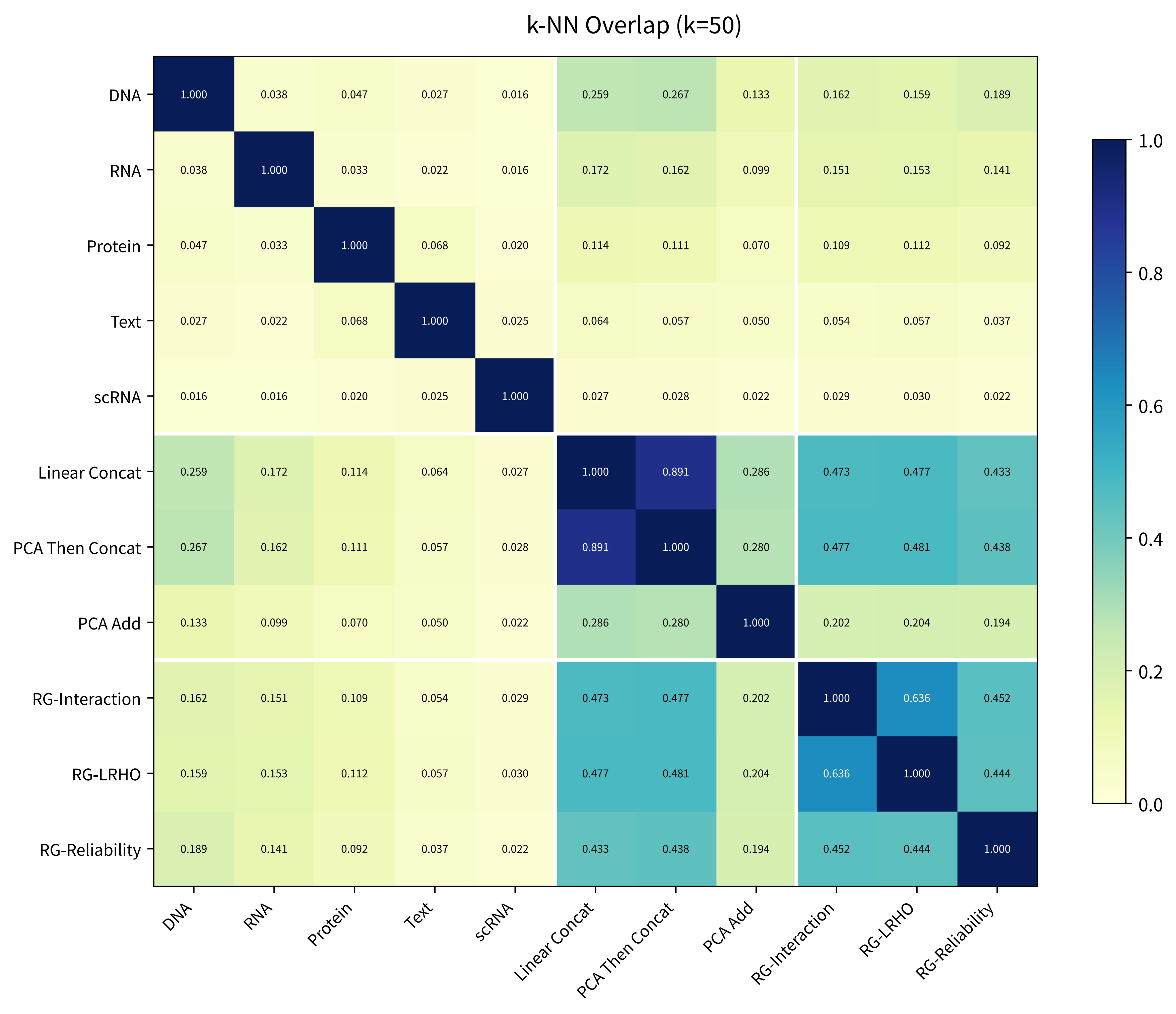

### overview_protocol.pdf

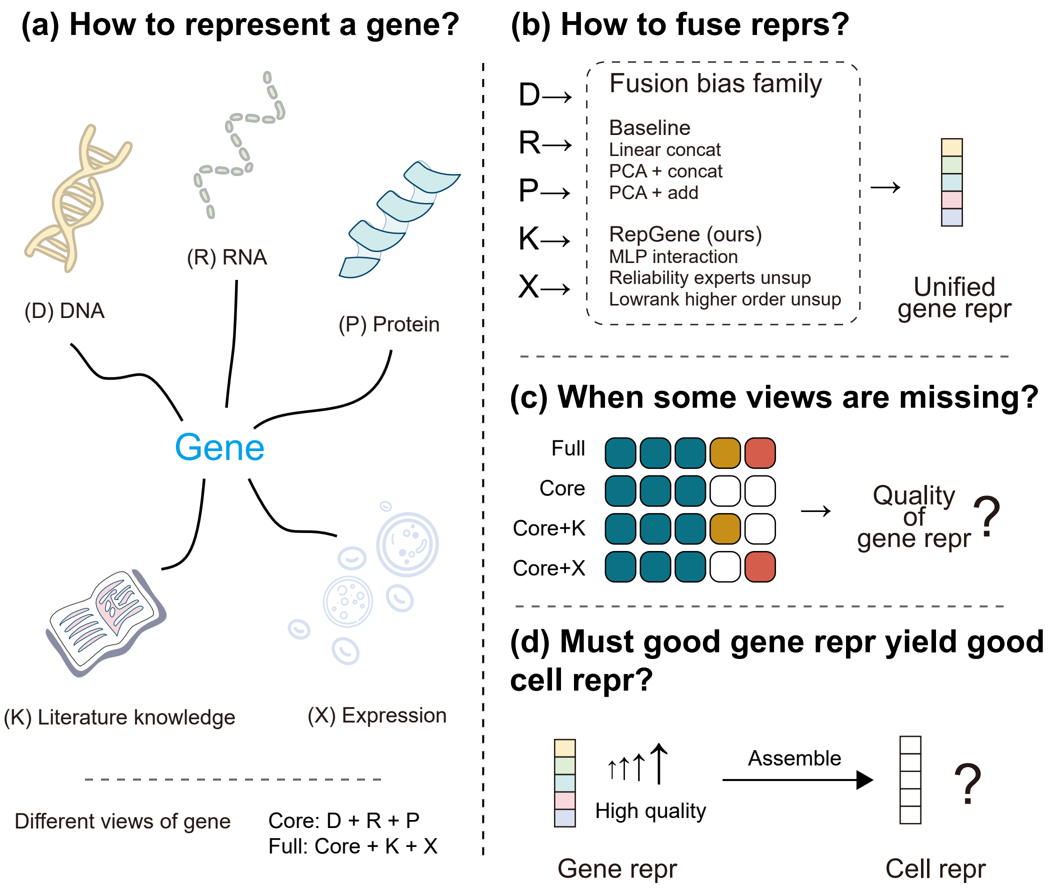
